# Creating a new oilseed crop, pennycress, by combining key domestication traits using CRISPR genome editing

**DOI:** 10.1101/2025.03.15.643467

**Authors:** Barsanti Gautam, Brice A. Jarvis, Maliheh Esfahanian, Michaela McGinn, Dalton Williams, Shengjun Liu, Mary E. Phippen, Nicholas J. Heller, Tad L. Wesley, Winthrop B. Phippen, Tim Ulmasov, M. David Marks, Ratan Chopra, John C. Sedbrook

## Abstract

Considerable offseason farmland lays fallow because there are few crops that can profitably fit between primary crops. To remedy, we employed CRISPR genome editing to the freeze-tolerant, rapid cycling wild Brassica, *Thlaspi arvense* L. (field pennycress). High-yielding domesticated pennycress varieties were created having seed compositions comparable to “double low” canola (low erucic acid and glucosinolate). Seed glucosinolate content was reduced 75 % by combining mutations in *R2R3-MYB (MYB28)* and bHLH *MYC (MYC3)* transcription factors. Pennycress weediness was greatly reduced by knockout of the bHLH transcription factor *TRANSPARENT TESTA8* (*TT8*), lowering seed dormancy and seed coat protections thereby mitigating pennycress re-emergence in fields. Domesticated pennycress offers farmers a profitable, low-carbon-intensity intermediate crop that confers ecosystem benefits while producing grain for renewable fuels and enhanced food security.

## INTRODUCTION

Row crops such as corn (*Zea mays*), soybean (*Glycine max*), and wheat (*Triticum aestivum*) are prevalent on farmland in temperate regions of the world, providing feed, food, fiber, and feedstock for biofuels. During the offseason, much of this farmland lays fallow or is planted in cover crops or intermediate crops. Cover crops (e.g. winter cereal rye (*Secale cereale*), annual ryegrass (*Lolium multiflorum*), radish (*Raphanus sativus*), clover (*Trifolium repens*, *T. pratense*), and hairy vetch (*Vicia hirsuta*)) are defined in the United States (US) as an offseason crop that cannot be harvested as a commodity but instead is typically terminated with herbicide and/or tilled before planting the next main-season crop. Cover crops provide many ecosystem benefits including enhanced soil health, improved nutrient cycling, reduced soil erosion, weeds suppression, shelter and provisions for insects and animals. Cover crops also result in reductions in greenhouse gas (GHG) emissions by sequestering carbon in soil and encouraging no-till farming practices (*1–4*). Intermediate crops (e.g. winter wheat, hybrid rye, winter canola, carinata, camelina, winter peas, oats, and clover) also confer cover crop benefits and, in addition, produce grain, forage, and fiber harvested as a commodity which incentivizes their use, allowing agricultural intensification without cropland expansion (*5–7*).

Year-round ground cover is considered essential to making annual cropping systems sustainable and meeting reduced GHG emissions goals (*8*, *9*), yet only seven percent of the 100 million hectares of U.S. Midwest farmland is planted in a crop in the offseason (*10*).This low adoption is largely due to cover crops being unprofitable even with government incentives and carbon market credits (*11*, *12*). Intermediate crops represent only a small fraction of that cover because most do not have short enough life cycles to fit between main-season crops and/or do not have the necessary freeze tolerance to survive temperate region winters (*13–17*).

Field pennycress (*Thlaspi arvense* L.) is a member of the Brassicaceae family related to Arabidopsis (*Arabidopsis thaliana*), rapeseed/canola (*Brassica napus* L. and *B. rapa* L.), carinata (*Brassica carinata* L.), camelina (*Camelina sativa* L.), and field cress (*Lepidium campestre* L.). Field pennycress is a ruderal species with notable freeze tolerance, native to Eurasia, naturalized to North America, and widely adapted to temperate regions on all continents except Antarctica (*18*, *19*). Both spring- and winter-type field pennycress populations naturally exist, with winter-types requiring vernalization to flower, typically germinating in the fall and overwintering as a rosette of leaves close to the ground before flowering in early spring (Fig. 1A). Pennycress reaches maturity by mid-May to early June in the lower US Midwest. These maturation dates are early enough for its grain to be harvested followed by planting of a full-season summer crop such as corn or soybean (Fig. 1B). Field pennycress produces copious amounts of seeds rich in oil and protein (∼33% and ∼23% dry weight, respectively; Fig. 1C, D), with wild accessions able to achieve 2,500 kg/hectare (2,200 lb/acre) grain yield before any breeding or genetic improvement.

**Fig. 1.**
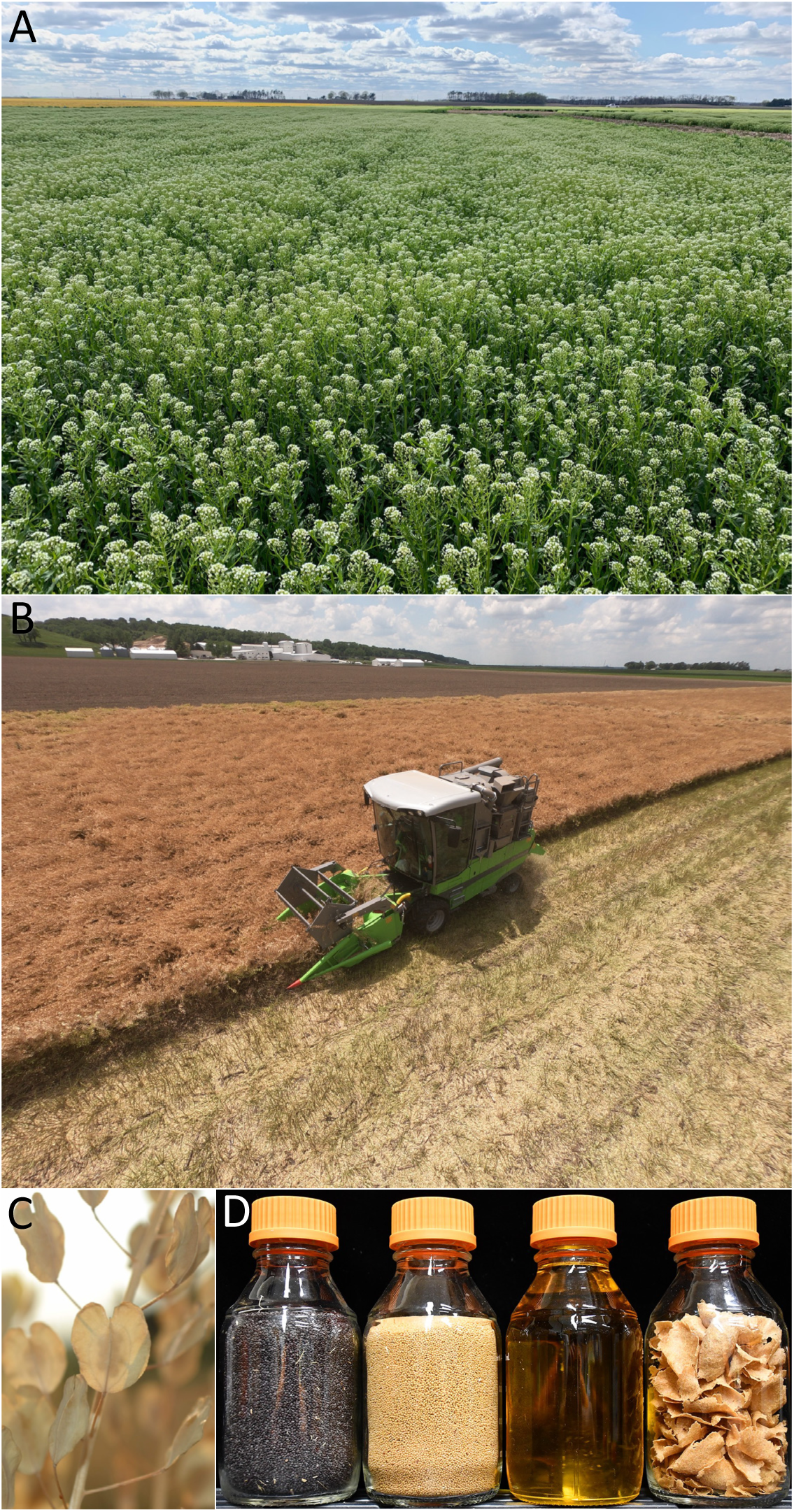
**(A)** Field of domesticated pennycress flowering near Arenzville, Illinois, USA (April 15th). Shown is variety CoverCress^TM^ Whole Grain which contains the low erucic acid and low seed coat fiber traits. **(B)** Harvest of CoverCress^TM^ Whole Grain (June 5th). **(C)** Senesced plants containing the *tt8* low fiber trait. Note the golden-colored seeds are visible in the silicles. **(D)** Left to right in 500 mL bottles: Wild-type Spring 32-10 seeds, *tt8 fae1* seeds, *tt8 fae1* oil and meal which were extracted with a screw press.

In the early 2000’s, researchers at the United States Department of Agriculture (USDA) proposed that field pennycress could serve as a winter annual oilseed crop fitting within corn-soybean rotations, producing oil suitable for biodiesel production (*20–25*). However, the potential weediness of field pennycress in some cropping systems was a concern (*19*, *26*), and compositional improvements to the seed oil and meal was necessary for optimal biofuels production and for the seed meal to be palatable at feed inclusion rates greater than 5 percent as was suggested in a study of broiler chickens (*27*).

Pennycress seed oil is in the form of triacylglycerides (TAG) with about 35 percent of the fatty acid content being erucic acid (C22:1). This very long chain fatty acid is deemed cytotoxic (based on limited data from animal feeding studies) and is therefore regulated in feed and foodstuffs (*28–30*). Pennycress seeds also contain over 100 µmol/gram sinigrin, which is an aliphatic glucosinolate converted by myrosinase to toxic isothiocyanate and nitrile compounds upon tissue damage, functioning in part as a defense against herbivory (*31–33*). While studies suggest sinigrin has therapeutic benefits (*34*), because of the toxicity of glucosinolate breakdown products and their unpalatability at high levels (sinigrin and its breakdown products are what makes horseradish taste so strong), the Association of American Feed Control Officials (AAFCO) has established a limit of 30 µmol/gram glucosinolates in Brassica seed meal. Unlike most other Brassicas which produce many different glucosinolates (*35–38*), field pennycress almost exclusively produces sinigrin, which is derived from the amino acid, methionine (*32*, *39*).

Erucic acid and glucosinolates are also prominent in the Brassica oilseed crop, rapeseed, which has been cultivated for thousands of years (*40*). In the 1960s and ‘70s, prompted by concerns of the edibility of rapeseed, breeders identified lines having substantially reduced seed oil erucic acid content and reduced seed glucosinolate content, combined those traits through breeding, and successfully overcame related yield deficiencies to create “double low” varieties trademarked as canola (*41–44*). Canola is now planted on an estimated 40 million hectares of farmland annually, yielding over 80 million metric tons of grain globally.

While glucosinolate pathways have been extensively studied in various Brassicaceae (*45–48*), important knowledge gaps remain with respect to what mutations or mutant combinations reduce glucosinolate content without negatively impacting plant growth and seed yields. Even though low glucosinolate content is a hallmark of canola, it remains unclear what is the genetic basis of this trait due to the complexity of its polyploid genome (*49*).

In 2012, with the creation of canola as an inspiration and basic research findings in Arabidopsis as a guide, we set out to domesticate field pennycress into the crop, domesticated pennycress (pennycress), using conventional mutagenesis methods and by developing an *Agrobacterium*-mediated floral dip transformation method allowing for targeted mutagenesis (*39*, *50–53*). Here, we report the use of clustered regularly interspaced short palindromic repeats (CRISPR) genome editing to identify pennycress mutations suitable for its domestication. We efficiently combined those mutations into lines with canola-like seed compositions that grow and yield similar to wild type pennycress. Similar mutations have been introduced into pennycress breeding lines to generate commercial varieties trademarked as CoverCress^®^. CoverCress^®^ and other domesticated pennycress varieties are poised to be planted on millions of hectares of farmland globally as an intermediate crop to provide feedstock for biodiesel, renewable diesel, and sustainable aviation fuel (SAF) production. Domesticated pennycress also produces nutritious seed meal well-suited for climate-smart animal production, enabling and incentivizing sustainable agricultural practices.

## RESULTS AND DISCUSSION

### Generating “double low” pennycress: Improving seed oil composition

We set out to domesticate field pennycress by identifying and combining mutations improving seed compositions and removing weediness traits without sacrificing plant growth and seed yields. We used Near Infrared Spectroscopic analysis (NIRS) and wet chemistry methods to screen the seeds of over 800 wild field pennycress populations collected from around the world for the low erucic acid and low glucosinolate seed traits which are hallmarks of “double low” canola. We found no accessions with low enough levels to suggest pennycress double low could be achieved through cross pollination-based breeding alone (*51*, *59*, unpublished). Therefore, to remove from pennycress seed oil the very long chain fatty acid, erucic acid (C22:1), which is considered antinutritional, we previously reported the use of CRISPR genome editing to generate insertion/deletion (indel) mutations in the *FAE1* gene thereby reducing erucic acid content to zero (*51*; Fig. 2A). Furthermore, to improve the oxidative stability of pennycress oil, we identified CRISPR-induced knockout mutations in the *ROD1* gene which, when stacked with *fae1*, reduced the polyunsaturated fatty acids linoleic (18:2) and linolenic (18:3) content to 16% and 17%, respectively, and increased oleic acid (18:1) to 62% of TAG, resulting in a fatty acid profile similar to that of canola oil (Fig. 2A) (*39*, *53*). *rod1 fae1* double knockout mutant plants grew and had seed yields similar to wild type. *rod1* knockout did not affect total seed oil content; however, *fae1* knockout resulted in about 10% less oil per seed compared to wild type on a seed dry weight basis (Fig. 2B). This *fae1*-related reduction was likely due to the shorter fatty acid chain lengths resulting in reduced TAG mass along with changes in metabolic flux (*66*).

**Fig. 2.**
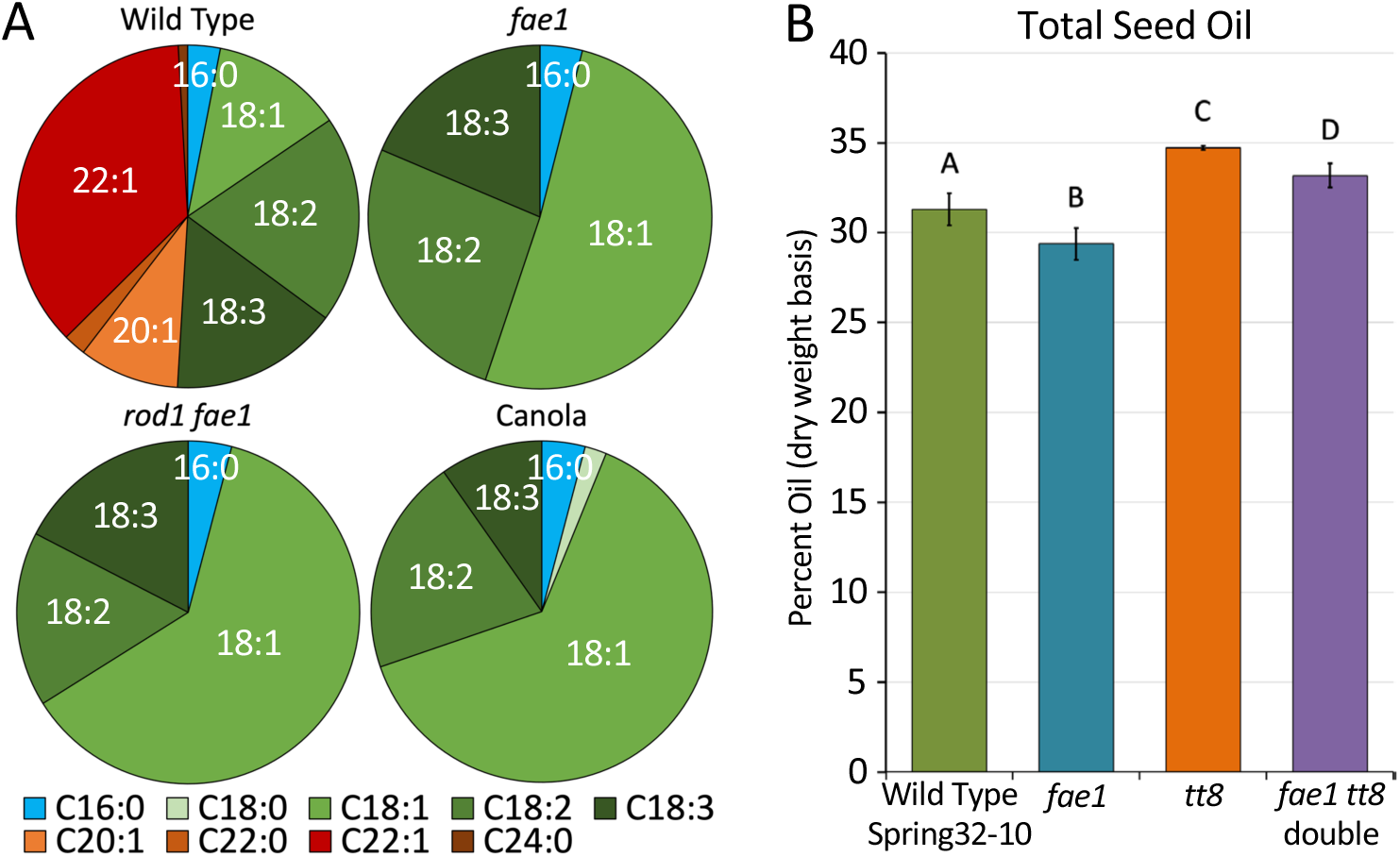
Seed oil fatty acid (FA) content in wild-type pennycress versus representative *fae1* and *rod1 fae1* knockout mutants, compared to canola. Note that *fae1* knockout abolishes the very long chain fatty acids (22:1 erucic acid, 20:1 eicosenoic acid, and 20:0 arachidic acid). Combining *rod1* and *fae1* knockout mutations decreased the oxidatively unstable polyunsaturated fatty acids (18:2 linoleic acid and 18:3 linolenic acid) and increased the more stable 18:1 oleic acid. (**B**) Pennycress total seed oil content (percent dry weight basis). Note that *fae1* (−4bp) and *tt8* (+A) knockout mutations significantly reduced and increased seed oil content, respectively, compared to wild type. Combining *fae1* and *tt8* knockout mutations had an additive effect. Columns represent averages, error bars represent standard deviations, and different letters represent significant differences using one-way ANOVA and Tukey’s test. 21≤n≤24.

### Generating “double low” pennycress: Reducing seed glucosinolate content

Field pennycress is commonly known as “stinkweed” owing to the plant’s pungent odor from the sulfur-containing secondary metabolite, glucosinolate, and corresponding isothiocyanate breakdown products (*32*, *67*, *68*). Glucosinolates have been shown to be synthesized in vegetative tissues and transported to developing seeds (*69–71*). In the pennycress wild-type Spring32-10 inbred line, we determined that glucosinolate accumulated to around 120 µmol/gram in mature seeds and 20 µmol/gram in cauline leaves (Fig. 3A, B).

**Fig. 3.**
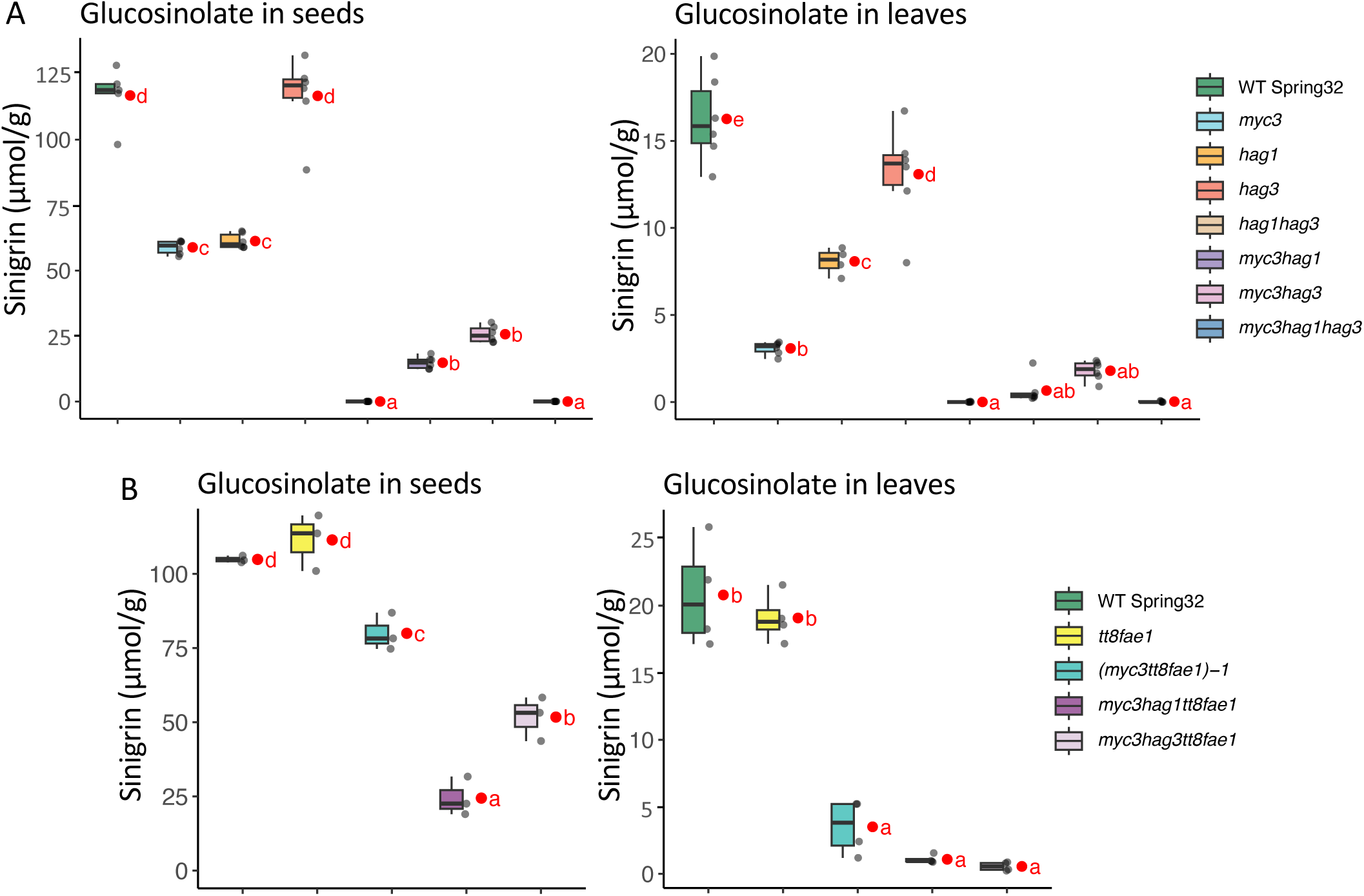
(**A, B**) Box plot graphs of glucosinolate (sinigrin) content in mature seeds (left two graphs) and in cauline leaves from flowering plants (right two graphs) of the various glucosinolate mutants without (**A**) or when combined with the *tt8* and *fae1* mutations (**B**), versus the wild-type (WT) Spring 32-10 control. Grey dots represent each sample data and red dots estimated marginal means. Means not sharing any letter are significantly different by the Sidak test at the 5% level of significance. n=6. Mutations were as follows (in parentheses; also see Supplemental Sequences file): *myc3* (+A); *hag1* (−4bp); *hag3* (−A); *hag1 hag3* (−4bp/−A); *myc3 hag1* (+A/−4bp); *myc3 hag3* (+A/−A); *myc3 hag1 hag3* (+A/−4bp/−A); *tt8 fae1* (+A/−4bp); (*myc3 tt8 fae1*)−1 (−17bp, −144bp /+A/−4bp); *myc3 hag1 tt8 fae1* (−17bp, −144bp/−4bp/+A/−4bp); *myc3 hag3 tt8 fae1* (−17bp, −144bp/595bp inversion/+A/−4bp).

In Arabidopsis, it was shown that mutations in the *MYB28*/*HIGH ALIPHATIC GLUCOSINOLATE 1* (*HAG1*) and *MYB29*/*HAG3* transcription factors reduced seed glucosinolate content by as much as 50 % (*72–74*). Moreover, genome wide association studies (GWAS) in *Brassica napus* found that *HAG1* and *HAG3* loci mapped to chromosomal intervals associated with reduced glucosinolate content (*49*, *75*, *76*). Therefore, we prioritized generating mutations in the putative pennycress *HAG1* (TAV2_LOCUS5186; Genbank CAH2044027), *HAG2* (TAV2_LOCUS20927; Genbank CAH2073631), and *HAG3* (TAV2_LOCUS20427; Genbank CAH2073629) genes which are predicted to encode proteins sharing 84%, 74%, and 79% amino acid sequence identities with the corresponding Arabidopsis proteins.

We generated multiple CRISPR-induced mutant lines containing insertion/deletion (indel) mutations that likely knocked out gene function, as most were frameshift mutations or deleted crucial portions of the genes’ coding sequences (data S1). In a representative experiment, wet chemistry and non-destructive Near Infrared Spectroscopic (NIRs) analyses revealed seed glucosinolate content in *hag1* indel mutants was 60 µmol/gm which is about a 50 percent reduction relative to wild-type controls (Fig. 3A and fig. S1). In contrast, *hag2* indel mutations resulted in no discernible glucosinolate reductions, and *hag3* indel mutations conferred slight glucosinolate reductions that were not always significant (Fig. 3A and fig. S1). All *hag1*, *hag2*, and *hag3* single mutant lines grew and had seed yields similar to wild type (fig. s2 to s6). While the amounts of glucosinolate varied from grow out to grow out, perhaps due to environmental effects as glucosinolate content is known to be induced by abiotic and biotic stressors (*46*, *47*, *77*, *78*), the relative differences between the mutant lines remained the same.

We combined *hag1*, *hag2*, and *hag3* mutations to make double and triple mutant lines, identifying a compositionally desirable 80 to 100 percent reduction in *hag1 hag3* double mutants’ seed glucosinolate content relative to wild type, to below 20 µmol/gm (Fig. 3A and fig. S1). However, *hag1 hag3* double mutant plants and *hag1 hag2 hag3* triple mutant plants exhibited relatively slower germination (fig S3) and at times displayed yellowish floral organs and reduced seed yields (fig. S2B and S2C), putting into question the suitability of these knockout mutant combinations for commercial use.

Previous studies implicated a family of three closely related basic helix-loop-helix (bHLH) transcription factors in regulating glucosinolate production: *MYC2*, *MYC3*, and *MYC4* (*33*, *79*). In Arabidopsis, *myc2 myc3 myc4* triple mutants produced seeds having near-zero glucosinolates content whereas single mutants showed no reductions (*33*). We performed a forward genetic screen of our ∼900-line pennycress mutant gene index collection of whole genome sequenced EMS and fast neutron mutagenized lines, employing our NIRS assay. This screen identified two lines having reduced seed glucosinolate content and predicted loss-of-function mutations in the putative *MYC3* gene. Pennycress *MYC3* (TAV2_LOCUS6314; Genbank CAH2046167) encodes a predicted 66.5 kD protein sharing 81% amino acid sequence identity with Arabidopsis MYC3 (AT5G46760). An EMS mutant line named D3_N13P3 contained a nonsense mutation (G to A transition) at nucleotide 198 of the 1,818 nucleotide-long *MYC3* coding sequence whereas a fast neutron mutant line named E3_202P2 had a single base deletion (−A) at nucleotide 256 resulting in a frameshift mutation. Wet chemistry analyses confirmed that seeds from these lines contained 66 µmol/gm and 47.4 µmol/gm glucosinolate, respectively, representing 45 percent and 59 percent reductions compared to 119 µmol/gm in wild-type control seeds.

It was unexpected to identify such large glucosinolate reductions in pennycress *myc3* mutants given that no glucosinolate reductions were observed in Arabidopsis *myc3* single mutants (*33*). To confirm the pennycress EMS mutants reduced glucosinolate phenotype was due solely to loss of *myc3* function and not second-site mutations, we employed CRISPR genome editing to precisely target indel mutations in *myc3*, identifying multiple independent mutant alleles (data S1). In a representative experiment, wet chemistry analyses revealed seed glucosinolate content of 59 µmol/gram in a *myc3* CRISPR-induced mutant containing a single nucleotide insertion (+A) in the *MYC3* coding sequence, compared to 123 µmol/gram in wild-type seeds (Fig. 3A). *myc3*(+A) mutant cauline leaf glucosinolate content was 3 µmol/gram compared to 16 µmol/gram in wild type grown alongside (Fig. 3A). These reduced glucosinolate levels were also observed in other *myc3* knockout mutant lines we generated (fig. S7) and were consistent with the reduced levels observed in the EMS-generated *myc3* mutant lines. All *myc3* mutant lines germinated, grew and had seed yields similar to wild type (fig. S2 to S8).

Since the *myc3*, *hag1*, and *hag3* single knockout mutations did not reduce seed glucosinolate content to the preferred 30 µmol/gm target, we generated combinatorial mutants through both cross-pollination and by employing CRISPR multiplexing to introduce simultaneous edits, to determine if seed glucosinolate content could be reduced further while maintaining plant health. In a representative experiment, we found that CRISPR-generated *myc3 hag1* double knockout mutants contained, on average, 15 µmol/gm seed glucosinolate content, and *myc3 hag3* double knockout mutant seeds contained 28 µmol/gm seed glucosinolate, representing desirable 88% and 79% reductions compared to wild-type controls (Fig. 3A). Cauline leaf glucosinolate content was 1 µmol/gm, 2 µmol/gm, and 16 µmol/gm for *myc3 hag1*, *myc3 hag3*, and wild type, respectively (Fig. 3A). *myc3 hag1* and *myc3 hag3* double mutant plants developmentally were comparable to wild type and had statistically indistinguishable seed yields in both growth chamber and field conditions (Fig. 4; fig. S2 to S8).

**Fig. 4.**
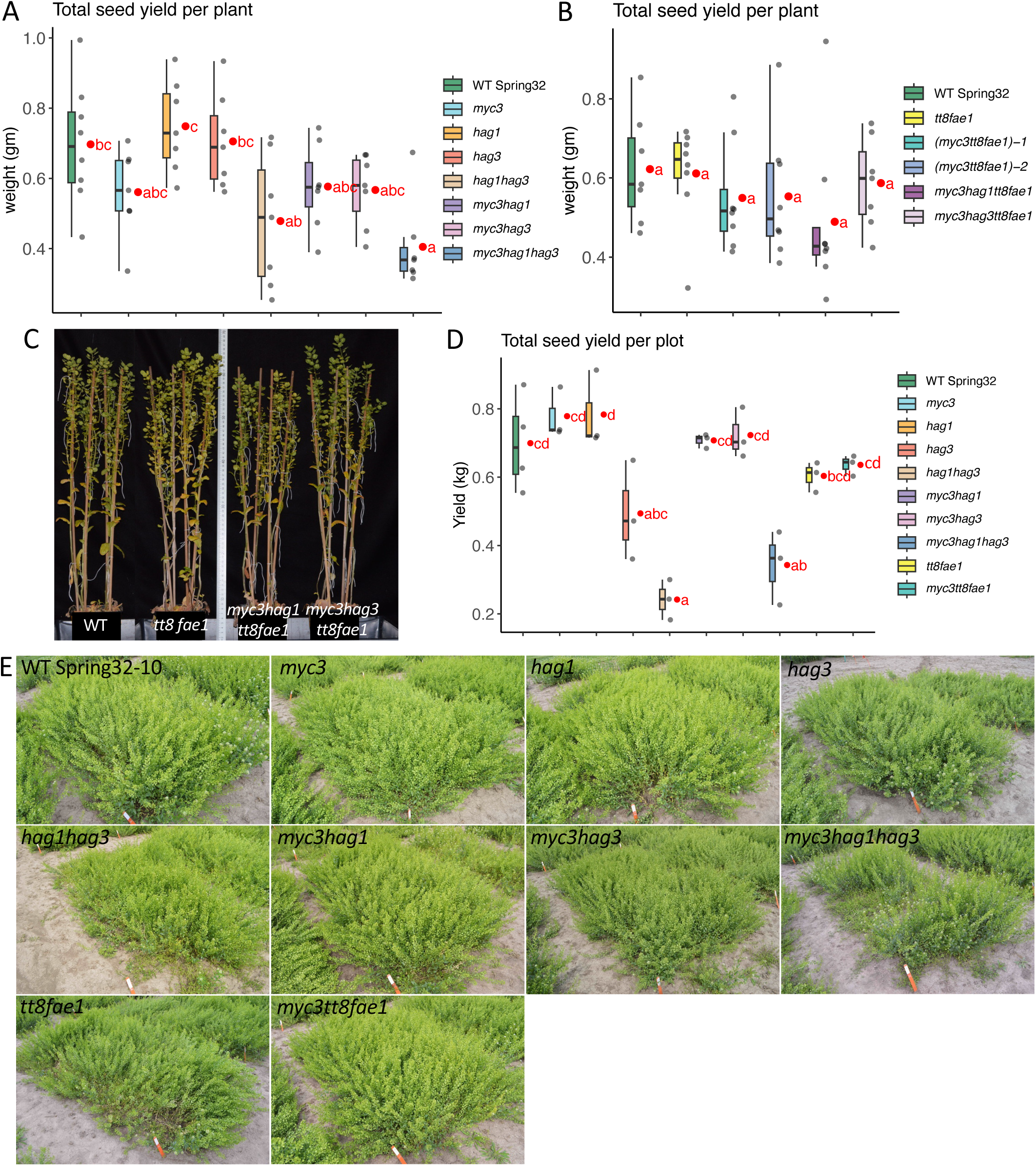
(**A, B**) Box plots of seed yield per plant of growth chamber-grown glucosinolate mutants versus wild-type (WT) and *tt8 fae1* controls. (**C**) Representative pictures of growth chamber-grown plants (four plants per 4-inch pot). (**D**) Seed yield per 5’x10’ plot of field-grown mutants versus wild-type (WT) and *tt8 fae1* controls. Note the average yield of the *hag3* plots was abnormally low due to the use of an older seed lot which did not germinate as well. (**E**) Field plots (5’x10’ in size) of the glucosinolate single and multiple mutant combinations grown in a common garden along with WT and *tt8 fae1* controls (plots were grown in triplicate; yield data in (**D**)). All lines grew comparably well to the controls except for lines containing *hag1 hag3* double knockout. For graphs, grey dots are each sample data, red dots estimated marginal means.Means not sharing any letter are significantly different by the Sidak test at the 5% level of significance. See Figure 3 legend for mutations information.

Notably, both chamber grown and field grown seeds of the *myc3* single mutants and *myc3 hag1* and *myc3 hag3* double mutants were, on average, significantly larger than wild type (10 to 25 % larger based on area measurements) and weighed significantly more on a per seed basis (20 to 40 % heavier than wild type; fig. S9). Moreover, *myc3*, *myc3 hag1*, and *myc3 hag3* seeds tended to have more oil and protein compared to wild type (fig. S10 and S11). The pennycress *myc3* larger seed findings were like findings in Arabidopsis where *myc2 myc3 myc4* mutant combinations produced larger seeds with more protein (*80*), and *myb76*/*hag2* mutant seeds were found to have increased amounts of total fatty acids (*81*). Taken together, these data suggest mutations in glucosinolate-related *MYC* and *MYB* family members result in tradeoffs between this secondary metabolite production and agronomically desirable seed size and compositional traits.

To gain insights into how the single *myb* and *myc* knockout mutations and mutation combinations affected expression of genes involved in glucosinolate biosynthesis and other processes, we purified RNA from cauline leaf tissue and performed RNAseq analysis, identifying overlapping yet distinct gene expression differences. Transcriptomic analyses revealed that knockout of *MYC3* function generally reduced expression of glucosinolate biosynthetic genes considerably more than *HAG1* or *HAG3* single knockout, suggesting functional redundancy between *HAG1* and *HAG3* but limited functional redundancy between *MYC3* and other glucosinolate-related transcription factors in pennycress. Consistent with this, double knockout of *HAG1* and *HAG3* resulted in drastic expression reductions of glucosinolate biosynthetic genes (Fig. 5, fig. S14). For example, the expression of four genes involved in the methionine side-chain extension portion of aliphatic glucosinolate biosynthesis (*ISOPROPYLMALATE ISOMERASE* (*IPMI*), *ISOPROPYLMALATE DEHYDROGENASE* (*IPMDH1*), *BRANCHED-CHAIN AMINO ACID TRANSFERASE4* (*BCAT4*), and *METHYLTHIOALKYLMALATE SYNTHASE* (*MAM1*)) were reduced 86 to 95 % in *myc3* versus 32 to 47 % in *hag1* and 38 to 57 % in *hag3*. In the double mutants, expression of these three genes were reduced by 98 to 99 % in *myc3 hag1*, 96 to 99 % in *myc3 hag3*, and 98 to 100 % in *hag1 hag3* (fig. S14). These relative expression reductions mostly mirrored the relative reductions in cauline leaf glucosinolate content (80% reduction in *myc3*, 48% in *hag1*, 16% in *hag3*, 96% in *myc3 hag1*, 88% in *myc3 hag3*, 100% in *hag1 hag3*; Fig. 3).

**Fig. 5.**
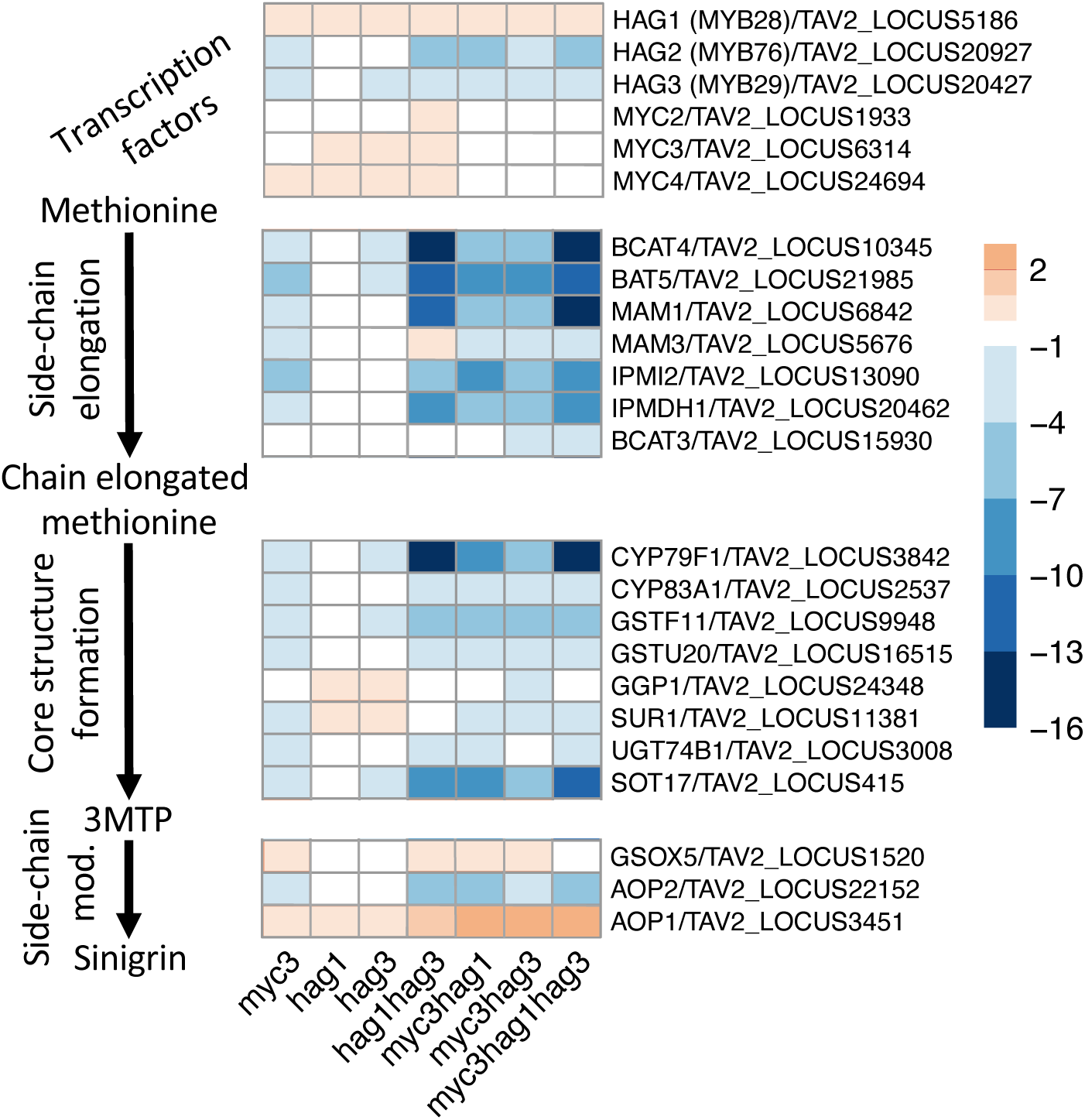
Transcript level differences of genes encoding transcription factors and biosynthetic enzymes involved in glucosinolate biosynthesis in the various glucosinolate single and combinatorial mutants versus wild type. Heat map shows log2 fold changes. The sinigrin biosynthetic pathway is depicted to the left, with the corresponding gene names/I.D.s to the right in the order they function in the pathway. Mutant names are listed below. Positive and negative values represent expression increases and decreases relative to wild type, respectively. See Supplemental Figure 14 and Supplemental RNAseq Data file for corresponding transcripts counts data.

Gene ontology analysis of the RNAseq data also revealed that *myc3*, *hag1*, and *hag3* loss of function, singly and in combinations, impacted the expression of genes involved in many different biological processes, in complex ways (fig. S15 to S17). For example, knockout of *myc3* significantly downregulated genes linked to the biosynthesis of aliphatic, indole, and sulfur amino acid metabolic processes, whereas neither *hag1* nor *hag3* single knockouts downregulated these genes’ expressions, but *hag1 hag3* double knockout did (fig. S15). *hag3* knockout was linked to upregulation of genes linked to hypoxia and decreased oxygen cellular responses whereas neither *myc3* nor *hag1* knockout significantly affected expression of genes linked to these biological processes. These knockout mutations, singly and in combinations, also altered the expression of many genes partaking in jasmonic acid and salicylic acid responses (fig S16 and S17) that in Arabidopsis are linked to resistance or susceptibility to pathogens such as *Pseudomonas syringae*, *Erwinia carotovora*, and *Alternaria brassicicola* (AT5G46050, AT5G01900, AT5G24530, AT2G17120, AT2G13810, *82*–*86*) and pests such as spider mites (AT1G73260, AT3G07195, *87*, *88*). While these pennycress *myc3*, *hag1*, and *hag3* single mutants and mutant combinations (except for *hag1 hag3* double mutants) generally grew well in farm fields from fall through spring (Fig. 4D and 4E), studies should be done to determine if pathogen and/or pest resistance is altered or if growth is affected under certain environmental conditions.

### Pennycress transparent testa8 (tt8) knockout enhances germination, improves seed compositions, and removes weediness traits

Field pennycress seeds have relatively thick seed coats which are dark brown in color due to condensed tannins which form upon oxidative polymerization of proanthocyanidins (Fig. 6A, *50*). Condensed tannins are commonly found in Brassica seeds and are implicated in deterring pests and pathogens and contributing to seed dormancy thereby maintaining viable seeds in soil for many years (*89–91*). Field pennycress plants can each produce thousands of seeds, with only a small portion germinating each season when conditions are favorable, particularly after the soil is disturbed (*18*). These are features of ruderal species and make field pennycress a weedy concern (*19*). Besides enabling field pennycress weediness, the thick resilient seed coat has relatively high indigestible fiber content (∼25% Acid Detergent Fiber (ADF)) which detracts from the nutritional value of the grain as well as the meal remaining after oil is pressed (Table 1, *27*).

**Fig. 6.**
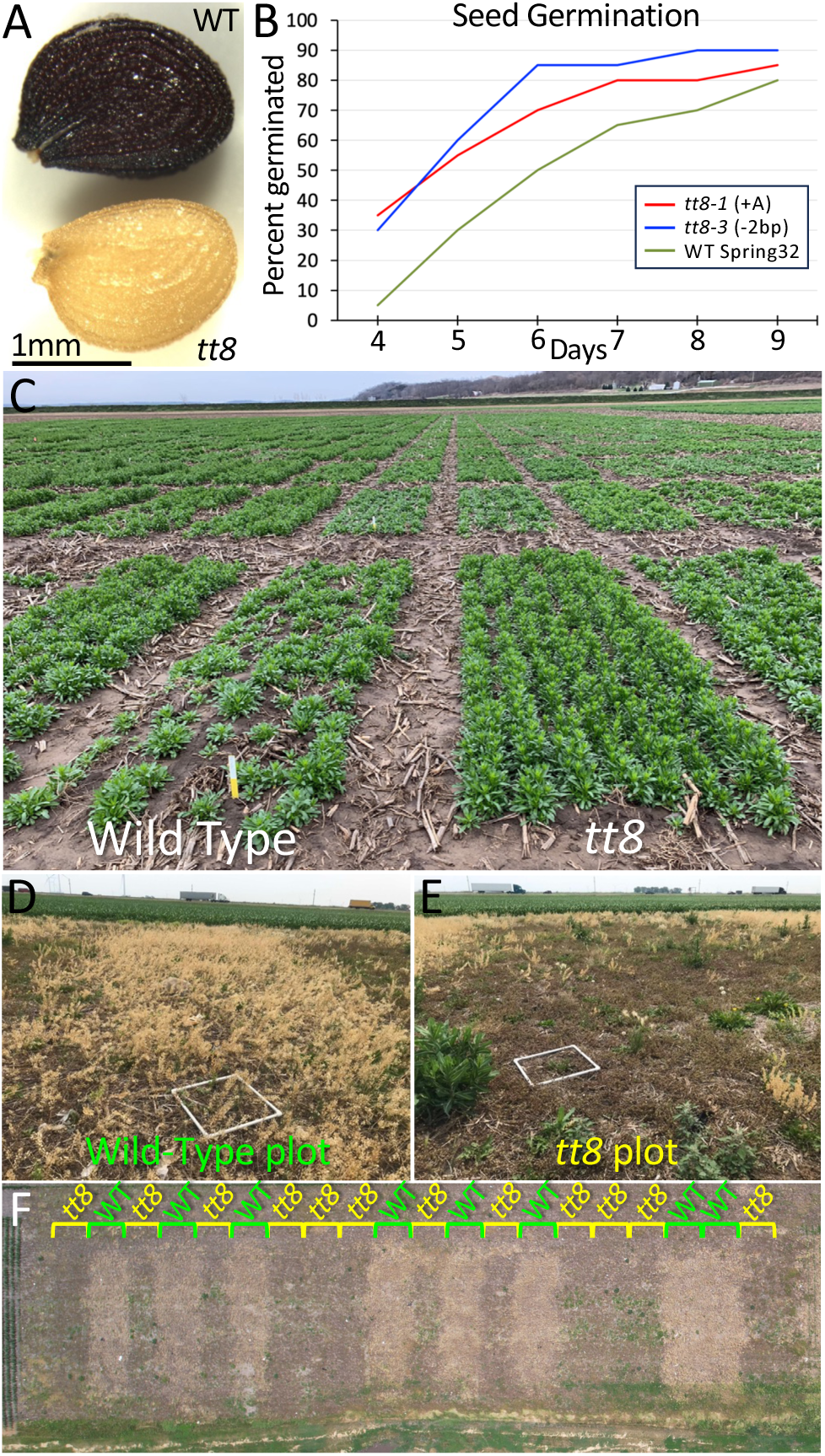
(**A**) A pennycress wild-type seed (top) and a *tt8* mutant seed (bottom). (**B**) Seed germination rates of two *tt8* knockout mutants versus wild type sowed on the surface of wet potting soil (n=100). (**C**) CoverCress, Inc. breeding plots near Arenzville, Illinois USA. The front left and front right plots are a breeding line without (Wild Type) and with a CRISPR-introduced *tt8* knockout mutation. Note the *tt8* line germinated and established markedly better. (**D-F**) *tt8* mutant and wild-type replicated plots, one year after the originally planted plots were harvested and then left undisturbed. Note the considerable re-emergence of the wild-type plants (**D, F**). No *tt8* plants were found in any of the plots due to the seeds rapidly losing viability in field settings (low persistence).

**Table 1.**
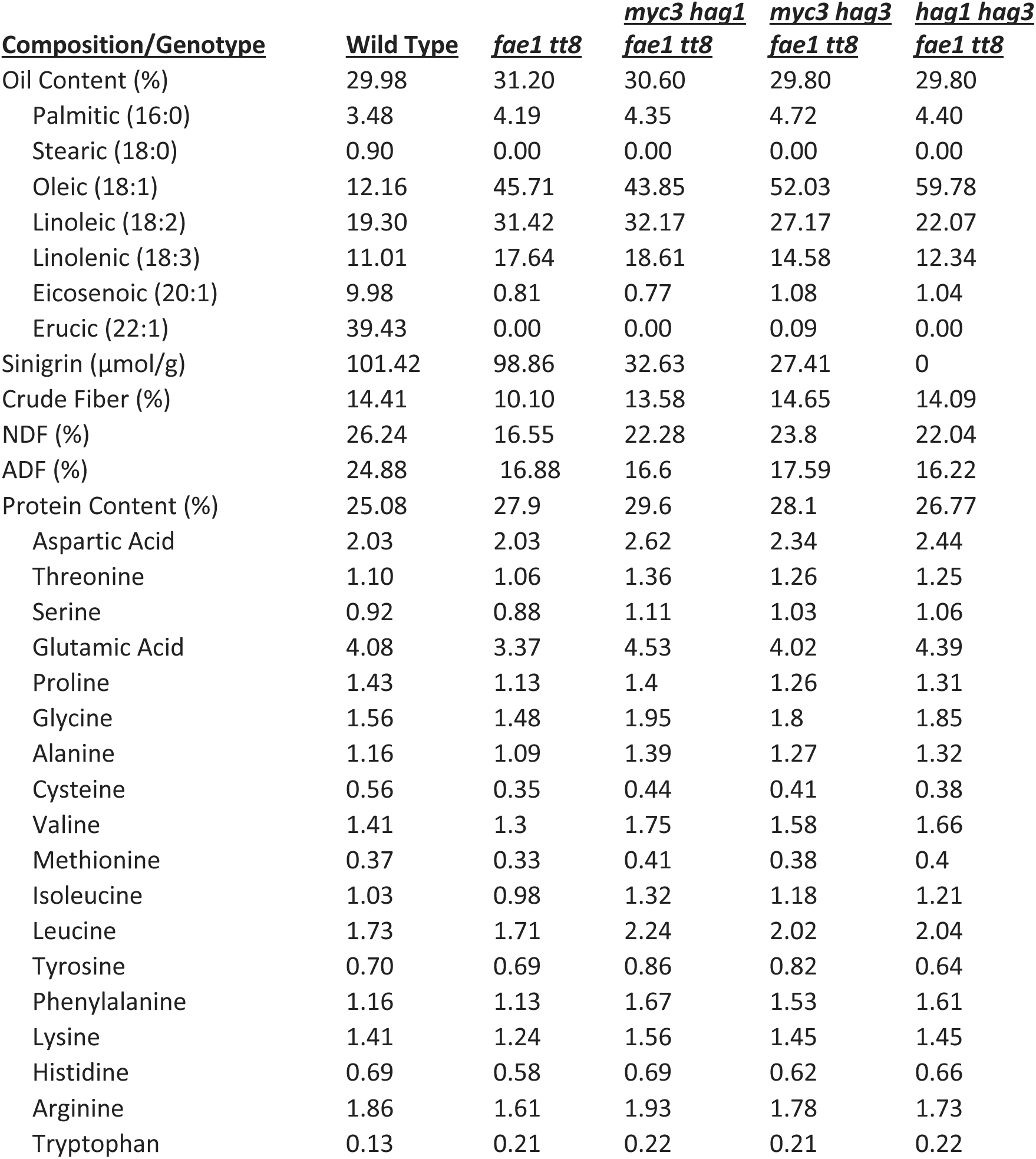
Seed compositions of pennycress lines containing mutations conferring the core domestication traits. Percentages are on a seed dry weight basis. The values for fatty acids (names indented below Oil Content) and amino acids (names indented below Protein Content) are a percent of the total fatty acids and protein, respectively.

We previously reported identifying field pennycress EMS mutant lines containing loss-of-function mutations in many of the biosynthetic and regulatory genes involved in proanthocyanidin biosynthesis and condensed tannins formation (*50*, *92*). These are collectively known as *TRANSPARENT TESTA* (*TT*) genes, given that the seeds of Arabidopsis loss-of-function mutants were identified as having somewhat transparent seed coats (testa) and tan to golden colors due to reduced or absent condensed tannins (*93*). Phenotypic analyses of these pennycress EMS mutants revealed that many grew similar to wild type, although unexpected growth phenotypes were observed in some of the EMS mutant lines possibly due to second site mutations. To systematically generate *tt* knockout mutants with no second site mutations, we employed CRISPR genome editing, choosing protospacers containing nucleotide sequences which perfectly matched only the target within each *TRANSPARENT TESTA* gene. Given that knockout mutations in the putative pennycress transcription factors *TT2*, *TT8*, *TTG1*, and *TT16* had the lightest seed coat colors (“golden/yellow”), we focused attention on those, in particular *TT8* given its reported role in Arabidopsis of negatively regulating fatty acid biosynthesis (*94*).

We found that CRISPR-induced pennycress knockout mutations in *TT8* produced a variety of agronomic improvements without impacting plant growth and seed yields. *tt8* mutant seeds germinated faster and more consistently than wild-type which, in the field, translated to better stand establishment (Fig. 6B and 6C). Moreover, *tt8* seeds had about 13% higher oil content, 11% higher protein content, and 24% lower crude fiber content compared to wild type, all of which are value-added traits (Fig. 2B, fig. S11, and Table 1). While two dimensional measurements suggested *tt8* seeds were smaller than wild type, per seed weight was not different from wild type (Fig. 6A and fig. S9B). Three-dimensional measurements revealed the seeds had the same volume as wild type but appeared smaller because they were more spherical instead of the flattened football shape of wild type (*tt8* seeds are designated “Golden” in Griffiths et al. (*95*)).

*tt8* mutant plants grew indistinguishable from wild type both singly and when *tt8* knockout mutations were stacked with other mutations conferring the core domestication traits, namely *fae1*, *myc3*, *hag1*, and *hag3* under both growth chamber and field conditions (Fig. 4 and 6, fig. S4 to fig. S8). Rai et al. (96) reported that Arabidopsis *tt8* loss of function led to increased sensitivity to multiple abiotic and biotic stressors. While systematic stress analyses of pennycress *tt8* mutants remain to be conducted, growth of *tt8* mutants in farm fields in six U.S. Midwestern states over multiple years, from fall to spring which exposed the germinating seedlings and plants to severe drought, cold, and heat, did not reveal any obvious growth differences compared to wild type.

Since *tt8* mutants had substantially reduced seed coat fiber, lacked condensed tannins, and germinated more readily than wild type, we assessed persistence of pennycress seeds in farm fields over multiple years to determine if lines carrying *tt8* mutations were less weedy. In one representative experiment, 20 replicated *tt8* and wild-type plots (ten 15’ by 50’ randomized plots of each) were established following corn or soybean in the fall of 2021, harvested in the spring of 2022, then the plots left undisturbed until June of 2023 allowing for volunteer plants to grow and mature. At maturity, the colors of the seeds in the pods were clearly visible and readily scorable. We found that all ten of the wild-type plots had thousands of volunteer plants with dark wild-type seeds whereas the ten *tt8* plots had relatively few pennycress plants (Fig. 6 D-F). Moreover, the plants in the *tt8* plots were all dark-seeded wild-type; no yellow-seeded *tt8* plants were found in any of the 20 plots.

Similar results were observed in two sets of field plots planted in the fall of 2020 and the fall of 2021 (fig. S18). In the 2020-2021 fall to spring field season, 84 5’ x 15’ plots were planted, 32 plots of which contained lines having *tt8* knockout mutations. In the 2021-2022 fall to spring field season, 84 5’ x 15’ plots were planted, with 28 plots containing lines having *tt8* knockout mutations and another two plots having plants with a *tt2* knockout mutation. All plots were harvested after reaching maturity and left undisturbed until June of 2023 at which time we looked for any plants with yellow seeds indicative of the *tt* mutants. No yellow-seeded plants were found in the locations where the 2020-2021 field season plots had been. Only six plants with yellow seeds were found amongst the tens of thousands of dark (wild-type) seeded plants (fig. S18C to S18E). Also of note, substantially more pennycress plants were present in the 2021-2022 plots location than in the 2020-2021 plots location (fig. S18C to S18E), signifying that field pennycress populations dwindle over time when facing competition from other weeds in soil left undisturbed. Taken together, these data demonstrate that seeds from pennycress *tt8* knockout plants that fall to the ground before or during harvest have very low persistence, likely due to greatly reduced seed coat protections.

## CONCLUSION

By employing CRISPR genome editing, we rapidly introduced and identified a core set of mutations improving seed oil compositions and substantially lowered seed glucosinolate content, creating “double low” pennycress varieties comparable to “double low” canola. These compositional changes make the pennycress oil and seed meal well-suited for biofuels and feed/food uses. Moreover, introduction of *tt8* loss of function improved seed germination and crop establishment in field settings and, importantly, greatly reduced pennycress weediness. While domesticated pennycress lines containing these combined traits grew and yielded similar to wild type both in growth chamber and field settings, transcriptome analyses revealed that the mutations in *MYB* and *MYC* transcription factors while reducing seed glucosinolate content also resulted in altered expression of many genes linked to hormonal and plant defense responses.

Work is underway to determine if abiotic and/or biotic stress responses of these mutants are negatively impacted under diverse field conditions. If so, given that these domesticated lines grow and yield similar to wild type, and given the successful creation of canola from rapeseed, it is likely that ongoing breeding efforts and targeted mutagenesis can overcome any genetic shortcomings that are identified. The domesticated pennycress described in this publication offers farmers a profitable, low-carbon-intensity offseason crop that delivers both ecosystem benefits and food security, while producing oil well-suited for easy conversion to renewable fuels.

## ACKNOWLEDGMENTS

We thank the numerous undergraduate students, graduate students, and technicians who helped with taking care of plants, seed and tissue measurements, field work, and other supporting work. We thank Manesh Shah, Chanaka Roshan Abeyratne, and Daniel Jacobson for generating the transcripts counts file and for training of B.G. at Oak Ridge National Laboratory, and Bill Perry for help with statistics and R coding. We thank Mason Tuite from CoverCress Inc. for performing analysis to quantify glucosinolate in leaf and seed samples.

## AUTHORS CONTRIBUTION

BG, BAJ, ME, MM, NJH, TLW, WBP, TU, MDM, RC, AND JCS: conceptualization; BG, BAJ, ME, MM, SL, MEP, NJH, TLW, WBP, TU, MDM, RC, AND JCS: methodology; BG, BAJ, ME, MM, DW, SL, MEP, NJH, TLW, WBP, TU, MDM, RC, AND JCS: investigation; WBP, TU, MDM, RC, AND JCS: resources; BG, WBP, MDM, RC, AND JCS: data curation; BG and JCS: writing - original draft; BG, BAJ, ME, MM, DW, SL, MEP, NJH, TLW, WBP, TU, MDM, RC, AND JCS: writing - review & editing; WBP, TU, RC, AND JCS: supervision; WBP, TU, MDM, RC, AND JCS: funding acquisition.

## Supplementary Materials

### MATERIALS AND METHODS

#### Seed germination and growth conditions

Pennycress seeds were surface sterilized with a brief rinse of 95% ethanol followed by a 20-minute incubation in a sterilization solution of 50% bleach and 0.01% sodium dodecyl sulfonate (SDS) for dark seeds without *TT8* mutated or a five-minute incubation for yellow seeds containing a *tt8* mutation. The seeds were then rinsed three times with sterile water and spread onto “agar growth media” consisting of 0.8% agar (BioWORLD bioPLUS Bacteriological agar) and one-half-strength Murashige and Skoog salts, in Petri dishes. The agar plates were wrapped with parafilm and placed in a Percival Scientific CU-36L5 incubator (22 °C with 16 h 4100 K fluorescent light 150–200 μE/ m^2^/ s and 8 h dark) for one week, after which seedlings were transplanted at a density of four plants per four-inch pot (Gage Dura Pots 4″ × 3–3/8″, OBC Northwest Inc. catalog no. PPG4) containing autoclaved potting soil mix of Miracle Gro Moisture Control and ProMix BX with Mycorrhizae (50/50 ratio) intermixed with 0.03 g/ four-inch pot of the insecticide Marathon (Marathon 1% Granular). Plants were grown in environment-controlled growth chambers cycling 16 h light/8 h dark (light was either 6500K fluorescent or a combination of 4100K fluorescent/incandescent lighting, 175–250 μE/ m^2^/ s light intensity), at 21 to 22°C.

Germination studies, flowering time determinations, plant height measurements, and seed dimensions analyses were performed as described in Jarvis et al. (*53*).

For field studies, seeds were either hand-planted in single five-foot rows (two-inch seed spacing, one-eighth inch depth) or in five-foot by 10-foot plots planted by drilling (one-eighth inch depth, 15 cm row spacing after tilling and harrowing, 2.6 gm seed per plot) or broadcasting after light vertical tilling. Only lines meeting USDA APHIS requirements were field planted. Planting occurred in late September/early October, and hand or research plot combine harvesting in late May/early June.

#### Vectors

The *TaHAG1* (TAV2_LOCUS5186), *TaHAG2* (GAKE01003245.1; TAV2_LOCUS20927), *TaHAG3* (GAKE01004525.1; TAV2_LOCUS20427), *TaMYC3* (GAKE01000848.1; TAV2_LOCUS6314), and *TaTT8* (GAKE01020678, GAKE01031871; TAV2_LOCUS19620) coding sequences (CDS) were obtained by performing Basic Local Alignment Search Tool (BLAST) searches of the *T. arvense* transcriptome shotgun assembly database (TSA; *54*) at the National Center for Biotechnology Information (NCBI) and the MN106 and Spring 32-10 reference genomes (*55*, *56*), using the *AtHAG1*/*AtMYB28* (AT5G61420), *AtHAG2*/*AtMYB76* (AT5G07700), *AtHAG3*/*AtMYB29* (AT5G07690), *AtMYC3* (AT5G46760), and *TT8* (AT4G09820) CDS as queries. Note that not all *Thlaspi arvense* sequences were accurate, requiring hand annotation.

CRISPR/SpCas9 binary vector constructs consisting of 4 guide RNAs or 2 guide RNAs in binary vector pHEE401E were assembled using methods similar to those as described by Wang et al. (*57*). The nucleotide sequences of all protospacers under consideration were queried against the entire *Thlaspi arvense* genome sequences using BLAST. Only those that had less than 90 % sequence identity with off target locations were chosen for use. The completed constructs were transformed into *Agrobacterium tumefaciens* strain GV3101 and stably introduced into pennycress using the *Agrobacterium*-mediated floral dip vacuum infiltration method described in McGinn et al., (*51*).

The ethylmethane sulfonate (EMS) mutant lines, in the MN106 background, were part of the mutant collection described in Chopra et al., (*50*).

#### Seed total oil, protein, fiber and fatty acid quantification

Seed compositions (crude protein along with amino acids, crude fat, crude fiber along with non-detergent fiber (NDF) and acid detergent fiber (ADF) were quantified by the University of Missouri-Columbia Agricultural Experiment Station Chemical Laboratories (ESCL) using American Oil Chemist Society (AOCS) and Association of Official Analytical Chemists (AOAC), International methodologies. Total seed oil content was quantified using a rapid nondestructive pulsed nuclear magnetic resonance (NMR) method on 150 mg or 450 mg seeds, employing a Bruker Minispec PC 120, 180-mm absolute probehead. Pennycress seed oil fatty acid composition was determined using a sodium methoxide-based methylation protocol to generate fatty acid methyl esters (FAME) from TAGs followed by detection/quantification using Gas Chromatography-Flame Ionization Detection (GC-FID) with a Supelco SP2380 column, as described in Moser et al., (*58*).

#### Seed and cauline leaf glucosinolate content determination

Seed glucosinolate content was determined as described in Chopra et al. (*59*), using both a non-destructive Near Infrared Spectroscopic (NIRS) analyses employing a Metrohm NIRSystems NIRS XDS MultiVial Analyzer (200 mg of seed placed within 1/3 Dram Clear Glass Vials), and a more precisely quantitative wet chemistry method. For the latter method, 0.05 g seed, four 3.175 mm ball bearings, and 1 mL 80% methanol solution were put into each 2 mL screw cap microcentrifuge tube and ground 300s using a bead beater. The ground seed tissue was pelleted by centrifuging at 2500 g for 5 min. 80 µL supernatant was moved to a new tube and evaporated completely, then 400 µL water (5 times dilution) was added and mixed well followed by filtration using a 0.45 µm nylon syringe filter and storage in a HPLC vial with insert. Chromatographic separation was performed on 20 μL injected onto a C18 Phenomenex SphereClone™ 5 µm ODS(2) column (250 x 4.6 mm, 80 Å) at a column temperature of 22°C using an Agilent HPLC system. The mobile phase was composed of 0.05 M ammonium sulfate (NH2)2SO4 with 1 mL/min isocratic elution for 15 min. Peaks were measured using an Agilent G1315A diode array detector (DAD) detector set at 229 nm. Sinigrin standards (0, 0.01, 0.1, 0.25, 0.5, 1, 2mM) were diluted from a 10 mM sinigrin stock in water by serial dilution. The sinigrin peak eluted at around 5.5 min.

#### RNAseq analysis

Cauline leaves of the same age from growth chamber-grown plants that had started to flower were collected (100 mg tissue per plant, five plants per line), and total RNA extracted using the Sigma-Aldrich Spectrum™ Plant Total RNA Kit. RNA sample quality control, mRNA library preparation, and generation of at least six gigabases (6 G) of raw sequence data per sample using a NovaSeq X Plus Series (PE150) were carried out by Novogene Corporation, Inc. Adapter trimming was conducted using BBduk (BBMAP_39.01, Joint Genome Institute) (*60*). Trimmed FASTQ v0.12.1 (*61*) files were aligned to the reference genome using STAR (version 2.7.11b) (*62*) with the parameter “quantMode GeneCounts”, which quantifies the number of reads per gene. MultiQC v1.22.1 was used to aggregate output files and obtain a summary of results (*63*). Raw counts data were filtered to retain genes with at least 10 counts across all samples. DESeq2’s (*64*) standard workflow was followed and Principal Component Analysis (PCA) was used to visualize sample clustering. Differentially expressed genes (DEGs) were identified using a log₂ fold-change (LFC) threshold of 1.0. To control for multiple testing, FDR correction was applied using the Benjamini-Hochberg method (alpha = 0.05) (*65*). Volcano plots and heat maps were generated to visualize significantly upregulated and downregulated genes. All statistical analyses and visualizations were conducted in RStudio v4.2.1.

#### Seed persistence trials

Fields trials at Normal, IL (40.527084, −89.007056 on Catlin silt loam soil) and Lexington, IL (40.674040, −88.767771 on Andres silt loam and Drummer and El Paso silty clay loam soils) were conducted comparing black-seeded and golden-seeded (*tt8* knockout mutation-containing) pennycress. Agronomic and breeding plots were established in the fall, and pennycress was harvested the following spring. The footprints of those plots were monitored for an additional year at one site and for an additional two years at the other. Volunteer pennycress from the prior experiment was allowed to establish, overwinter, and flower. The mature plants were documented for the number of plants containing black-seeded pennycress versus golden-seeded pennycress.

#### Statistical analyses

All statistical analyses were performed in RStudio v4.2.1. For Figs. 3 and 4 and Figs. S1, S5, S7, S9, S10, and S11, a one-way ANOVA was conducted to assess differences across groups. To check for homogeneity of variances, Levene’s test was performed, which showed no significant violation (p > 0.05). Post hoc pairwise comparisons were conducted using the Tukey/Sidak adjustment for multiple testing. Results were considered significant for p-values < 0.05. **For Figs. S4 and S6, a** Kruskal-Wallis test was performed to assess differences across groups, and post-hoc pairwise comparisons were conducted using Dunn’s test with Sidak adjustment for multiple comparisons. Results were considered significant for p-values < 0.05.

**fig. S1.**
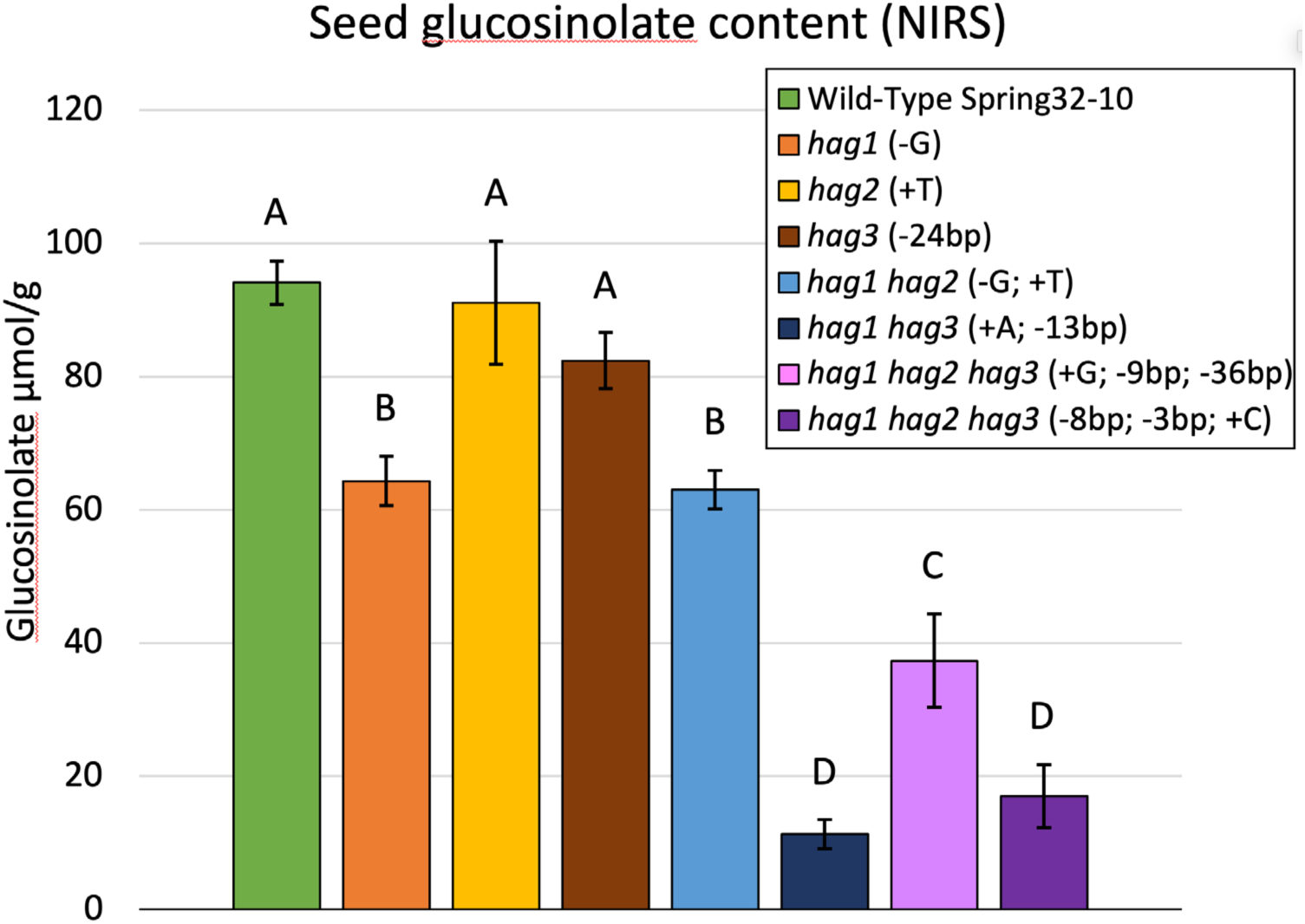
Seed glucosinolate content in pennycress *hag* single, double, and triple mutants (growth chamber grown), as determined by near infrared spectroscopic (NIRS) analysis. Mutations in each allele are in parentheses after the names of the genes mutated. Error bars represent standard deviations. Different letters above columns represent statistical differences (*p*<0.05) based on one-way ANOVA and Tukey’s test. n=4.

**fig. S2.**
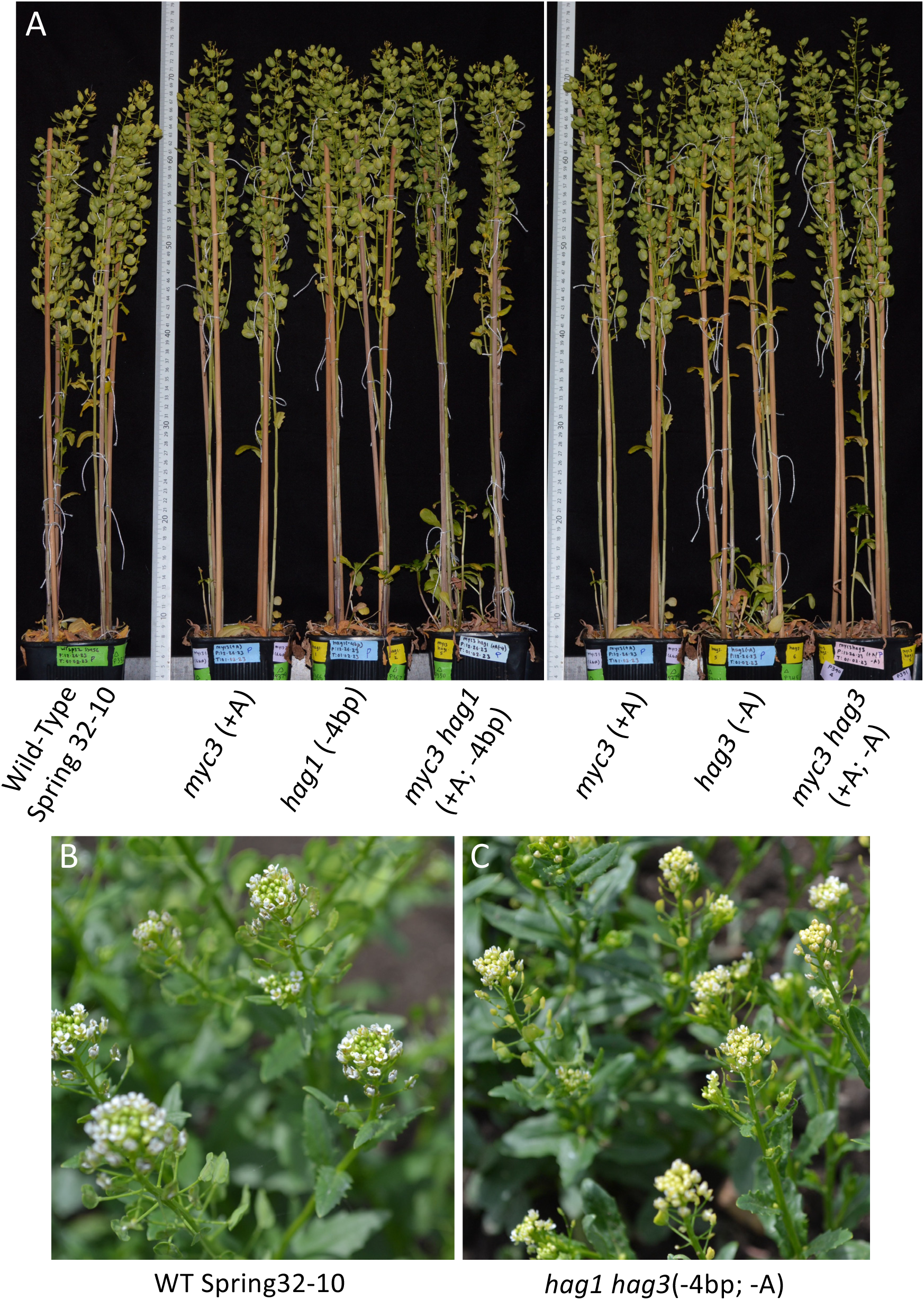
(**A**) Growth chamber-grown mutant plants compared to wild-type Spring 32-10 (four-inch pots each containing four plants). Mutations in each allele are in parentheses after the names of the genes mutated. (**B, C**) Field growing wild-type Spring 32-10 (**B**) and *hag1 hag3* double knockout mutant plants (**C**). Note the yellowish color of the *hag1 hag3* floral tissues and developing silicles in (**C**).

**fig. S3.**
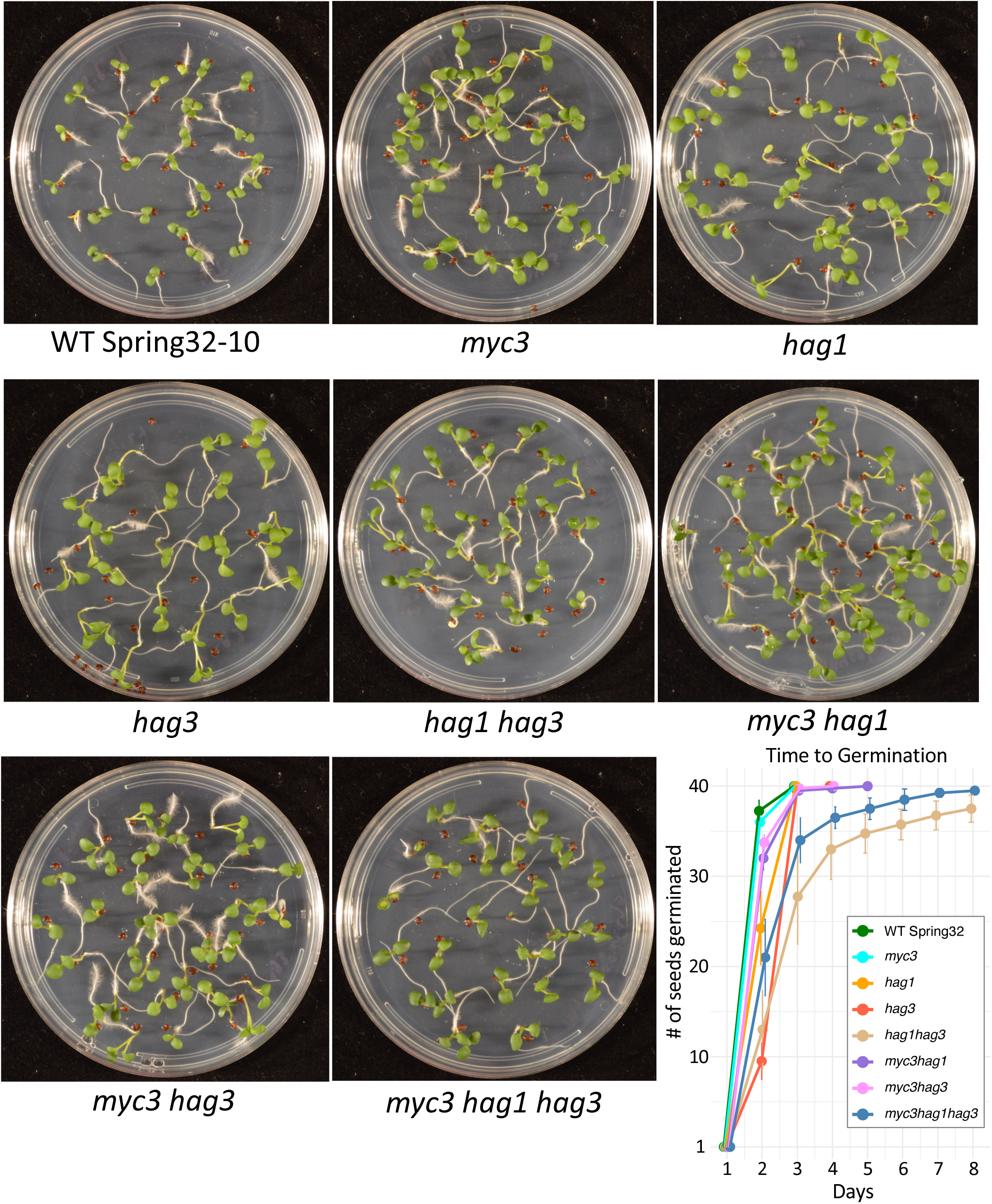
Seed germination of the various glucosinolate single mutants and mutant combinations compared to wild-type (WT Spring 32-10). Pictures were taken 7 days after plating surface sterilized seeds on 1/2-strength MS agar medium. n=40 seeds, 4 biological reps. Bars are standard deviation. See Fig. 3 legend and data S1 file for mutations information.

**fig. S4.**
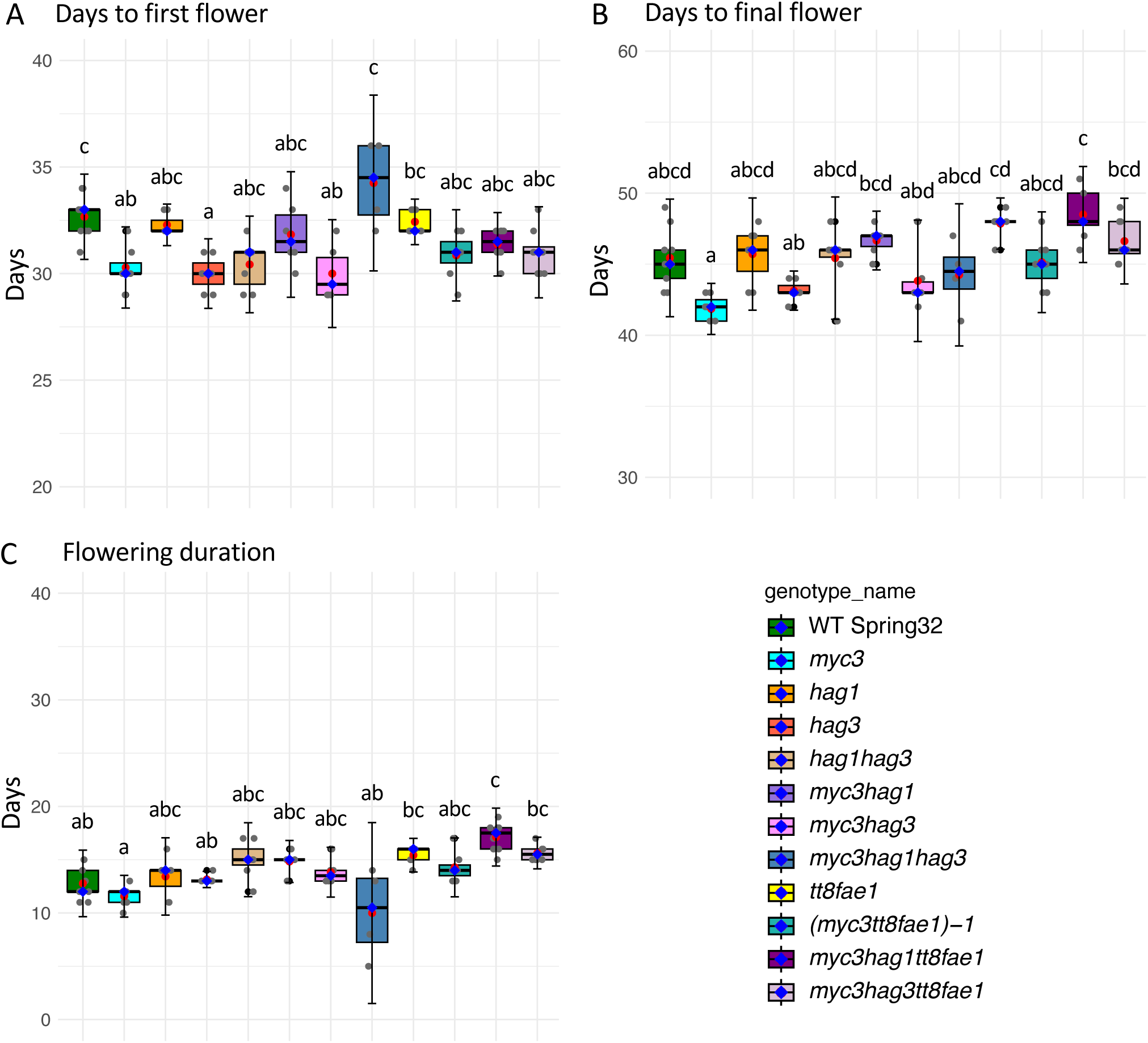
Box plot graphs of the number of days to first flower (**A**), last flower (**B**), and flowering duration (**C**) for the various mutant lines containing domestication traits versus wild type (growth chamber grown). All lines were grown together in a growth chamber, four plants per four-inch pot. n=8 plants. Measurements in all three graphs are from the same plants. Grey dots represent each sample data, red dots estimated marginal means, and blue dots median. Means not sharing any letter are significantly different by the Sidak test at the 5% level of significance. See Fig. 3 legend and data S1 file for mutations information.

**fig. S5.**
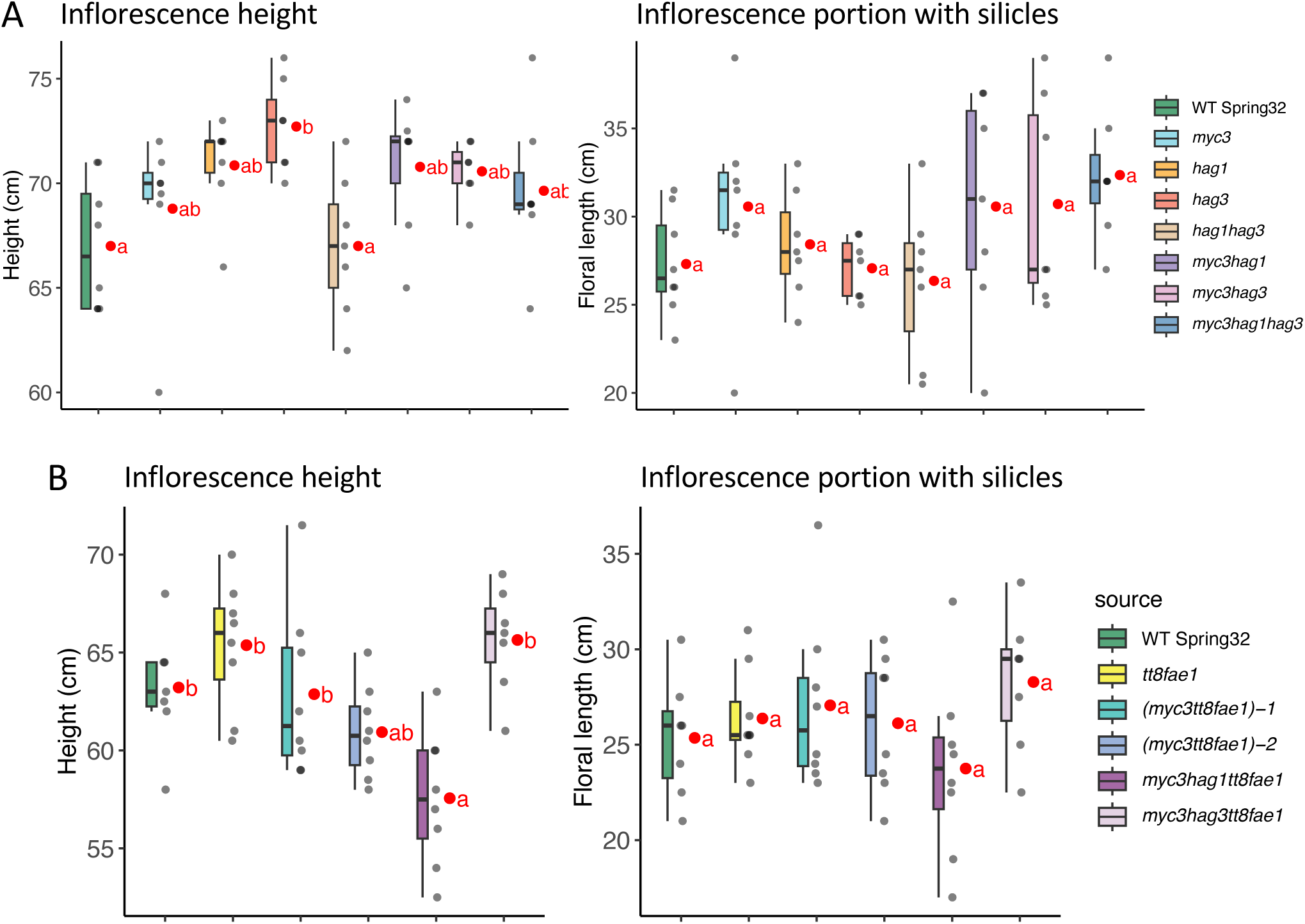
(**A, B**) Box plot graphs of the heights of fully mature inflorescence stems (left two graphs) and the portions of the inflorescence stems having silicles (seed pods; right two graphs). Lines in each graph were grown together in a growth chamber, four plants per four-inch pot. n=8 plants. Grey dots represent each sample data and red dots estimated marginal means. Means not sharing any letter are significantly different by the Sidak test at the 5% level of significance. See Fig. 3 legend and data S1 file for mutations information.

**fig. S6.**
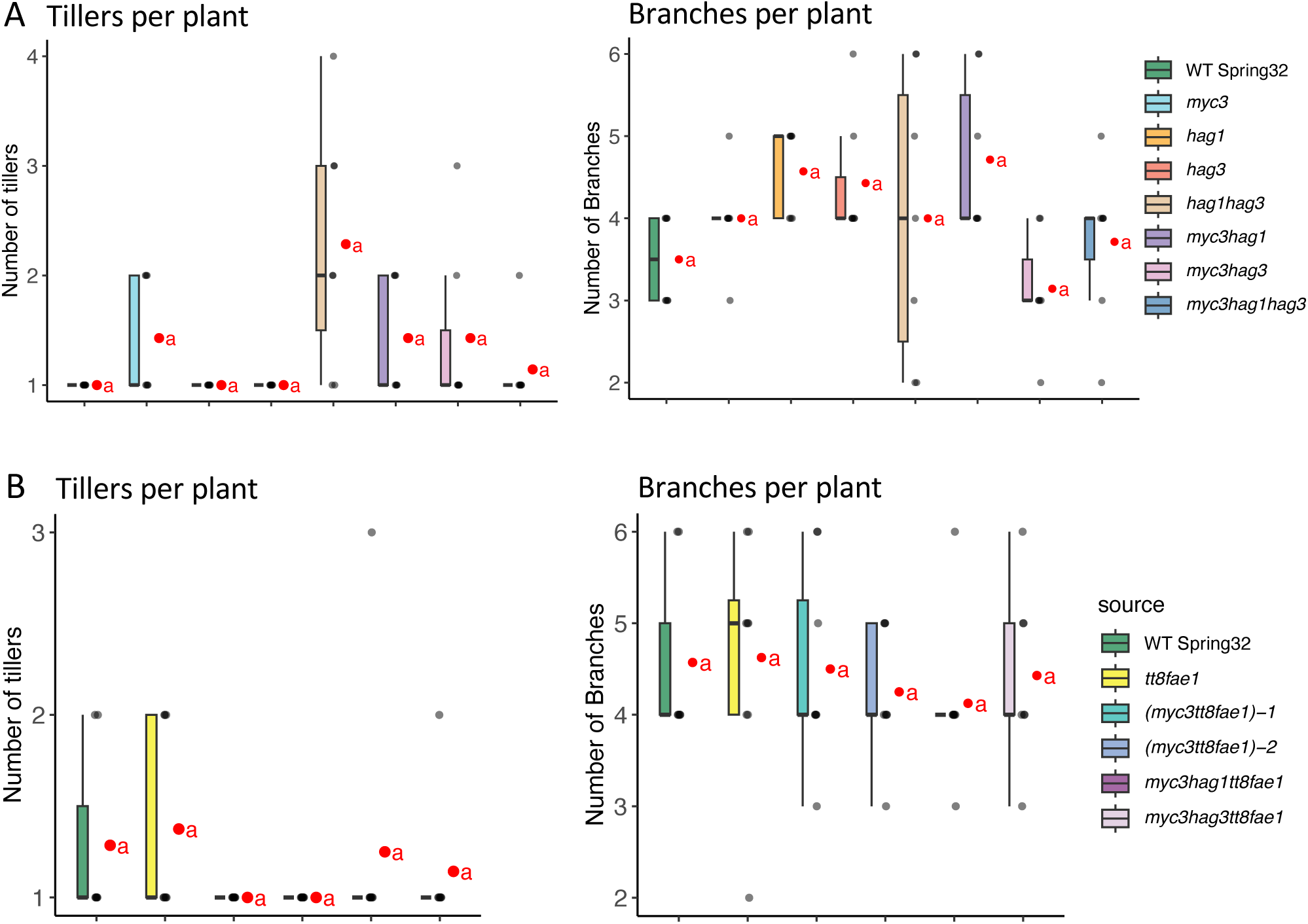
Box plot graphs of the numbers of tillers and branches of plants (growth chamber grown) containing the various glucosinolate gene mutations without (**A**) or when combined with the *tt8* and *fae1* mutations (**B**). n=8 plants. Grey dots represent each data and red dots estimated marginal means. Dunn’s post-hoc test with Sidak adjustment (non-parametric test) was used for pairwise comparisons. No significant differences were determined (*p*<0.05). See Fig. 3 legend and data S1 file for mutations information.

**fig. S7.**
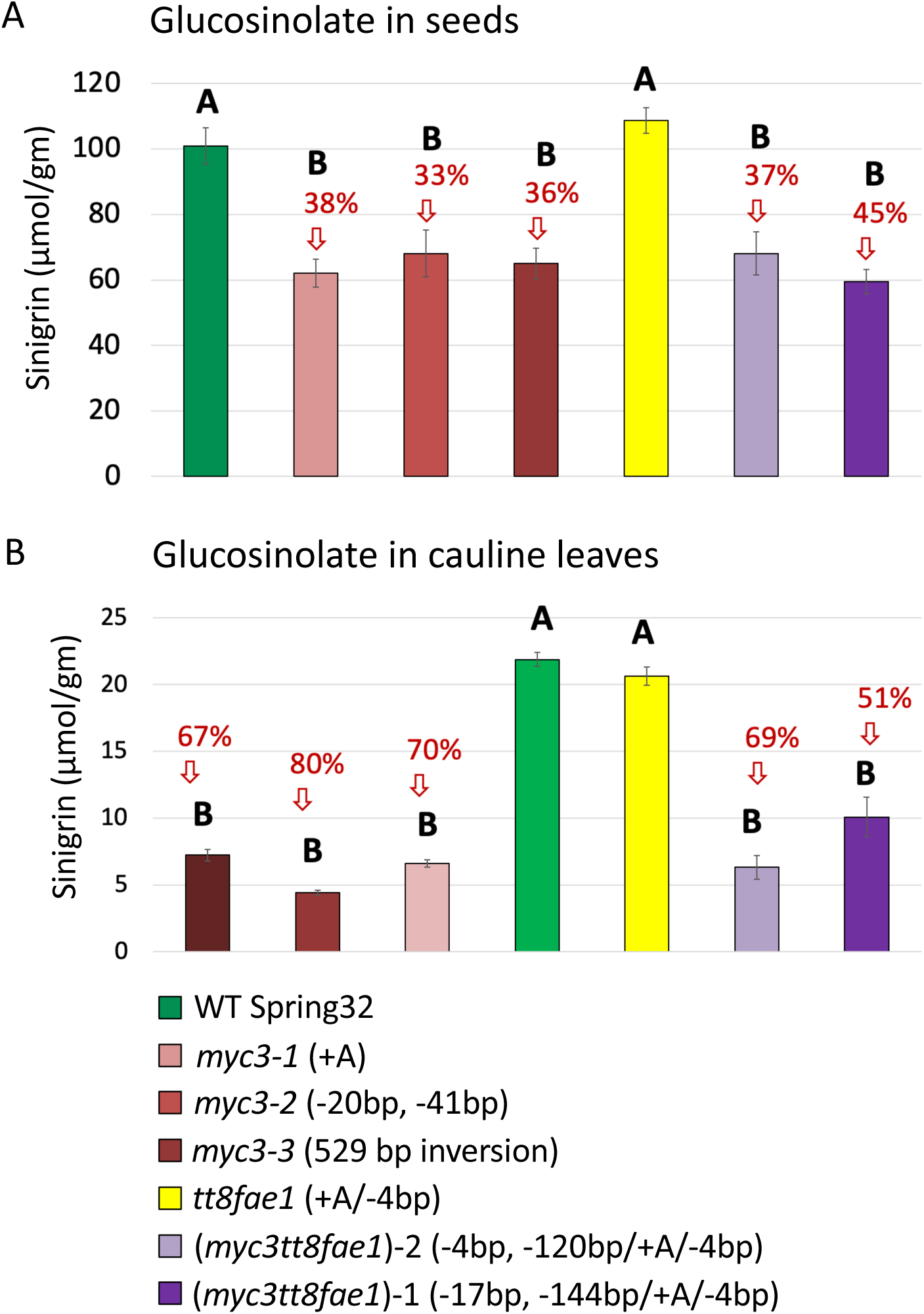
(**A, B**) Graphs of glucosinolate (sinigrin) content in mature seeds (**A**) and in cauline leaves from flowering plants (**B**) of *myc3* mutants without or when combined with *tt8* and *fae1* knockout mutations. Mutations in each allele are in parentheses after the names of the genes mutated. Note that *myc3-2* has two separate deletions (−20 bp and −41 bp) within the *MYC3* gene. Percentages represent reductions versus the corresponding wild-type (WT Spring32) and *tt8 fae1* controls. Means not sharing any letter are significantly different by one-way ANOVA plus Tukey’s test at the 5% level of significance. n=8

**fig. S8.**
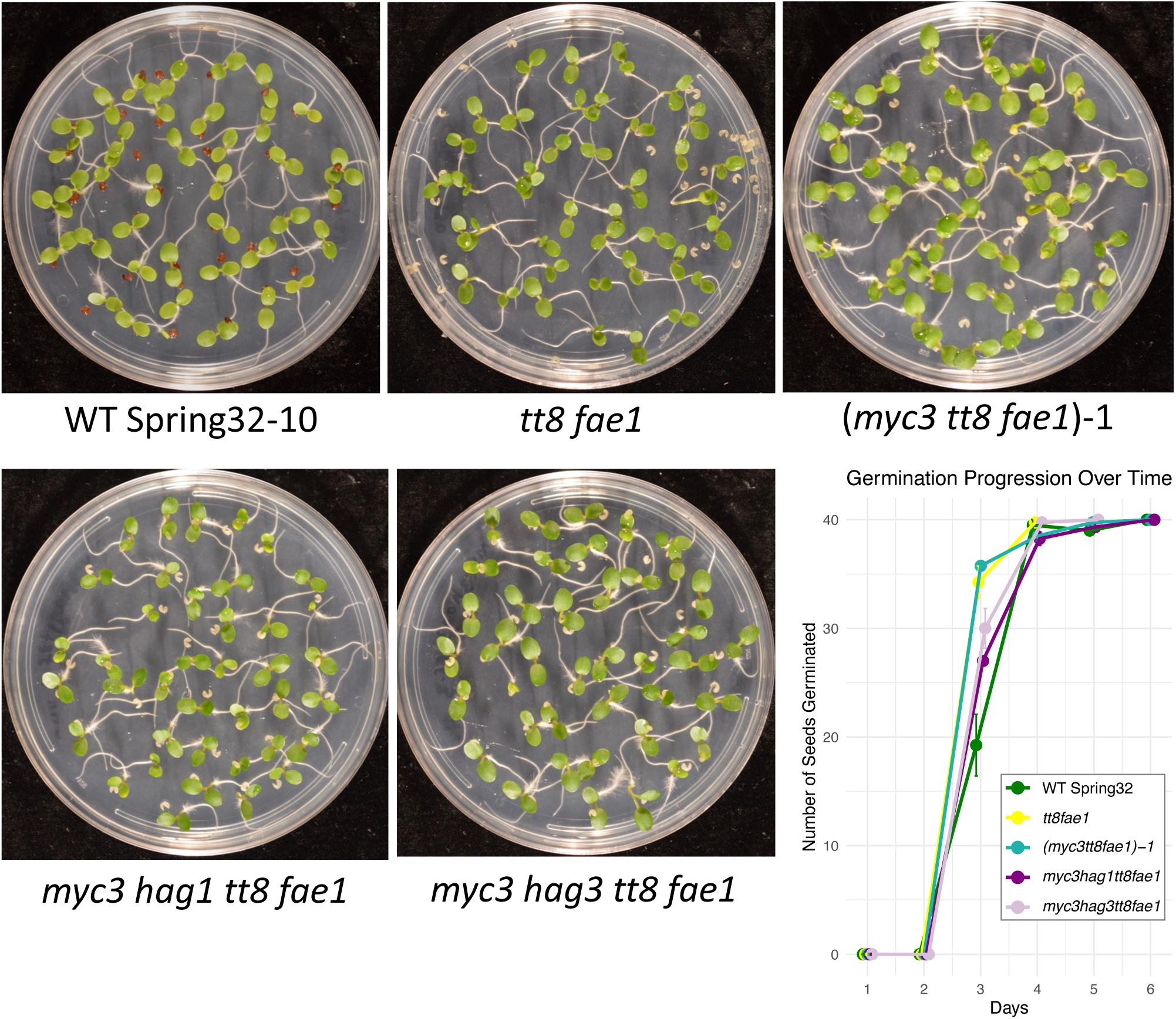
Seed germination of mutant lines compared to wild type (WT Spring 32-10). Pictures were taken 7 days after plating surface sterilized seeds on 1/2-strength MS agar medium. n=40 seeds, 4 biological reps. Bars are standard deviation. See Fig. 3 legend and data S1 file for mutations information.

**fig. S9.**
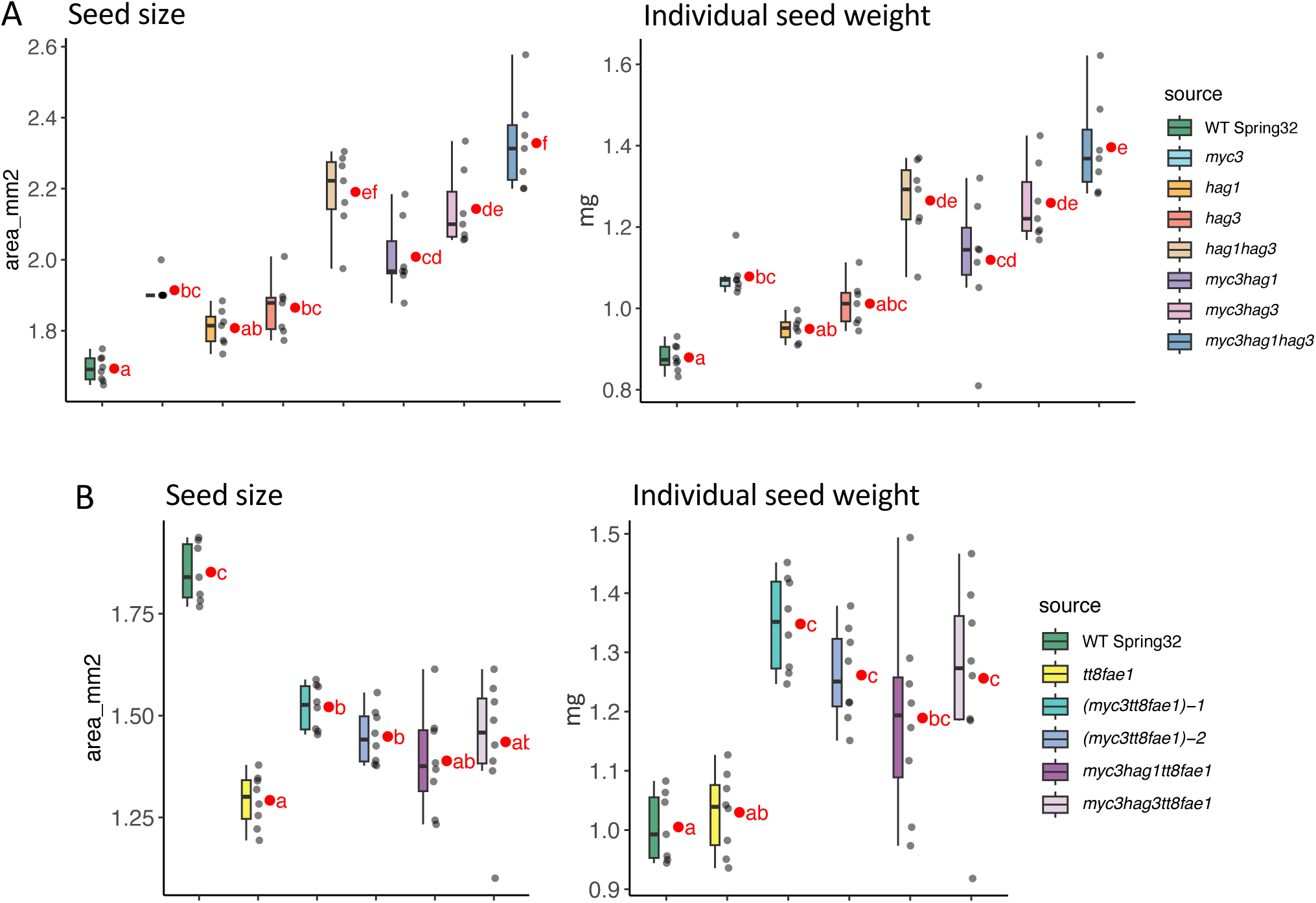
Box plot graphs of the area per seed and the average weight per seed for the various glucosinolate mutants (growth chamber grown) without (**A**) or when combined with *tt8* and *fae1* knockout mutations (**B**). n=8. Grey dots represent each sample data and red dots estimated marginal means. Means not sharing any letter are significantly different by the Sidak test at the 5% level of significance. Note that *myc3* mutations result in seeds weighing more than wild type. Note that *tt8* seeds are two-dimensionally smaller than wild type but are the same volume (not shown) as a consequence of being relatively more spherical and not flattened. See Fig. 3 legend and data S1 file for mutations information

**fig. S10.**
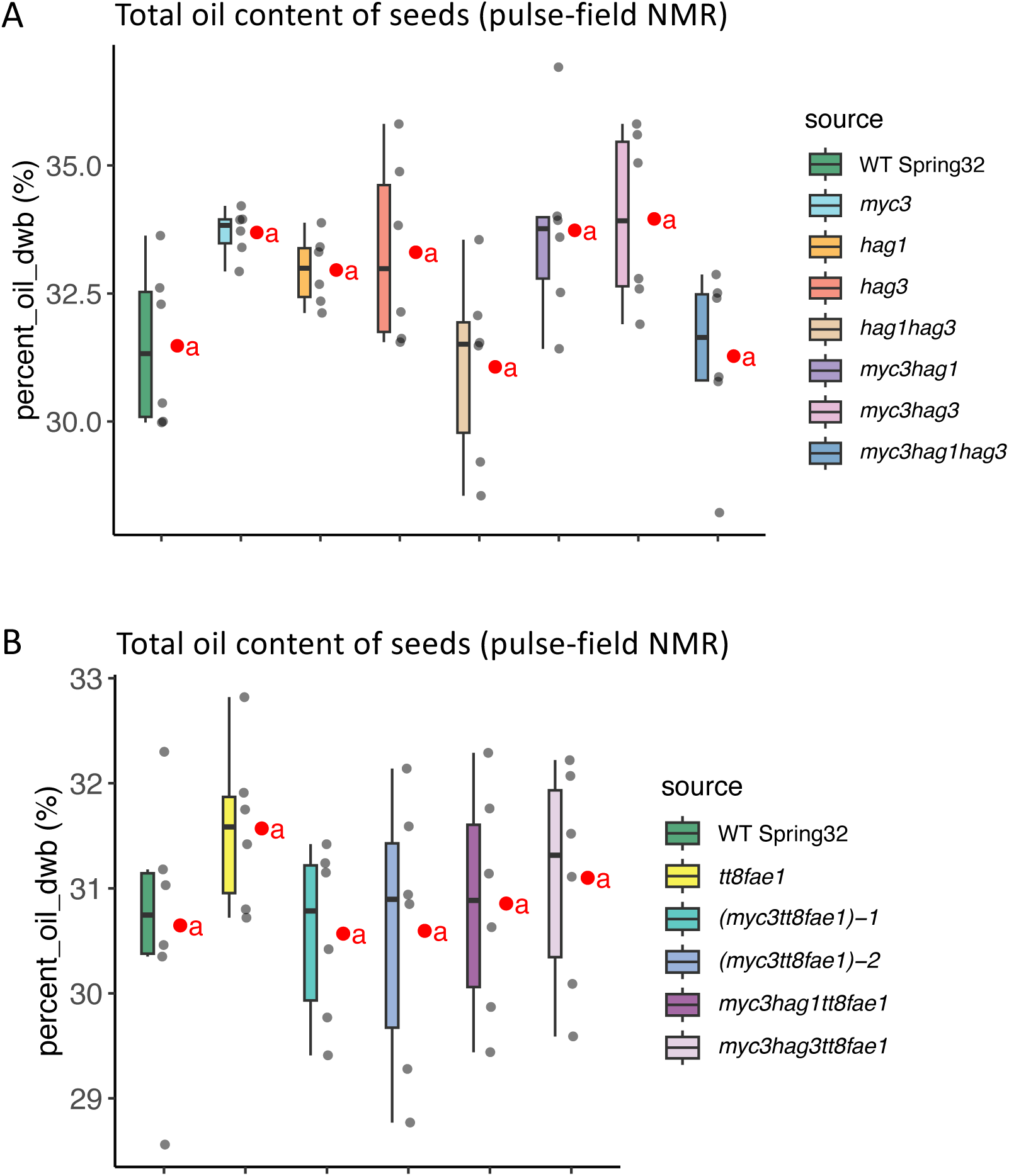
Box plot graphs of the total oil content of seeds from the various glucosinolate mutants (growth chamber grown) without (**A**) or when combined with *tt8* and *fae1* knockout mutations (**B**), as determined by pulse field NMR analysis. n=6. Grey dots represent each sample data and red dots estimated marginal means. There were no significant differences between each line based on the Sidak test at the 5% level of significance. Note that the content shown here was calculated for 450 mg of seeds, not a single seed basis. See Fig. 3 legend and data S1 file for mutations information.

**fig. S11.**
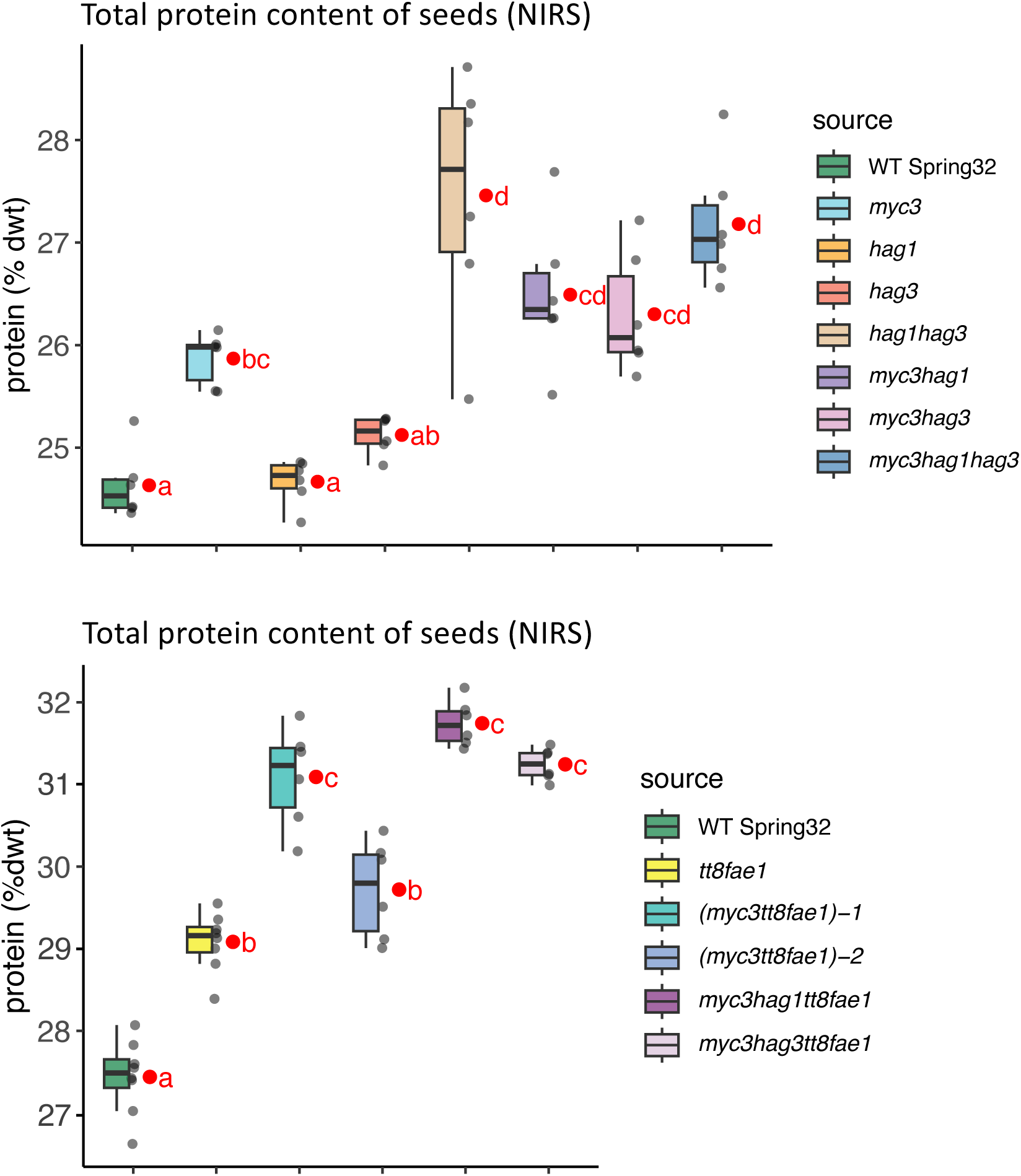
Box plot graphs of the total protein content of seeds from the various glucosinolate mutants (growth chamber grown) without (**A**) or when combined with *tt8* and *fae1* knockout mutations (**B**), as determined by NIR spectroscopic analysis (NIRS). n=6. Grey dots represent each sample data and red dots estimated marginal means. Means not sharing any letter are significantly different by the Sidak test at the 5% level of significance. Note that the content shown here was calculated on bulk seeds, not a single seed basis. See Fig. 3 legend and data S1 file for mutations information.

**fig. S12.**
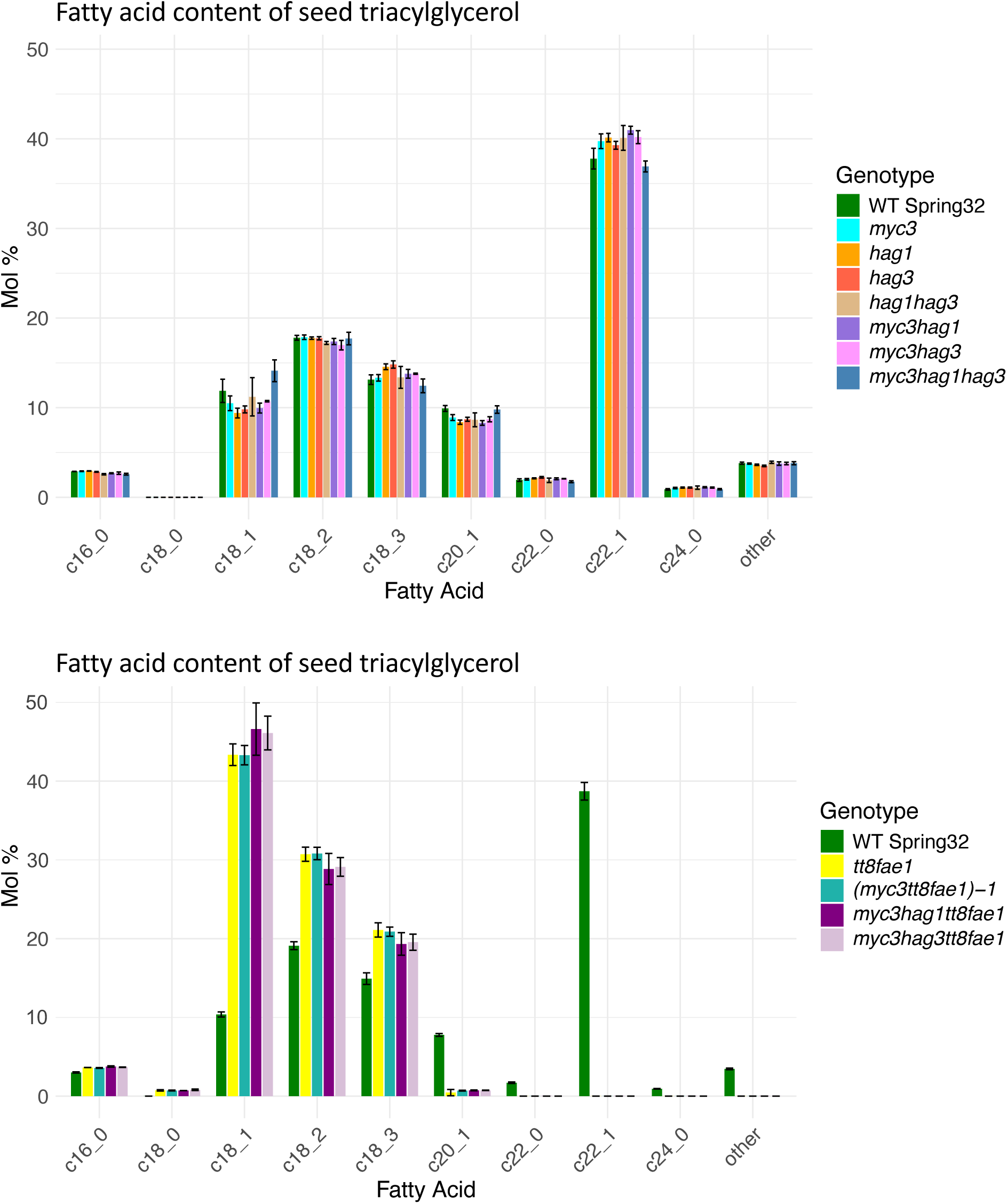
Graphs of the fatty acid content of seed oil (triacylglycerol) from the various glucosinolate mutants (growth chamber grown) without (**A**) or when combined with *tt8* and *fae1* knockout mutations, in mole percents (**B**). Error bars represent standard deviations. n=6. See Fig. 3 legend and data S1 file for mutations information.

**fig. S13.**
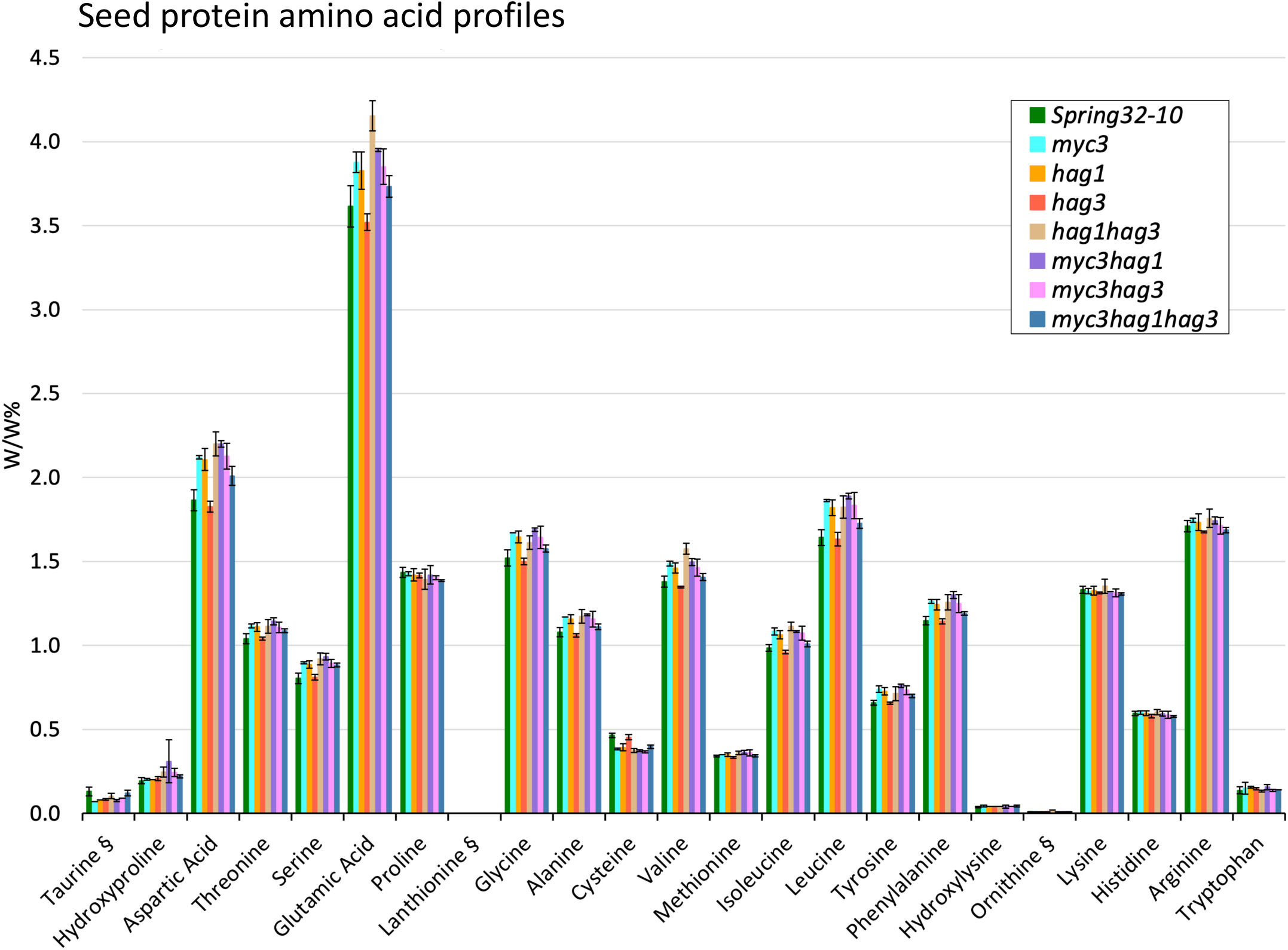
Amino acid compositions of crude protein extracted from seeds of the glucosinolate single mutants and combinatorial mutants (field grown) along with wild-type Spring 32-10. W/W% is grams amino acid per 100 grams of crude protein. Amino acids delineated with a section symbol (§) are non-proteinogenic. Error bars represent standard deviations. n=3.

**fig. S14.**
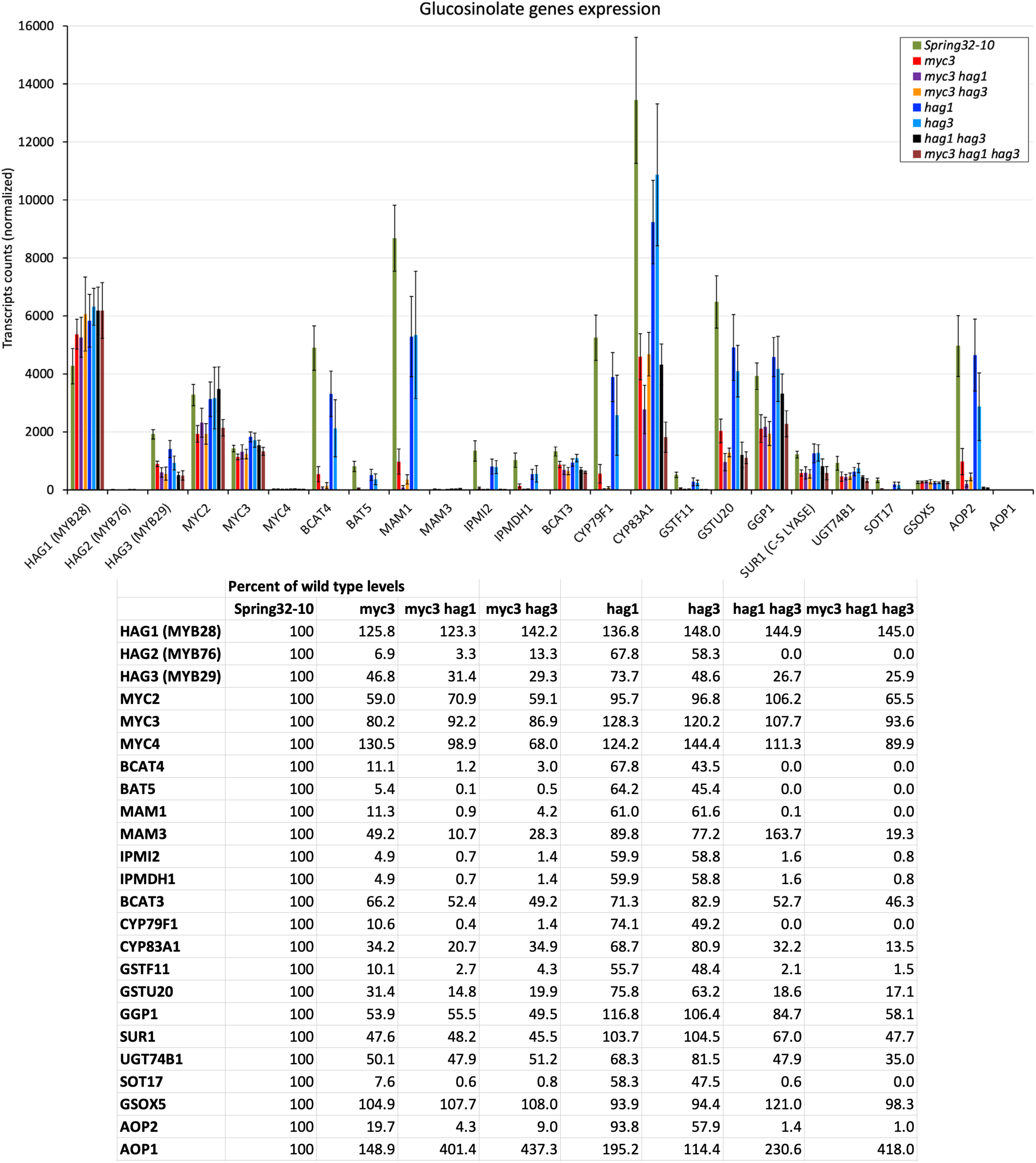
(**A**) RNAseq-derived normalized transcript counts, from cauline leaf tissue, of genes encoding transcription factors and biosynthetic enzymes involved in glucosinolate biosynthesis in the various glucosinolate single and combinatorial mutants versus wild type. Columns are means and bars represent St. Dev. (**B**) Table showing transcripts counts as percents of wild type levels.

**fig. S15.**
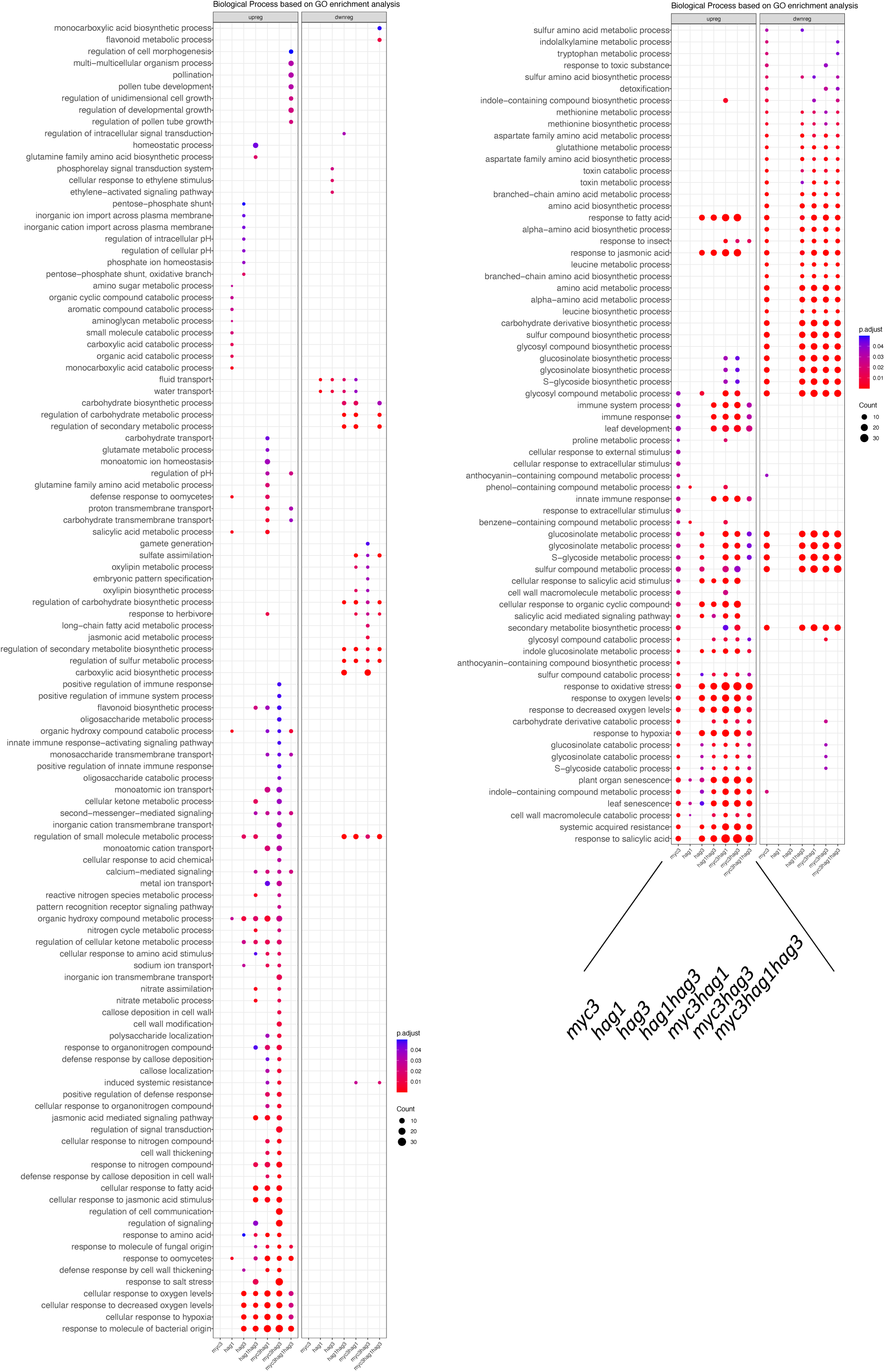
Gene ontology (GO) enrichment analysis sorted by biological process, for transcripts isolated from cauline leaves of the various mutants relative to wild type.

**fig. S16.**
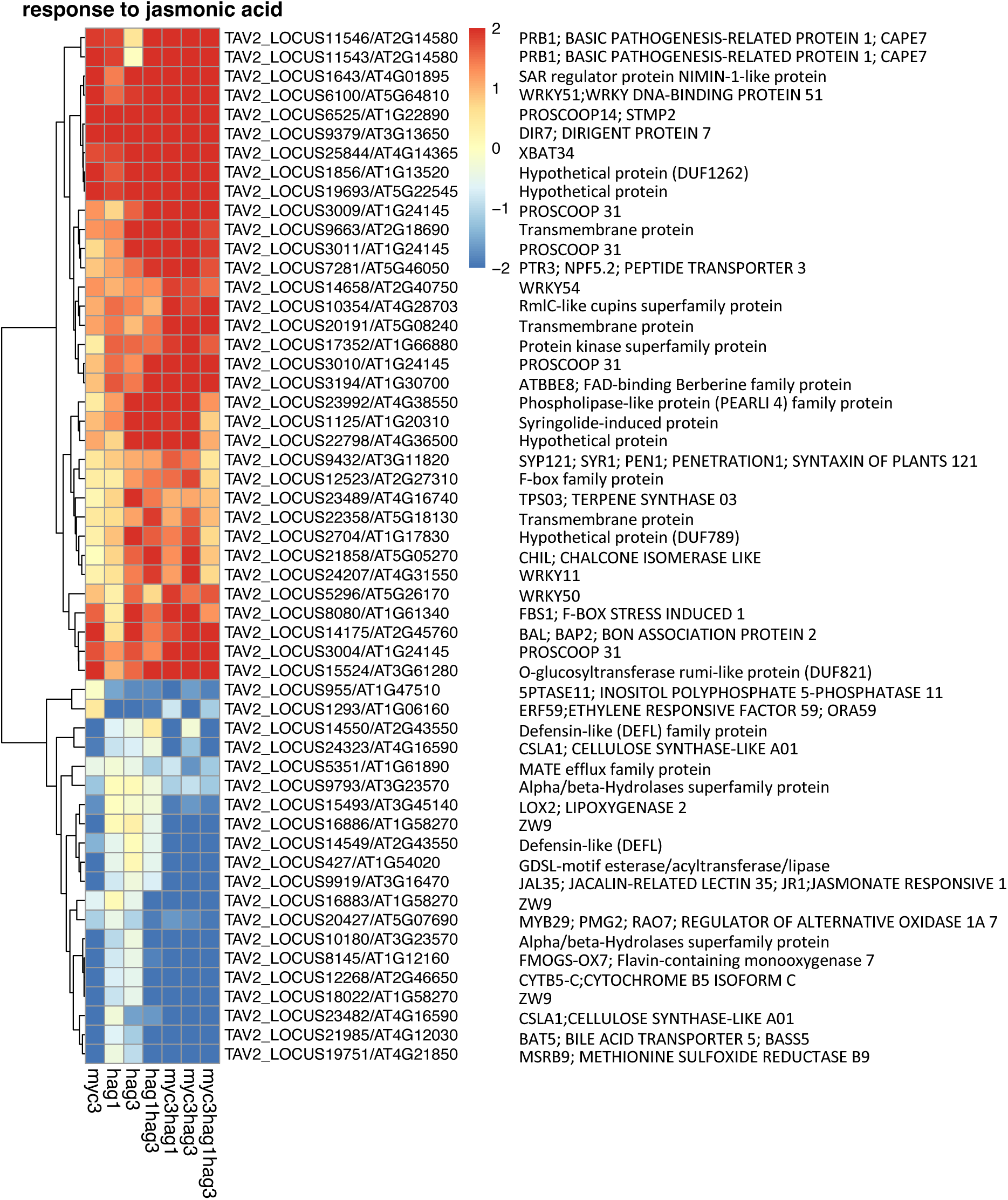
Relative expression changes of genes predicted to be linked to responses to jasmonic acid. Heat map shows log2 fold changes (threshold of 2) in transcript levels in pennycress mutants’ cauline leaf tissue relative to wild type. Pennycress gene I.D.s and closest Arabidopsis homologue I.D.s are listed to the right of the corresponding gene expression levels; mutant names listed below. Positive and negative values represent increases and reductions, respectively.

**fig. S17.**
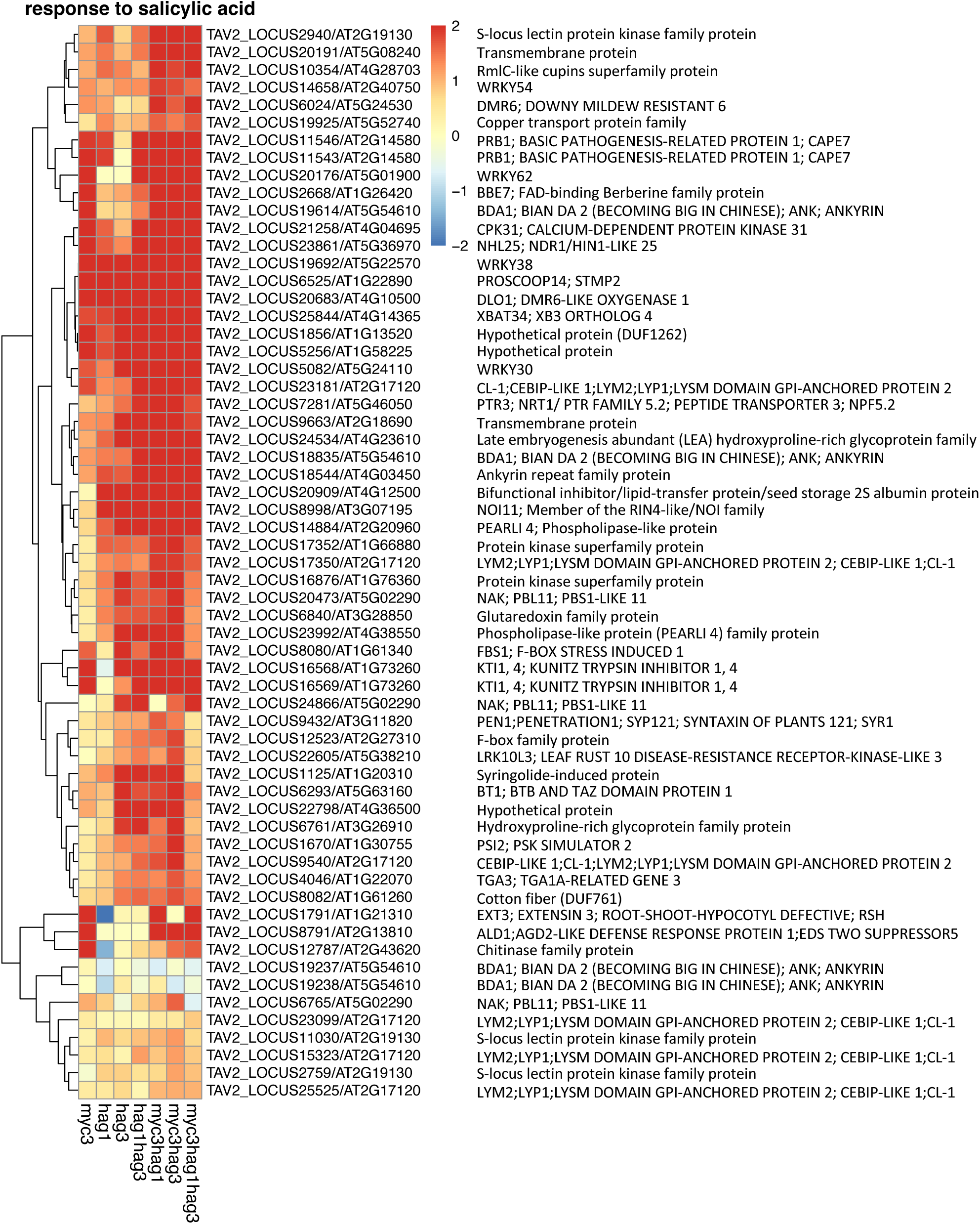
Relative expression changes of genes predicted to be linked to responses to salicylic acid. Heat map shows log2 fold changes (threshold of 2) in transcript levels in pennycress mutants’ cauline leaf tissue relative to wild type. Pennycress gene I.D.s and closest Arabidopsis homologue I.D.s are listed to the right of the corresponding gene expression levels; mutant names listed below. Positive and negative values represent increases and reductions, respectively.

**fig. S18.**
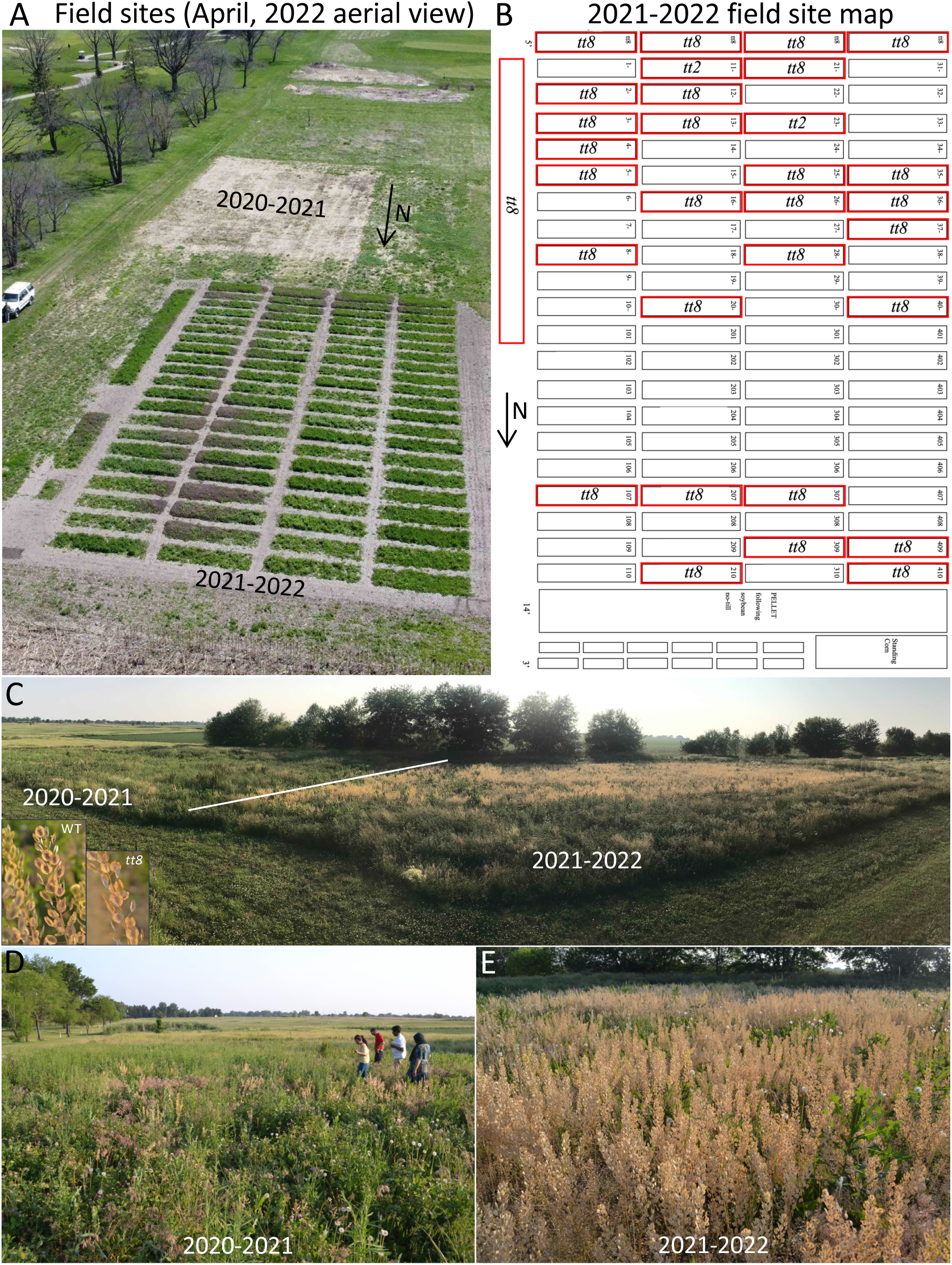
Pennycress *tt8*-containing yellow seeded plants have very low persistence compared to wild-type dark-seeded plants. **(A)** Aerial view of the 2020-2021 and 2021-2022 field sites (picture taken April, 2022). **(B)** Plots map for the fall 2021 to spring 2022 (2021–2022) field site. *tt8* plots are highlighted in red. Arrow points north. **(C)** Panoramic view of field sites (picture taken June, 2023, at the time of walking through the plots). **(D, E)** Walking the plots to score any yellow seeded (*tt8*-containing) plants. Six yellow seeded pl}ants were found in the 2021-2022 field site location **(C, E)** whereas none were found in the 2020-2021 field site **(C, D)**. Note that substantially fewer pennycress plants grew in the 2020-2021 field site location signifying wild pennycress is outcompeted by other weeds over time.

**fig. S19.**
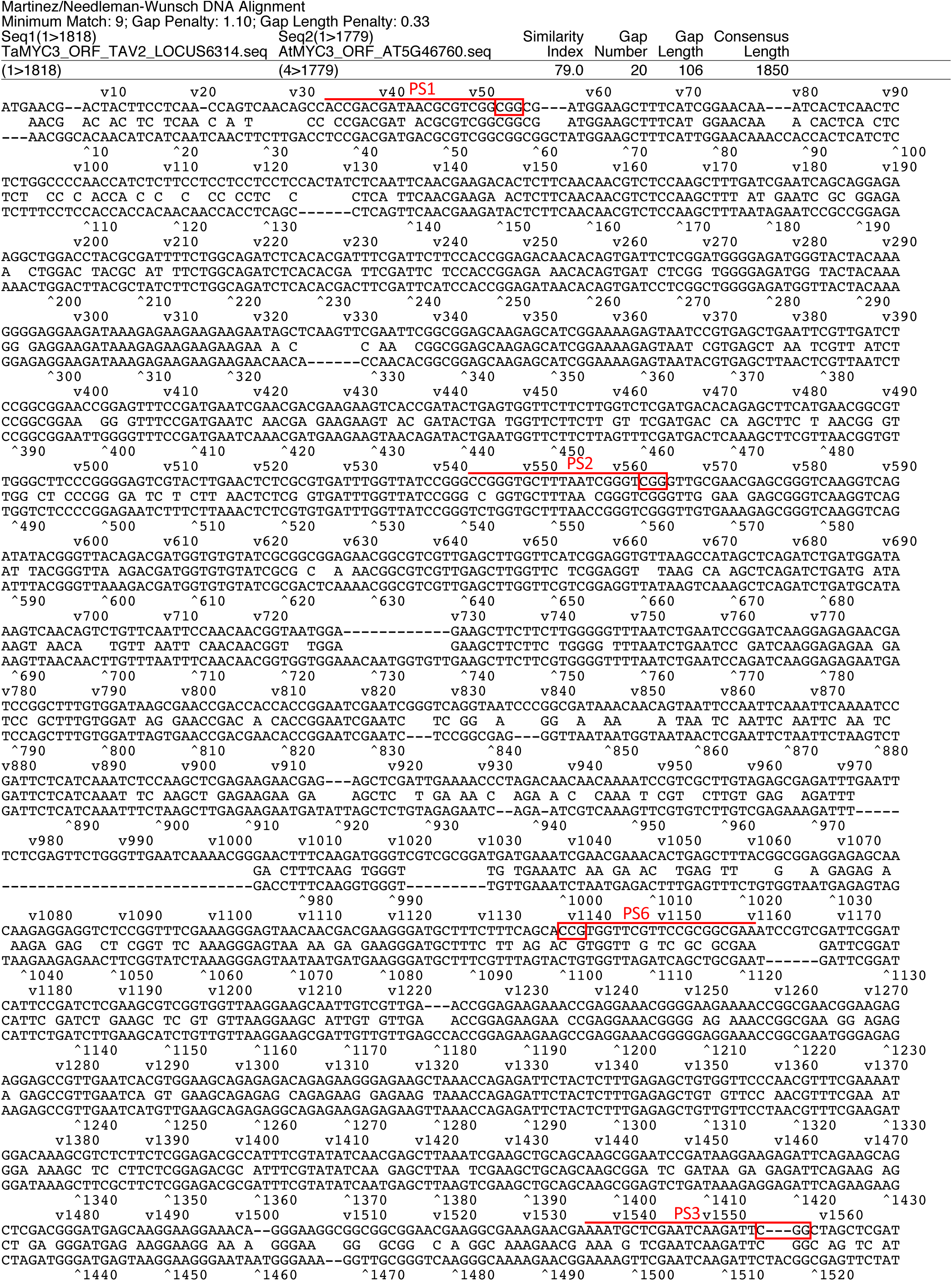
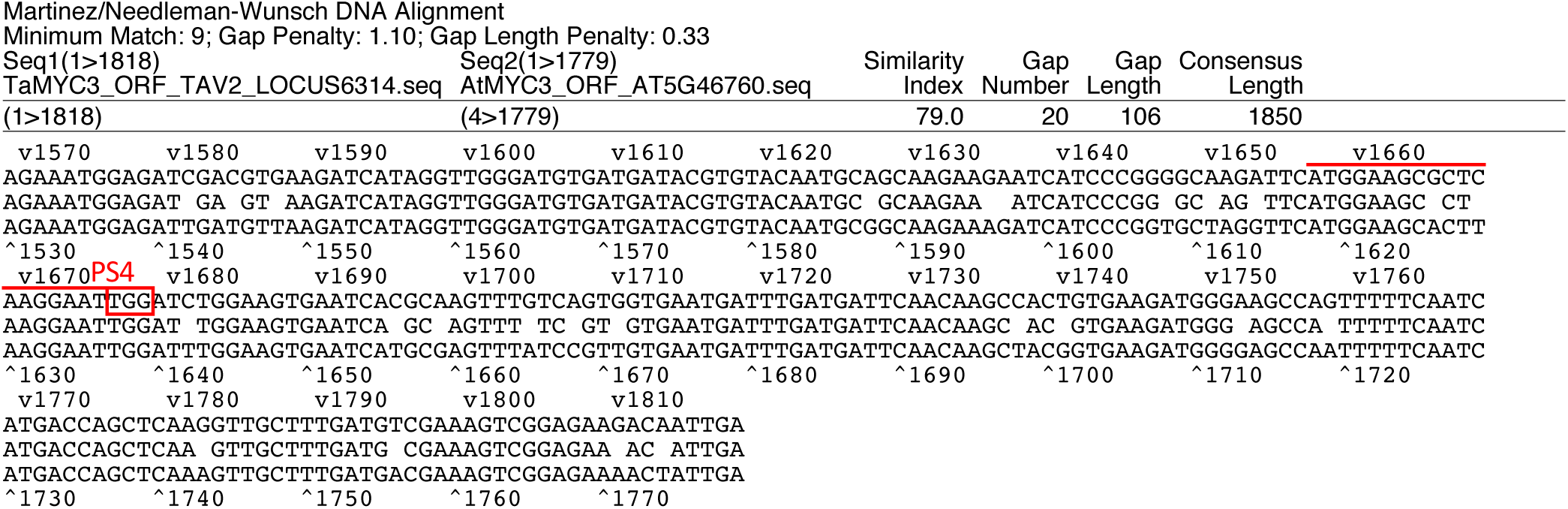
Nucleotide sequence alignment of the pennycress *TaMYC3* (TAV2_LOCUS6314) open reading frame (ORF; top sequence) to that of Arabidopsis *AtMYC3* (AT5G46760; bottom sequence). Letters between the top and bottom sequences denote identity. Locations of the protospacers used for CRISPR gene editing are marked with red lines along with the corresponding PAM sites (boxed in red).

**fig. S20.**
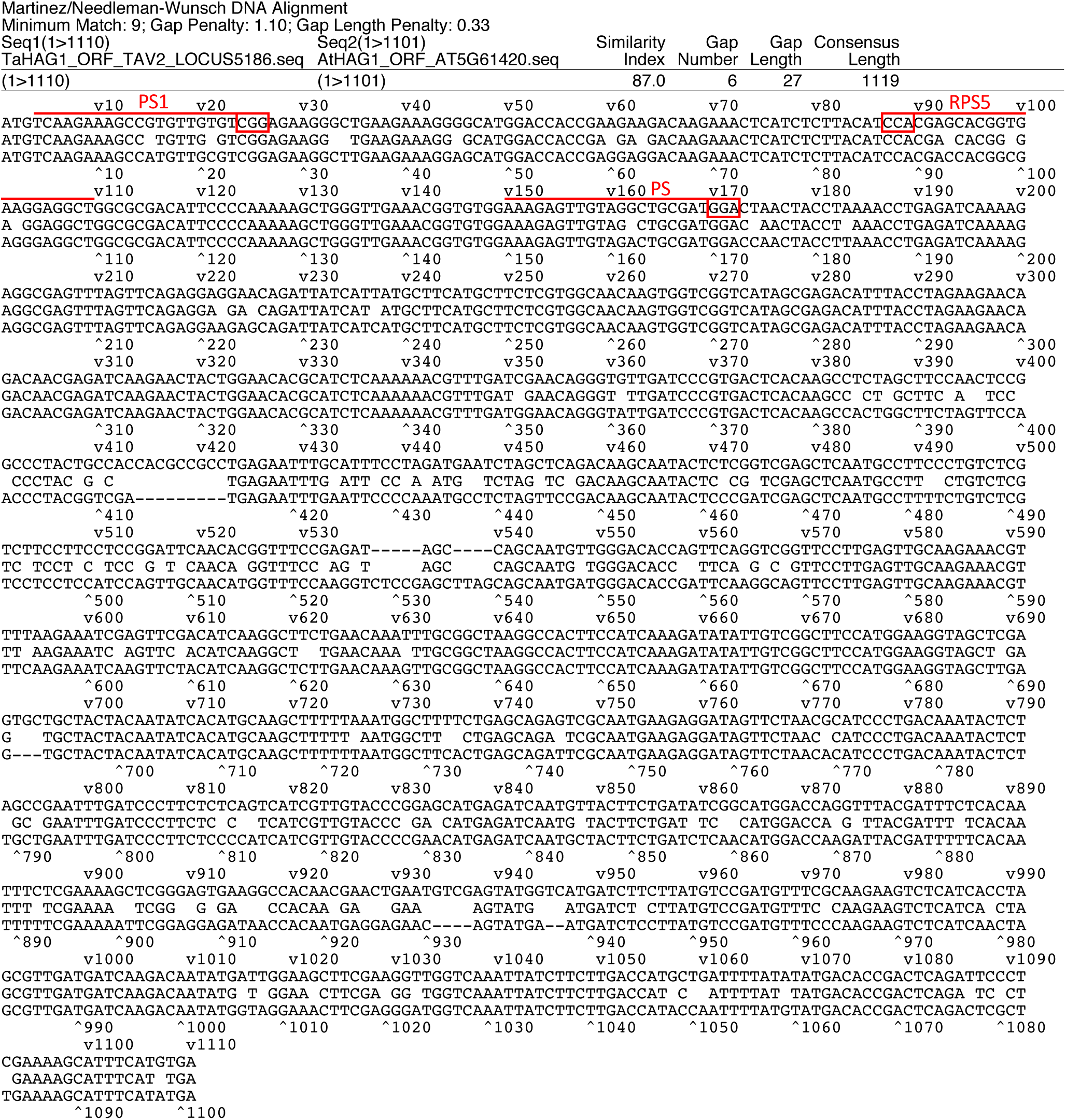
Nucleotide sequence alignment of the pennycress *TaHAG1* (TAV2_LOCUS5186) open reading frame (ORF; top sequence) to that of Arabidopsis *AtHAG1* (AT5G61420; bottom sequence). Letters between the top and bottom sequences denote identity. Locations of the protospacers used for CRISPR gene editing are marked with red lines along with the corresponding PAM sites (boxed in red).

**fig. S21.**
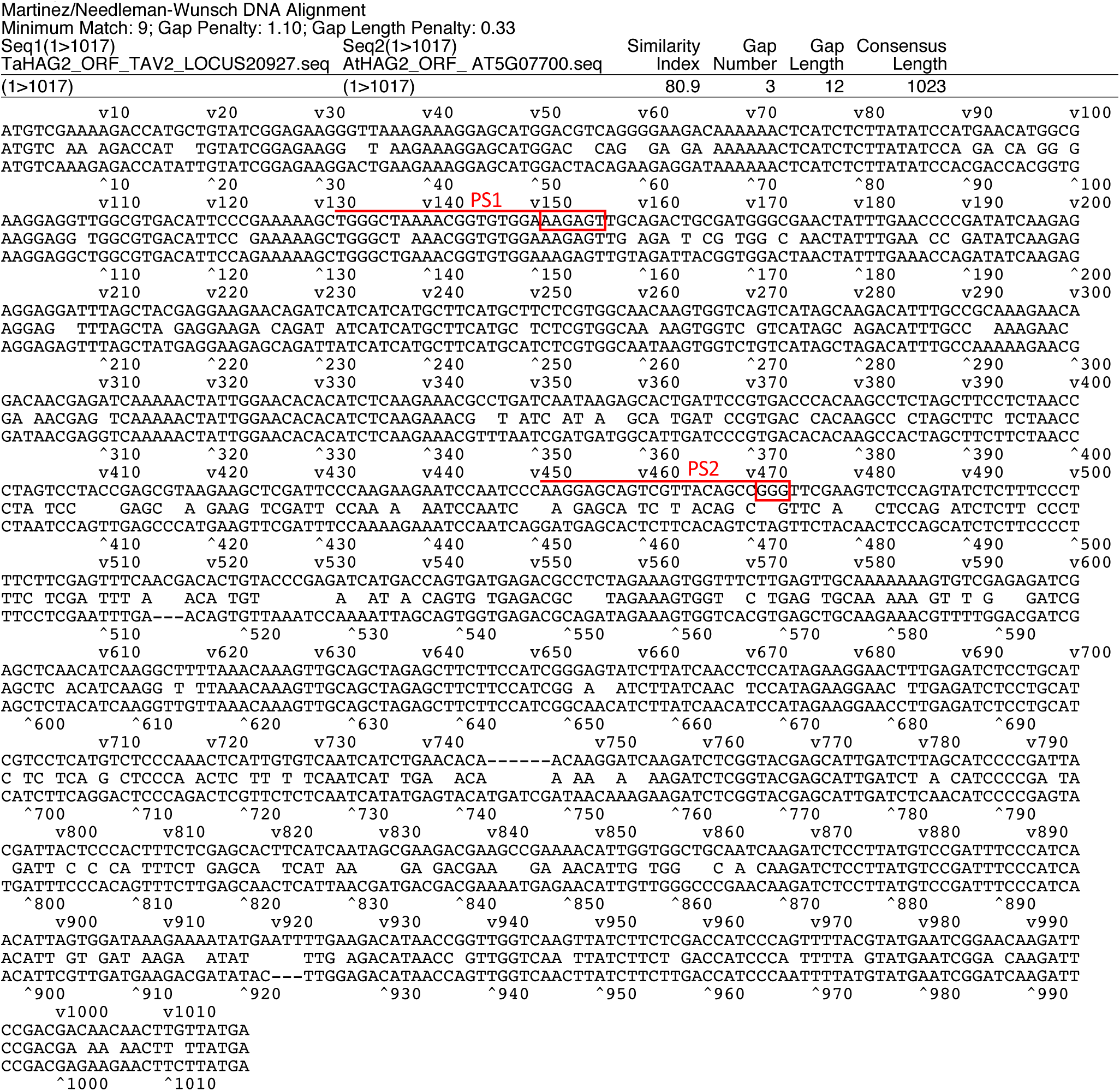
Nucleotide sequence alignment of the pennycress *TaHAG2* (TAV2_LOCUS20927) open reading rame (ORF; top sequence) to that of Arabidopsis *AtHAG2* (AT5G07700; bottom sequence). Letters between the top and bottom sequences denote identity. Locations of the protospacers used for CRISPR gene editing are marked with red lines along with the corresponding PAM sites (boxed in red). Note that PS1 was used with *Staphylococcus aureus* Cas9 (SaCas9) whereas PS2 was used with *Staphylococcus pyogenes* Cas9 (SpCas9).

**fig. S22.**
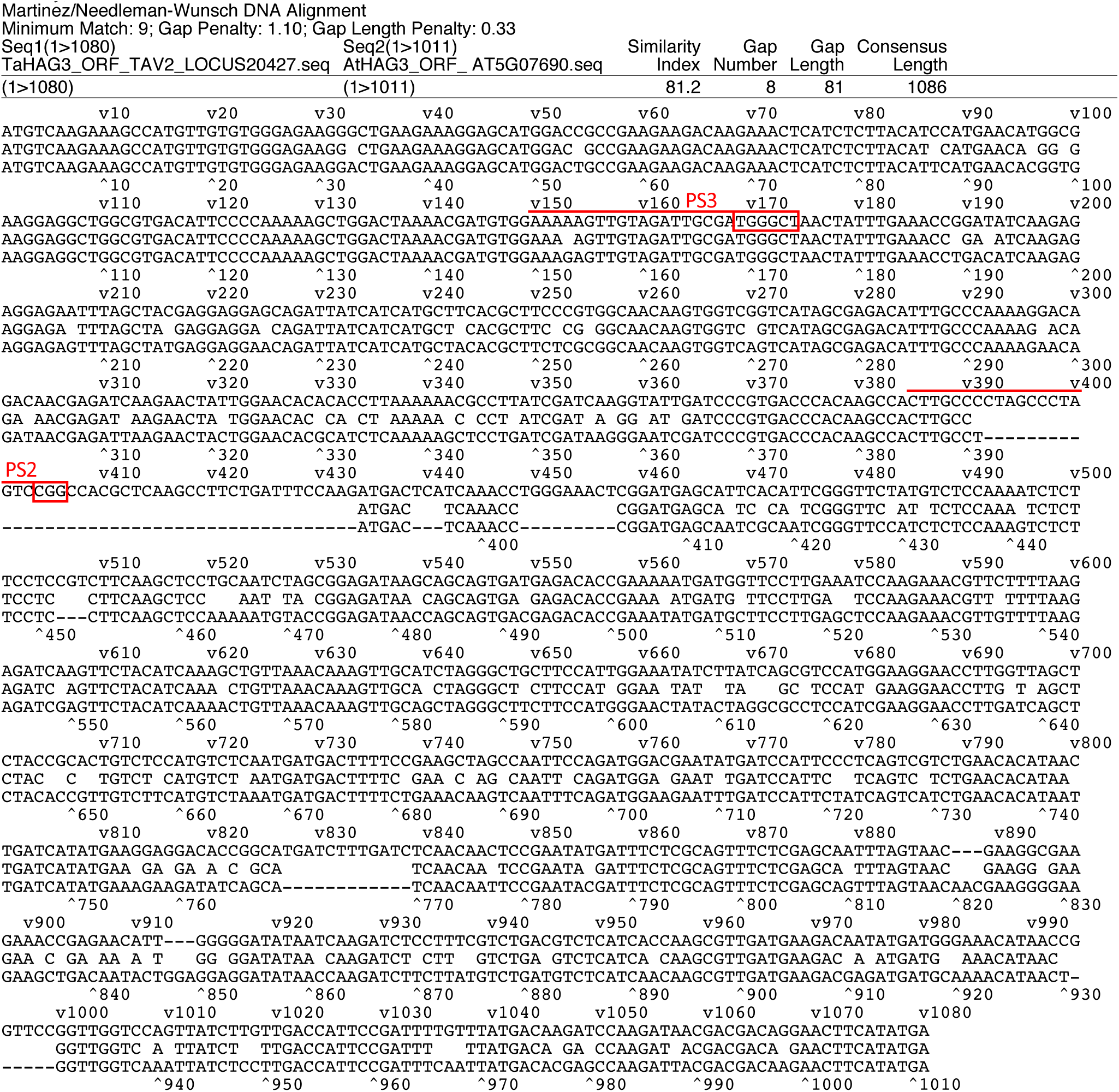
Nucleotide sequence alignment of the pennycress *TaHAG3* (TAV2_LOCUS20427) open reading rame (ORF; top sequence) to that of Arabidopsis *AtHAG3* (AT5G07690; bottom sequence). Letters between the top and bottom sequences denote identity. Locations of the protospacers used for CRISPR gene editing are marked with red lines along with the corresponding PAM sites (boxed in red). Note that PS3 was used with *Staphylococcus aureus* Cas9 (SaCas9) whereas PS2 was used with *Staphylococcus pyogenes* Cas9 (SpCas9).

**fig. S23.**
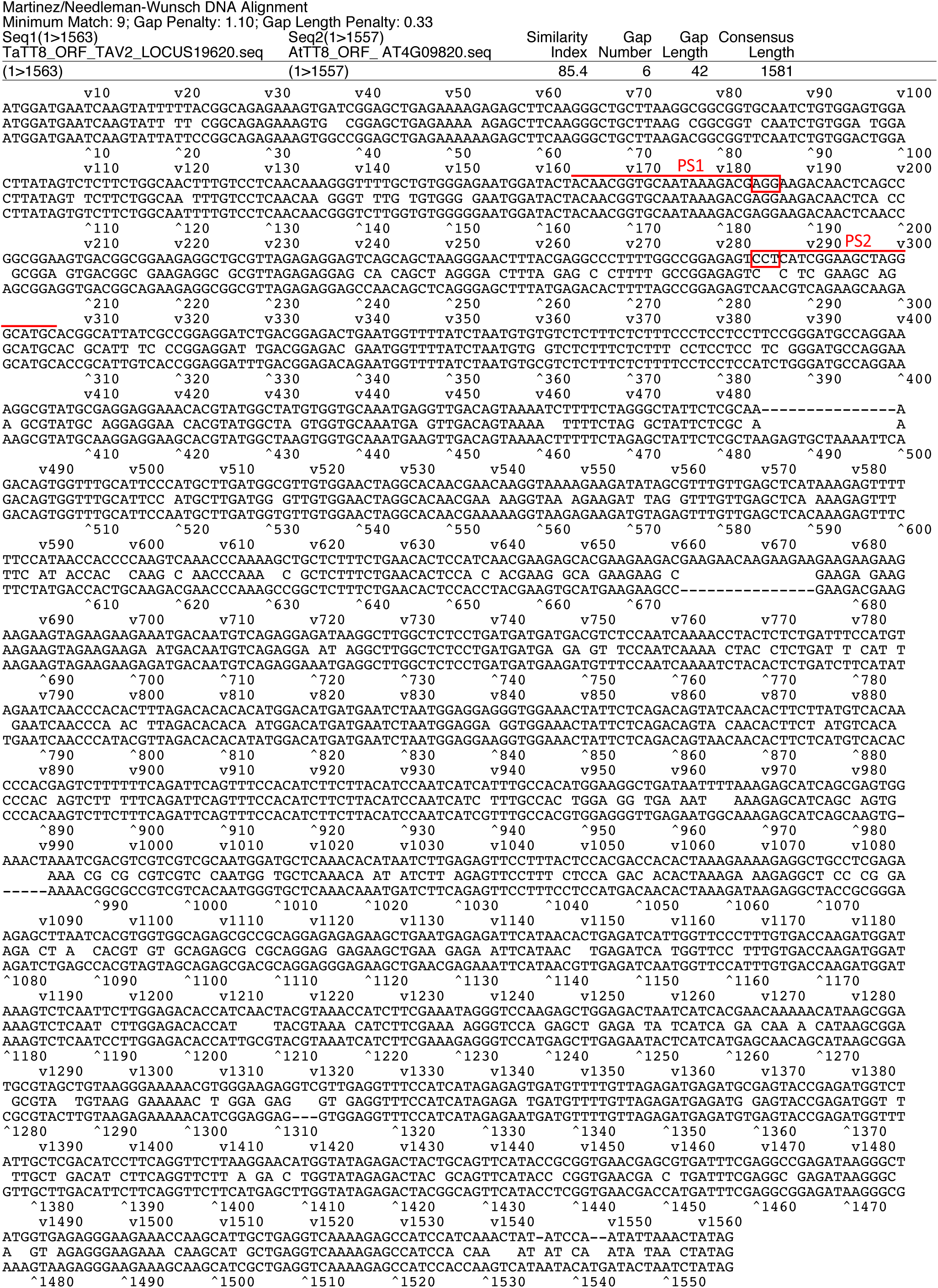
Nucleotide sequence alignment of the pennycress *TaTT8* (TAV2_LOCUS19620) open reading frame (ORF; top sequence) to that of Arabidopsis *AtTT8* (AT4G09820; bottom sequence). Letters between the top and bottom sequences denote identity. Locations of the protospacers used for CRISPR gene editing are marked with red lines along with the corresponding PAM sites (boxed in red).

### Data S1

<u>Pennycress protospacer sequences:</u>

Green Bold: Protospacer sequence locations
Blue: PAM sites (NGG) for *SpCas9*
Blue: PAM sites (NNGRRT) for *SaCas9*

*MYC3* protospacer sequences:

PS1: ACCGACGATAACGCGTCGG
PS2: CCGGGTGCTTTAATCGGGT
PS3: AAATGCTCGAATCAAGATT
PS4: ATGGAAGCGCTCAAGGAAT
PS6: TTCGCCGCGGAACGAACCA (5’ TGGTTCGTTCCGCGGCGAA 3’)

*HAG1* protospacer sequences:

HAG1 CGAGCACGGTGAAGGAGGC
HAG1 AAAGAGTTGTAGGCTGCGA
HAG1 (PS1) TCAAGAAAGCCGTGTTGTGT
HAG1 (RPS5) AGCCTCCTTCACCGTGCTCG

*HAG2* protospacer sequences:

HAG2 (PS1) GACCATGCTGTATCGGAGAA
HAG2 (PS2) AAGGAGCAGTCGTTACAGCC
HAG2 (PS4) TGAACCCCGATATCAAGAG
HAG2 (PS5) CACAACAAGGATCAAGATCT
HAG2 (PS6) TCGGAGAAGGGTTAAAGAA

*HAG3* protospacer sequences:

HAG3 CTTGCCCCTAGCCCTAGTC
HAG3 CTGGGCTAAAACGGTGTGGA
HAG3 (PS2) ACTTGCCCCTAGCCCTAGTC
HAG3 (PS3) GAAAAAGTTGTAGATTGCGA
HAG3 (PS4) GCATCTAGGGCTGCTTCCAT
HAG3 (PS5) ATCTTATCAGCGTCCATGGA
HAG3 (PS6) GAAGCTAGCCAATTCCAGA

*TT8* protospacer sequences:

TT8 (PS1) ACAACGGTGCAATAAAGACG

#### >Wild-Type *MYC3* ORF

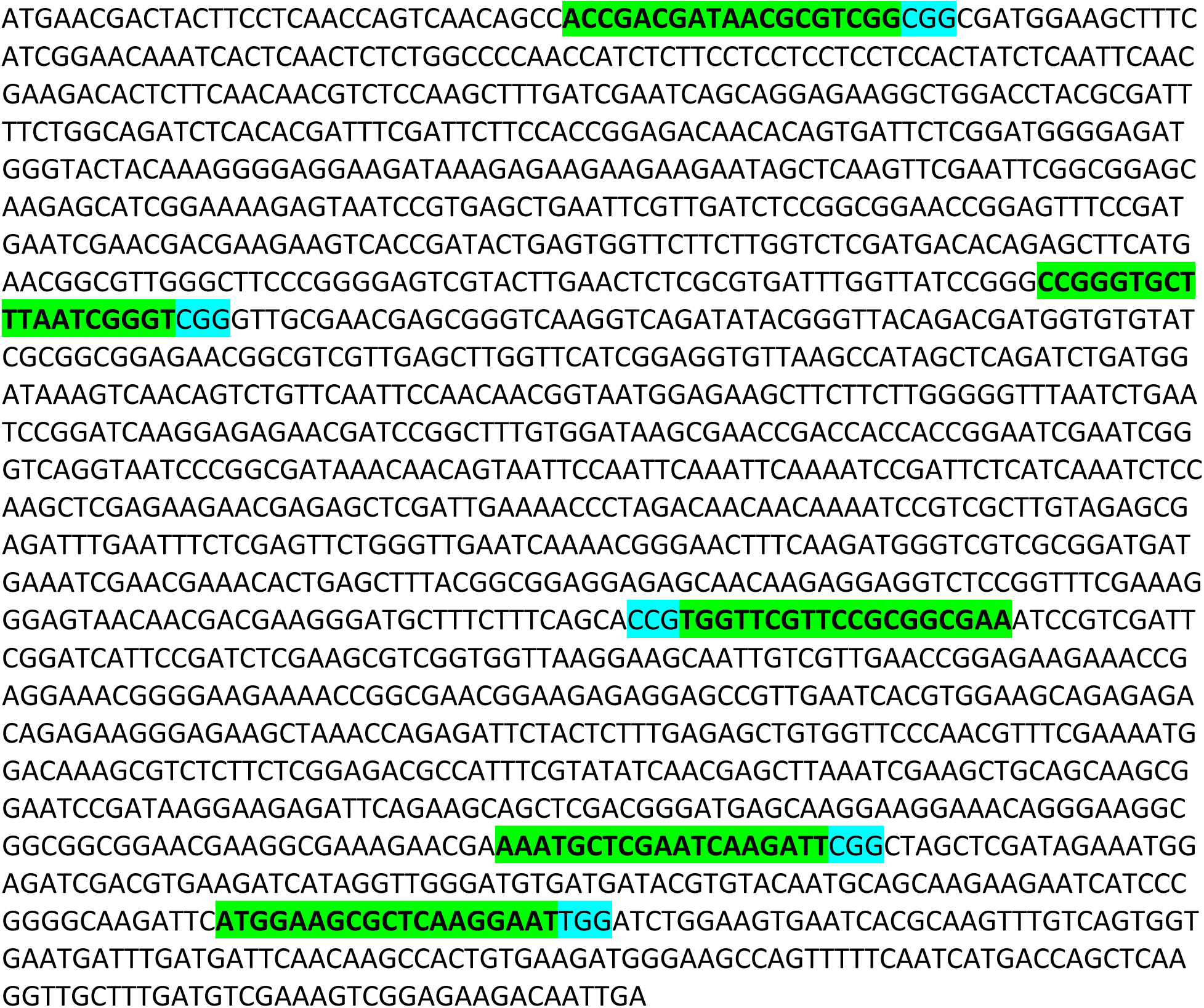

#### >Wild-Type *MYC3* amino acid sequence

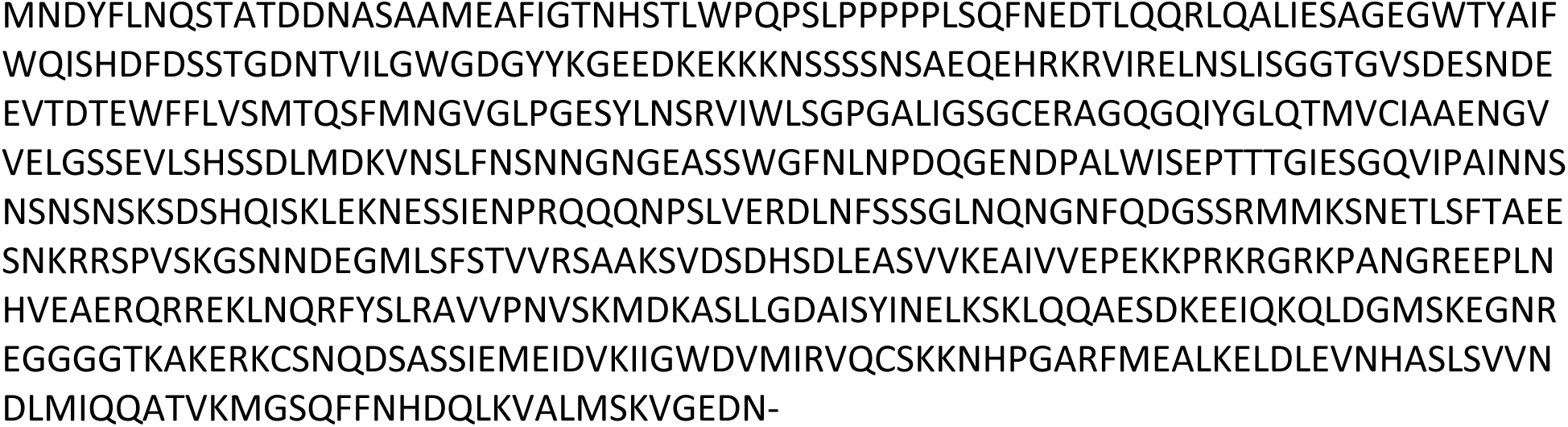

#### >*myc3-1* (+A generated with PS6) (Insertion red bold underlined)

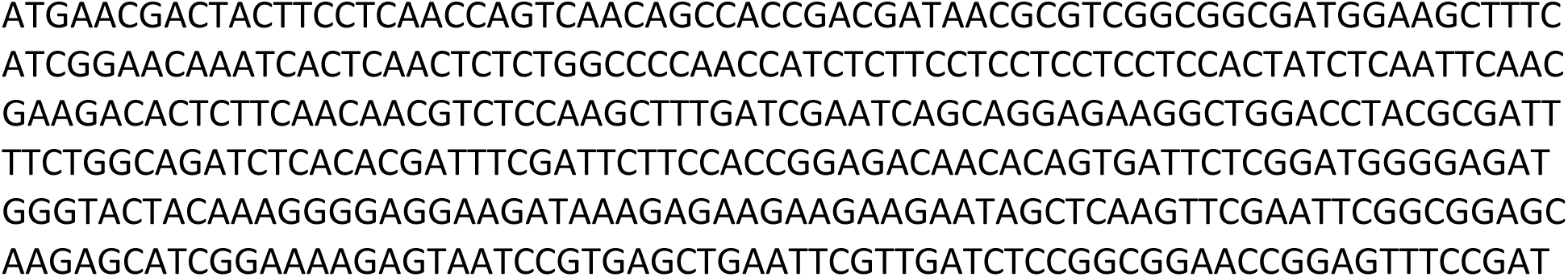
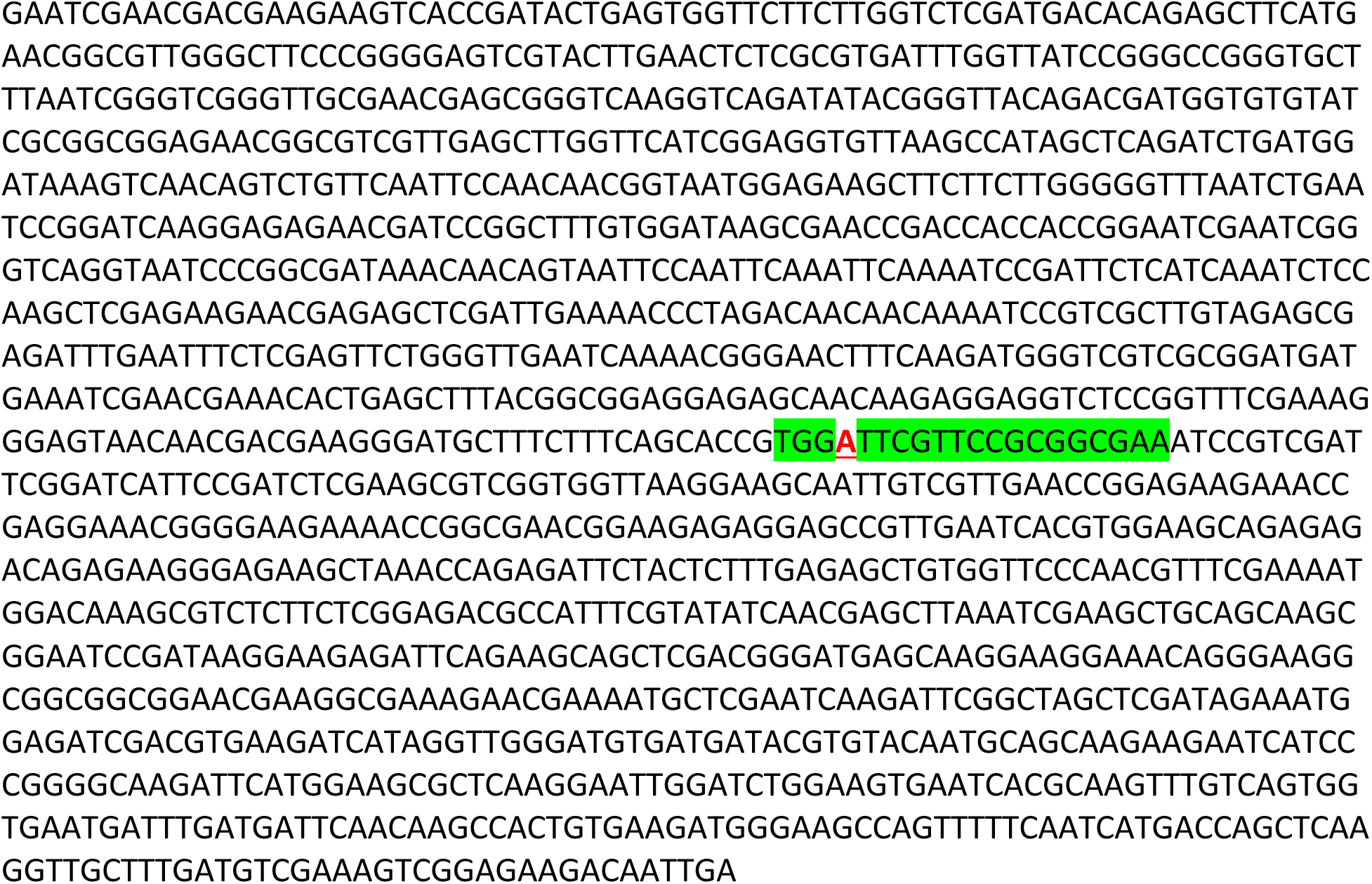

#### >*myc3-1* predicted amino acid sequence

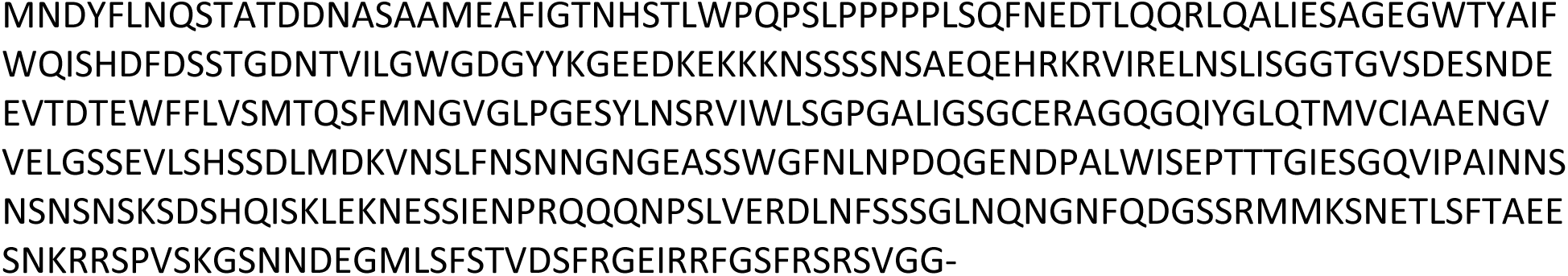

#### >*myc3-2* (−20bp generated with PS6, −41bp generated with PS4) (deletion red bold strikethrough)

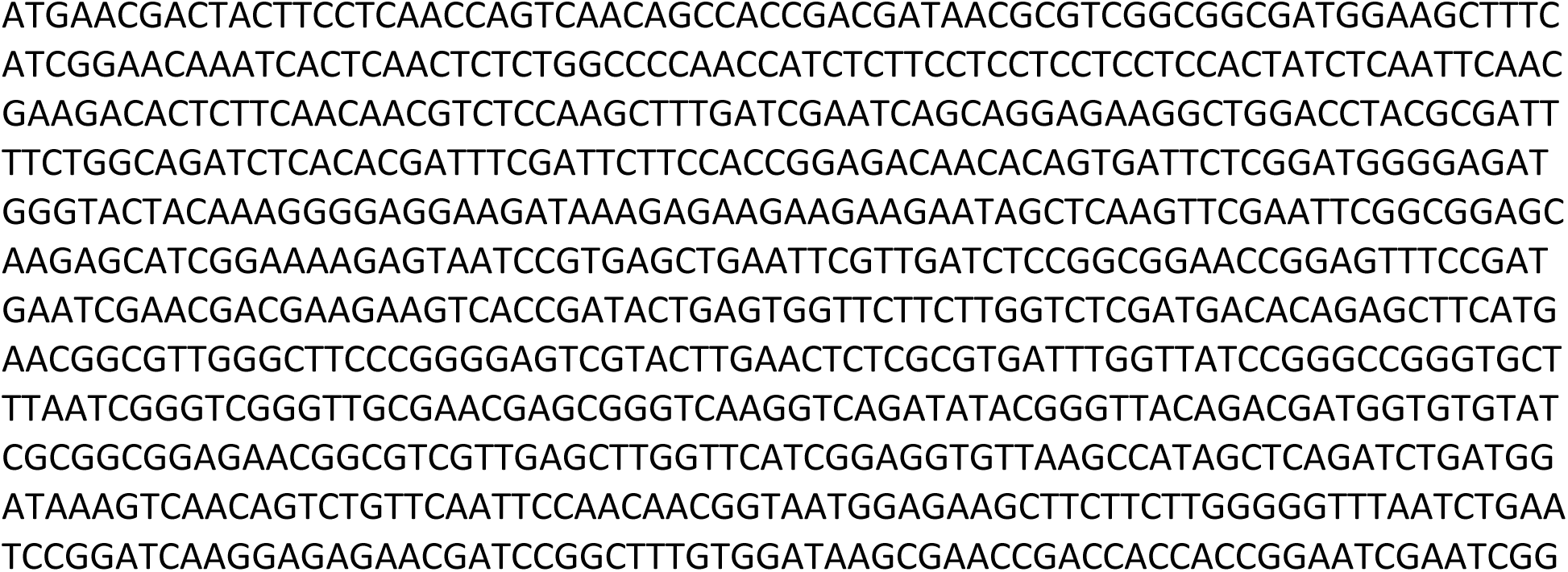
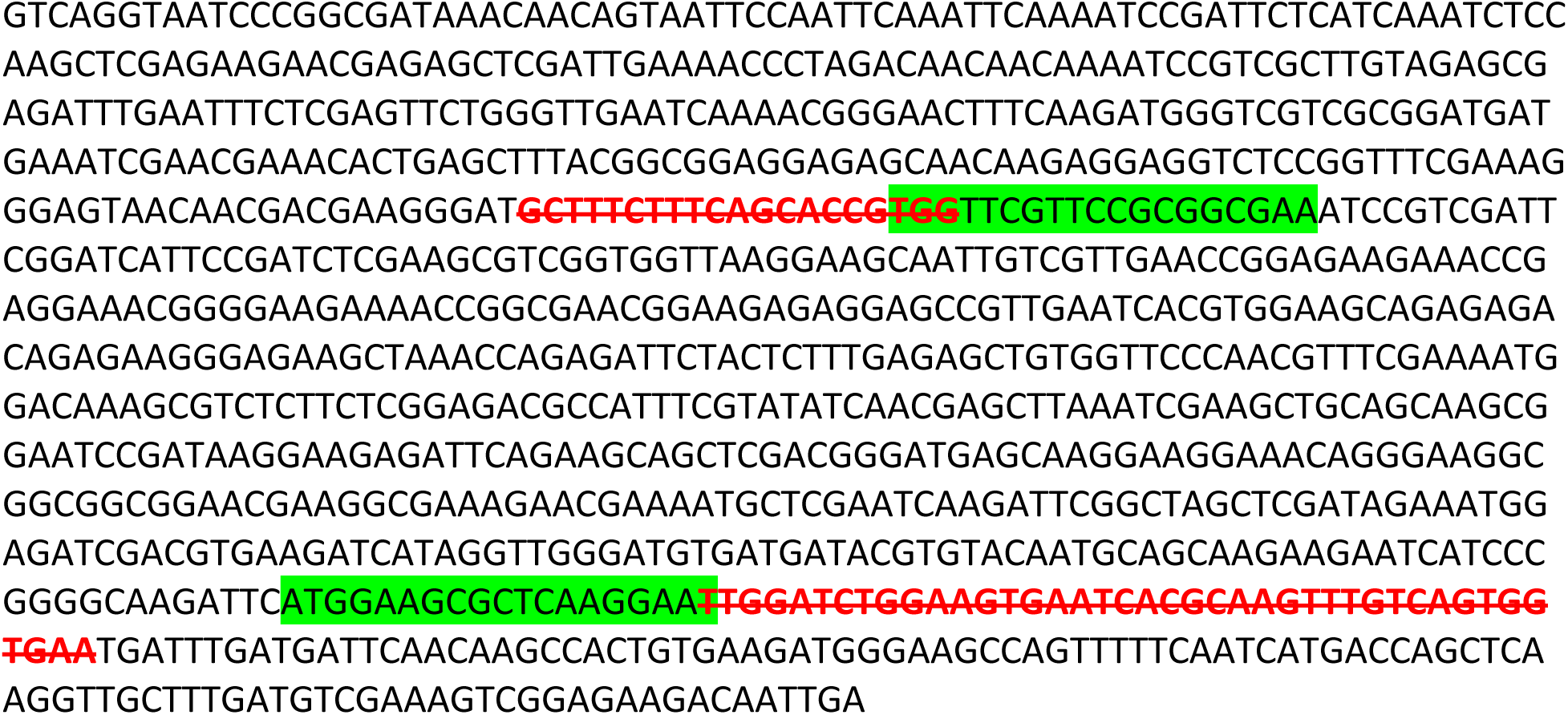

#### >*myc3-2* predicted amino acid sequence

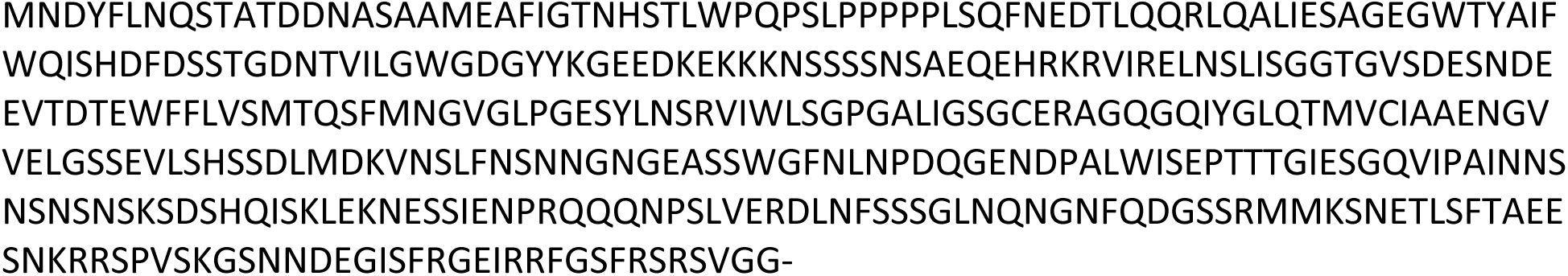

#### >*myc3-3* (inversion of 529bp between PS6 and PS4) (Inversion bold orange)

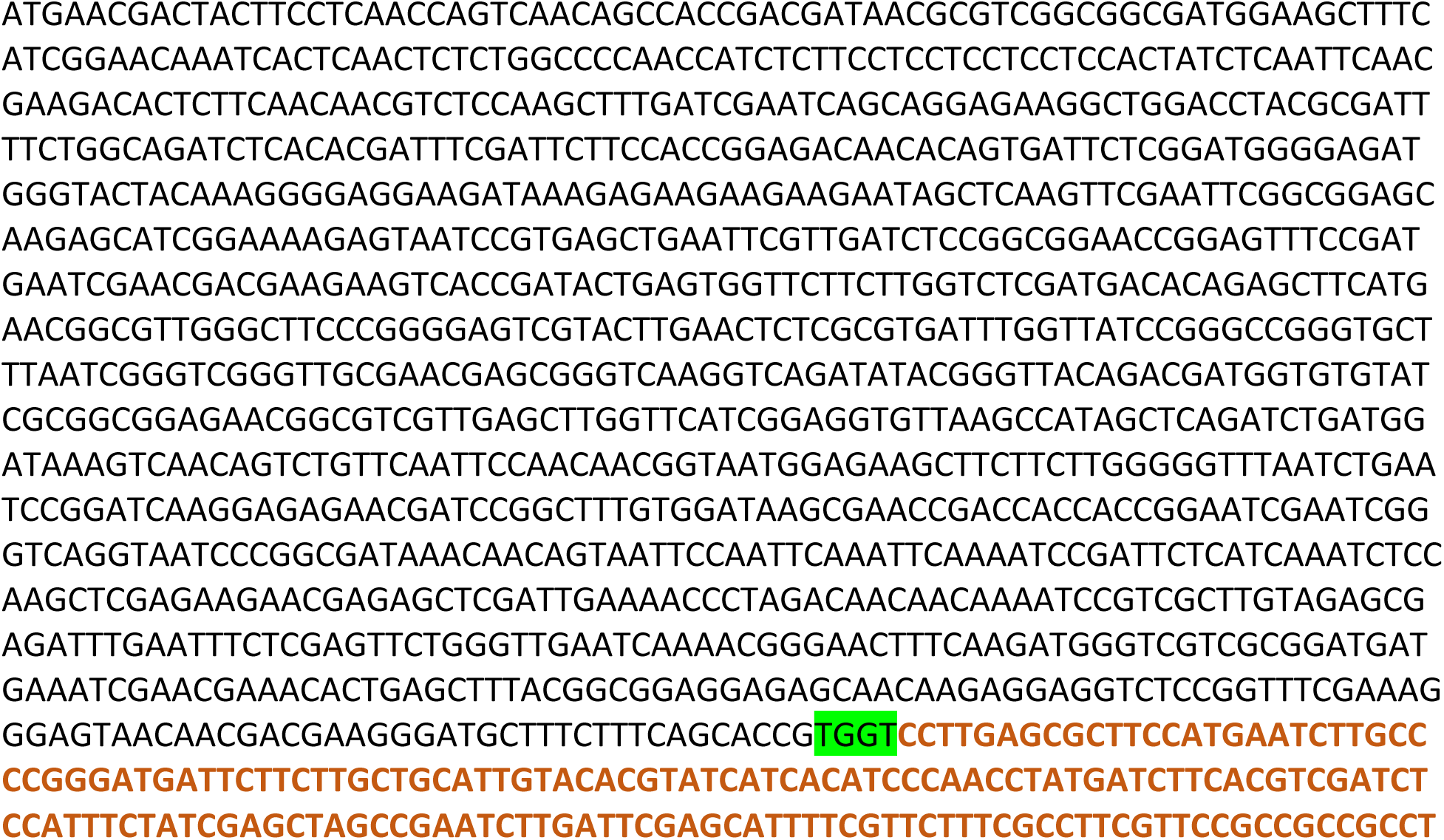
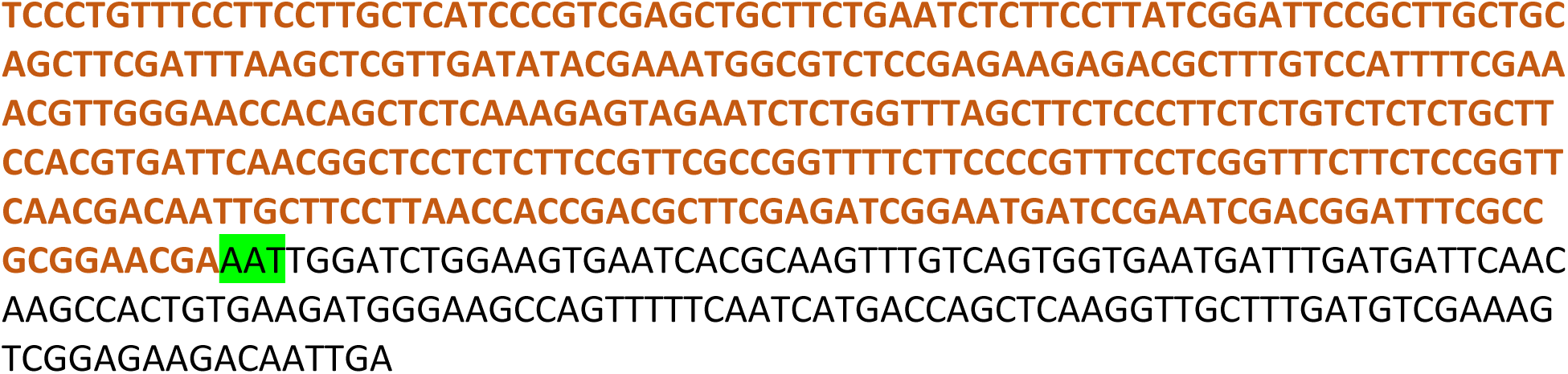

#### >*myc3-3* predicted amino acid sequence

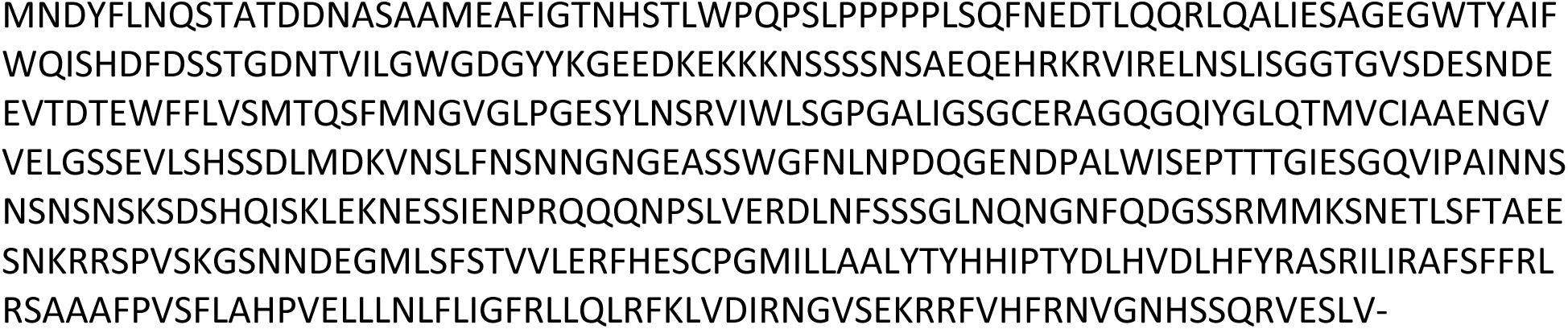

#### >*myc3* allele in triple mutant line (*myc3tt8fae1*)−1 (−17bp generated with PS2, −144bp between PS3 and PS4) (red bold strikethrough)

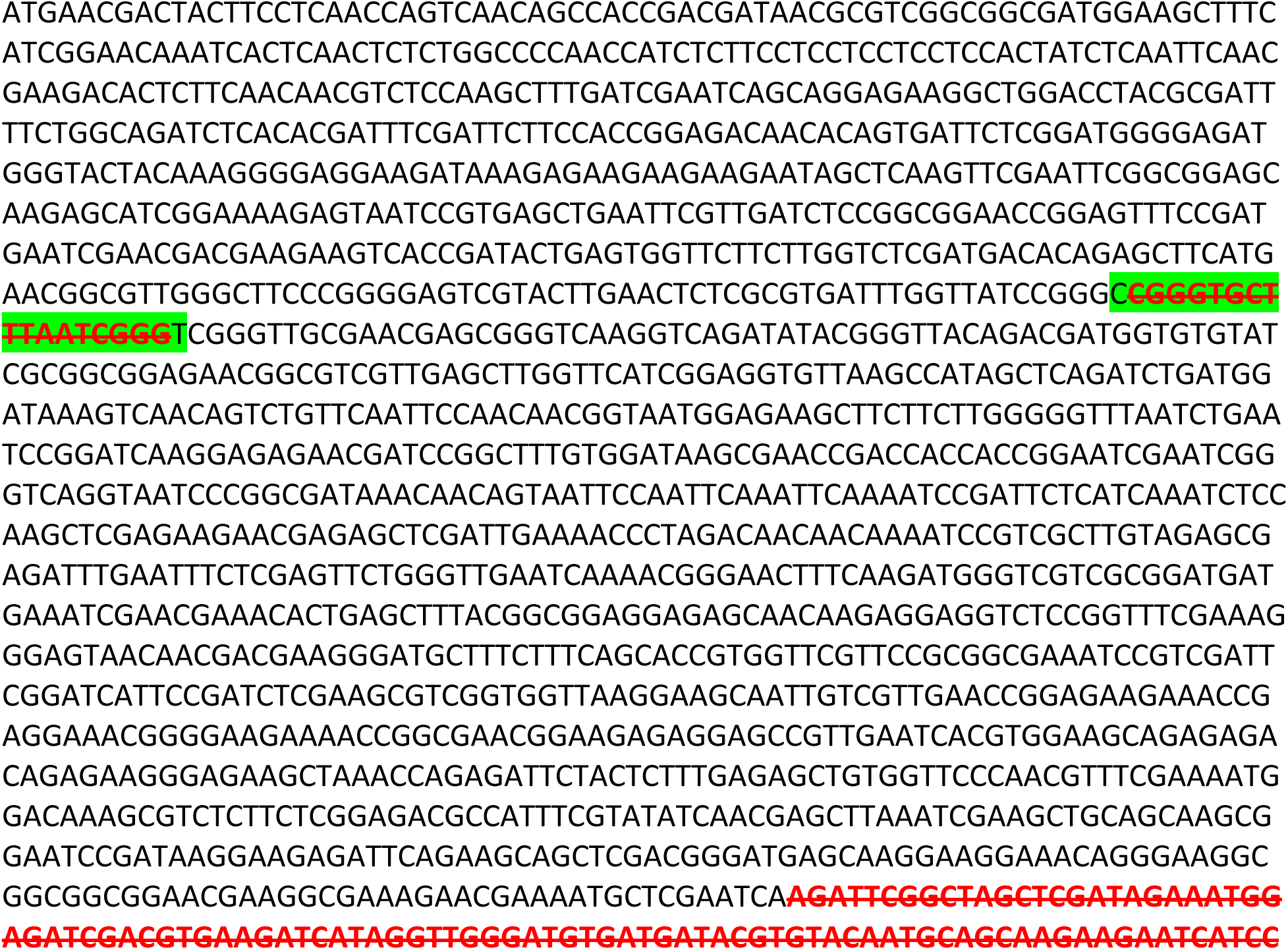
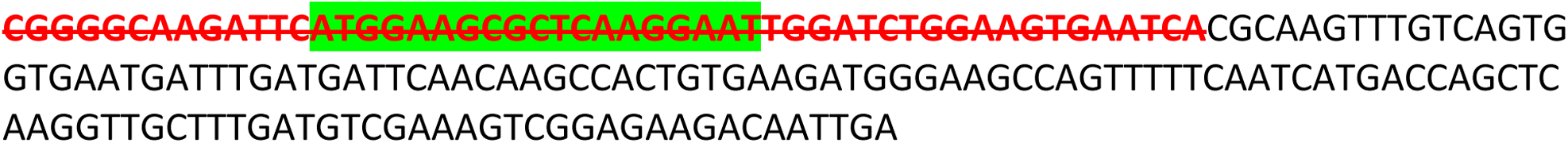

#### >*myc3* in triple mutant line (*myc3tt8fae1*)−1 predicted amino acid sequence

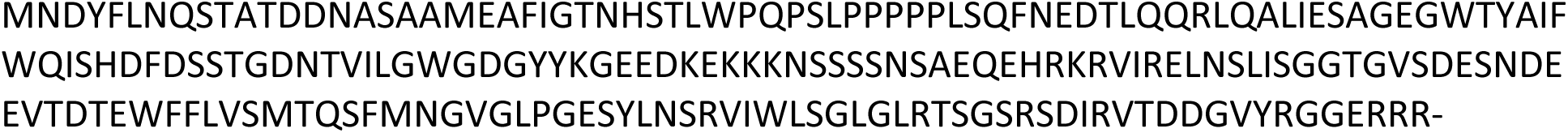

#### >*myc3* allele in triple mutant line (*myc3tt8fae1*)−2 (−4bp in PS1, −120bp in PS34) (red bold strikethrough)

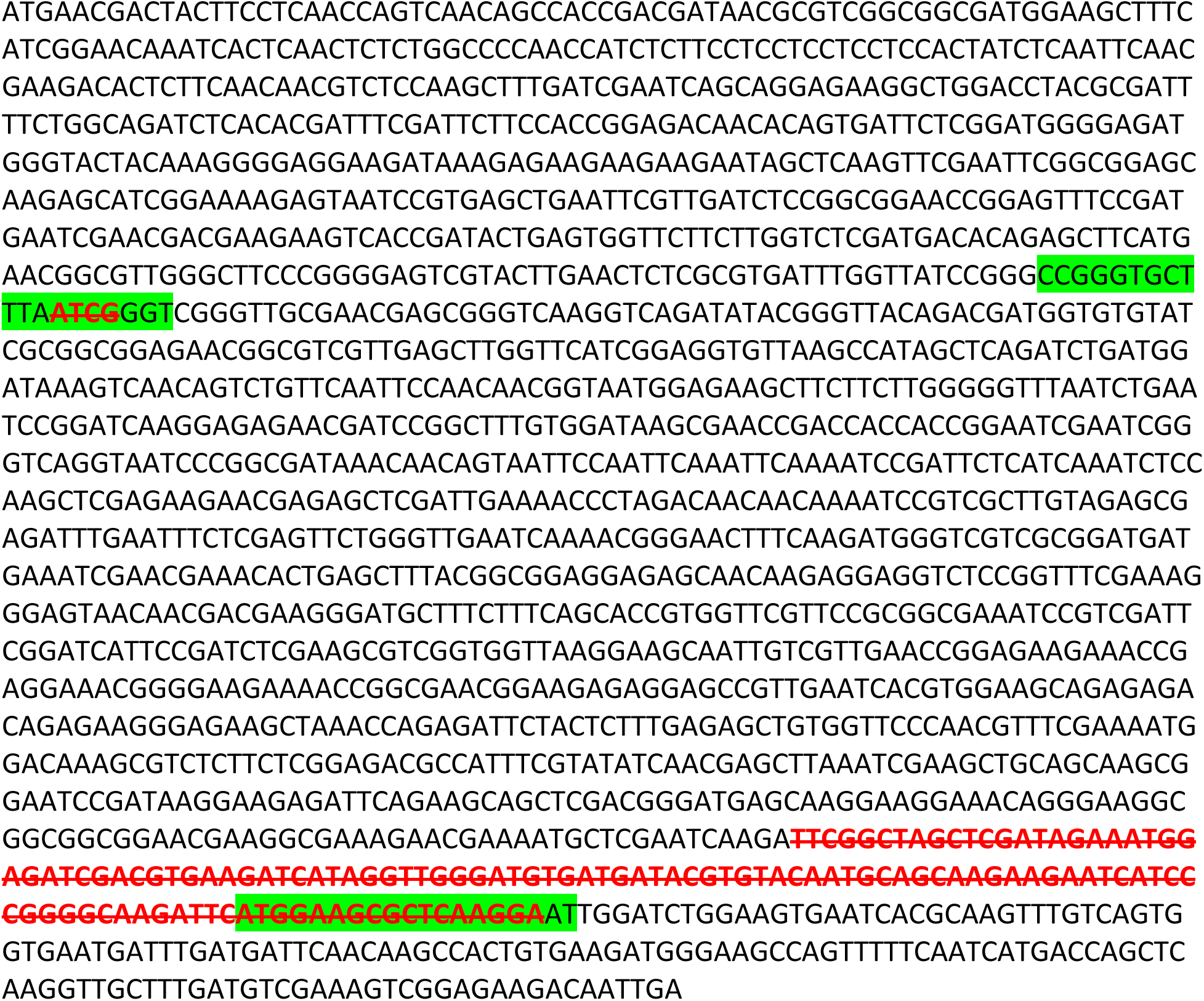

#### >*myc3* allele in triple mutant line (*myc3tt8fae1*)−2 predicted protein sequence

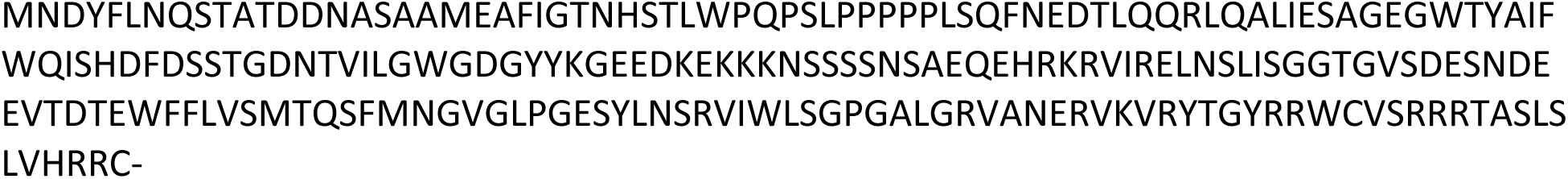

#### >Wild-Type *HAG1* ORF

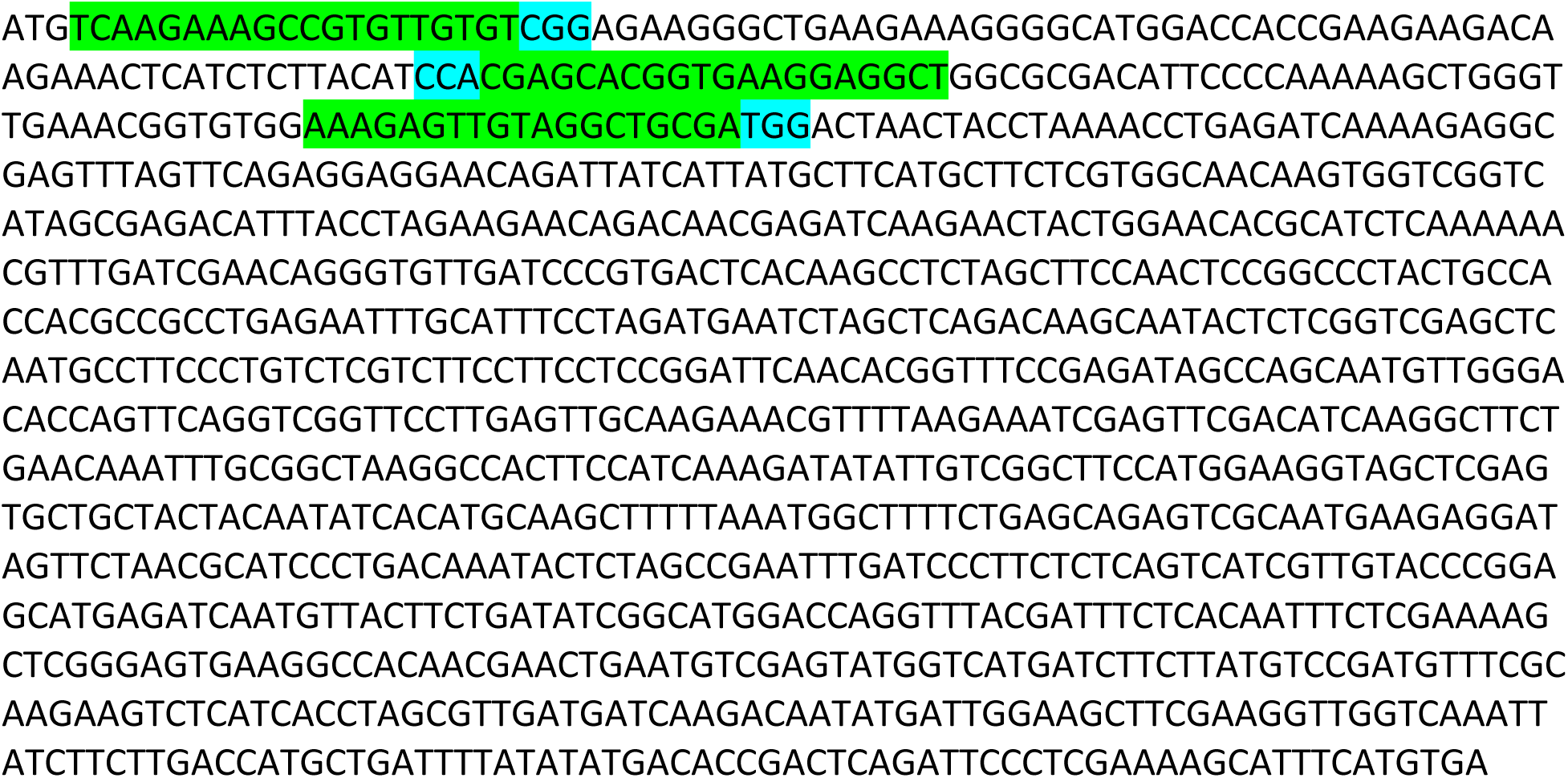

#### >Wild-Type *HAG1* amino acid sequence

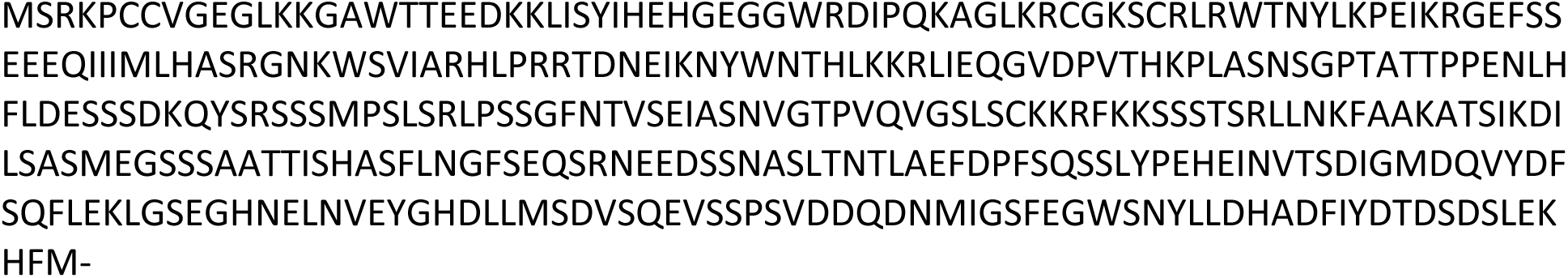

#### >*hag1-1* (−4bp) (red bold strikethrough)

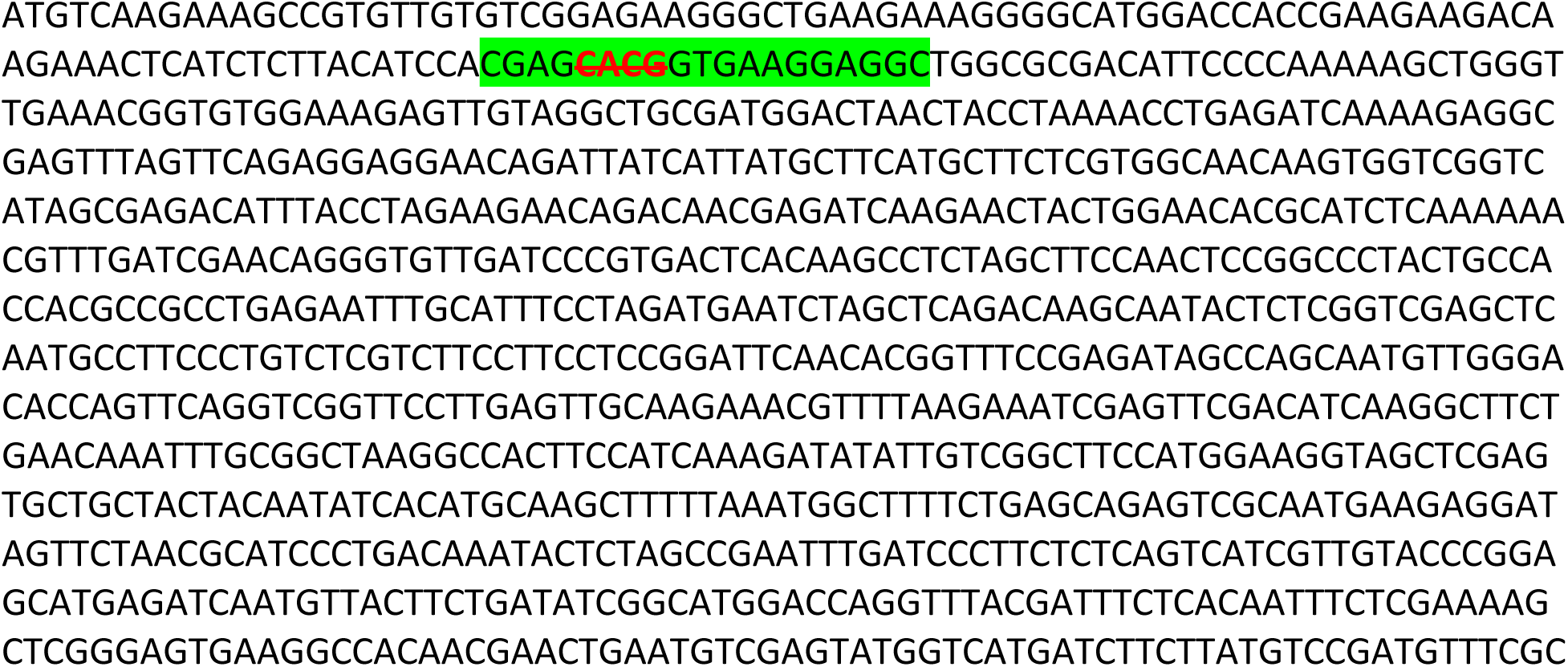
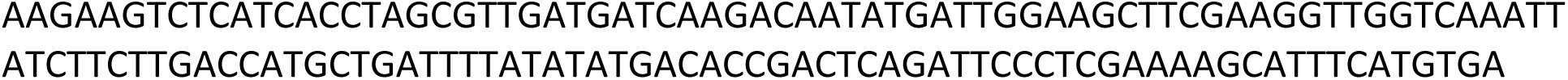

#### >*hag1-1* (−4bp) predicted amino acid sequence

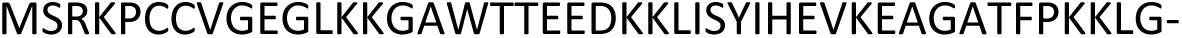

#### >*hag1* (−G) (red bold strikethrough) in *hag1* single and *hag1hag2* (−G; +T) double mutant

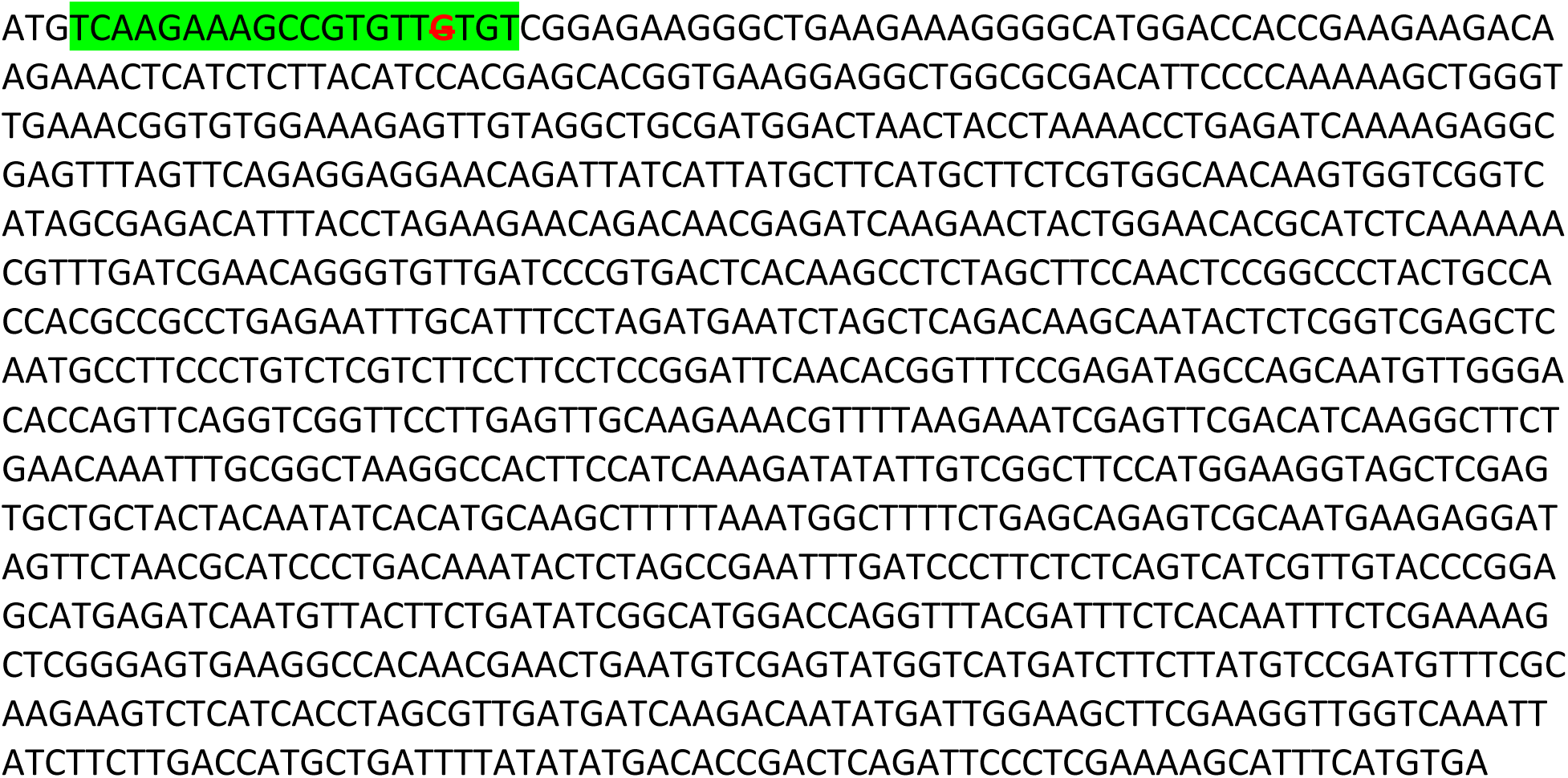

#### >*hag1* (−G) predicted amino acid sequence in *hag1* single and *hag1hag2* (−G; +T) double mutant

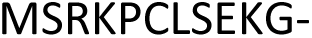

#### >*hag1* (+A) (red bold underlined) in *hag1hag3* (+A; −13bp) double mutant

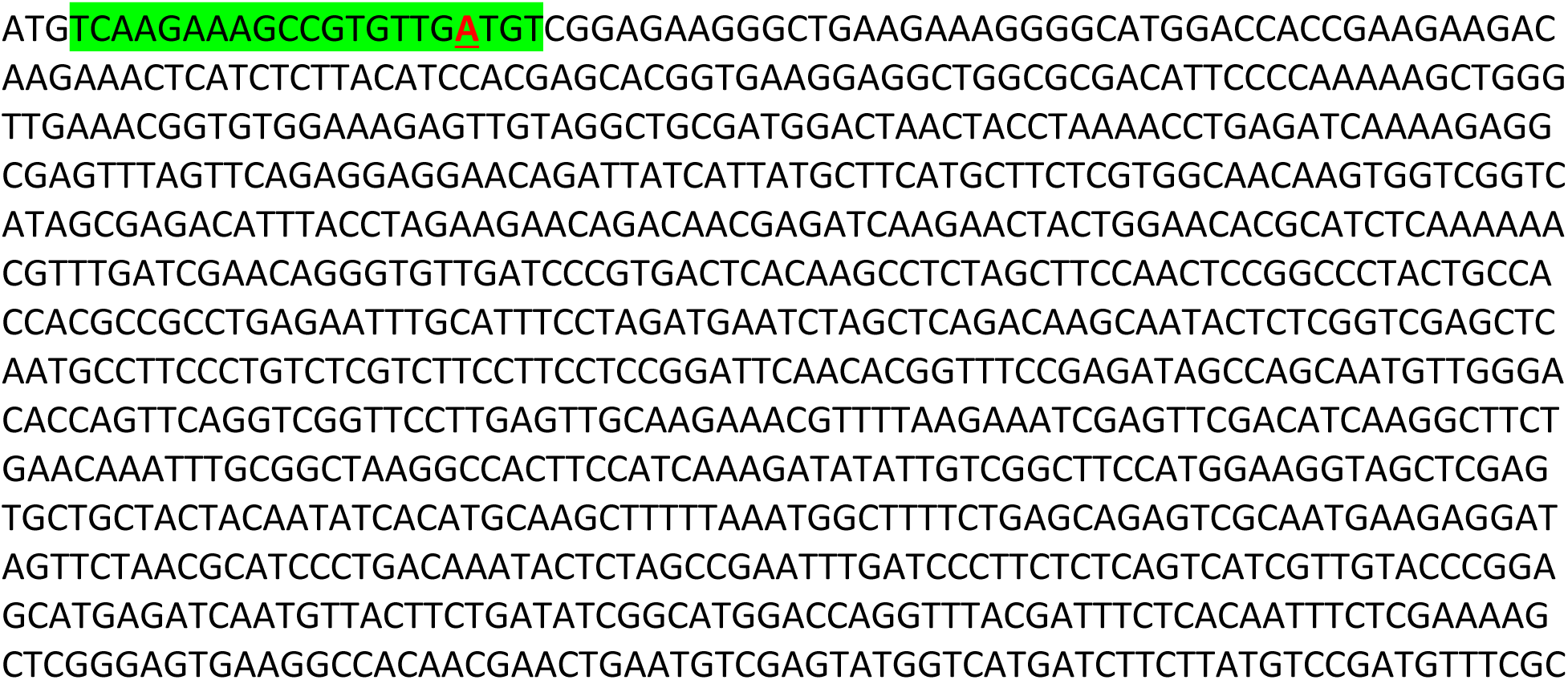
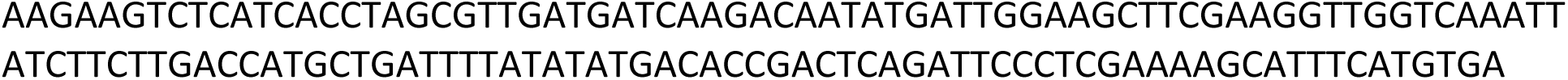

#### >*hag1* (+A) predicted amino acid sequence in *hag1hag3* (+A; −13bp) double mutant

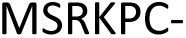

#### >*hag1* (+G) (red bold underlined) in *hag1hag2hag3* (+G; −9bp; −36bp) triple mutant

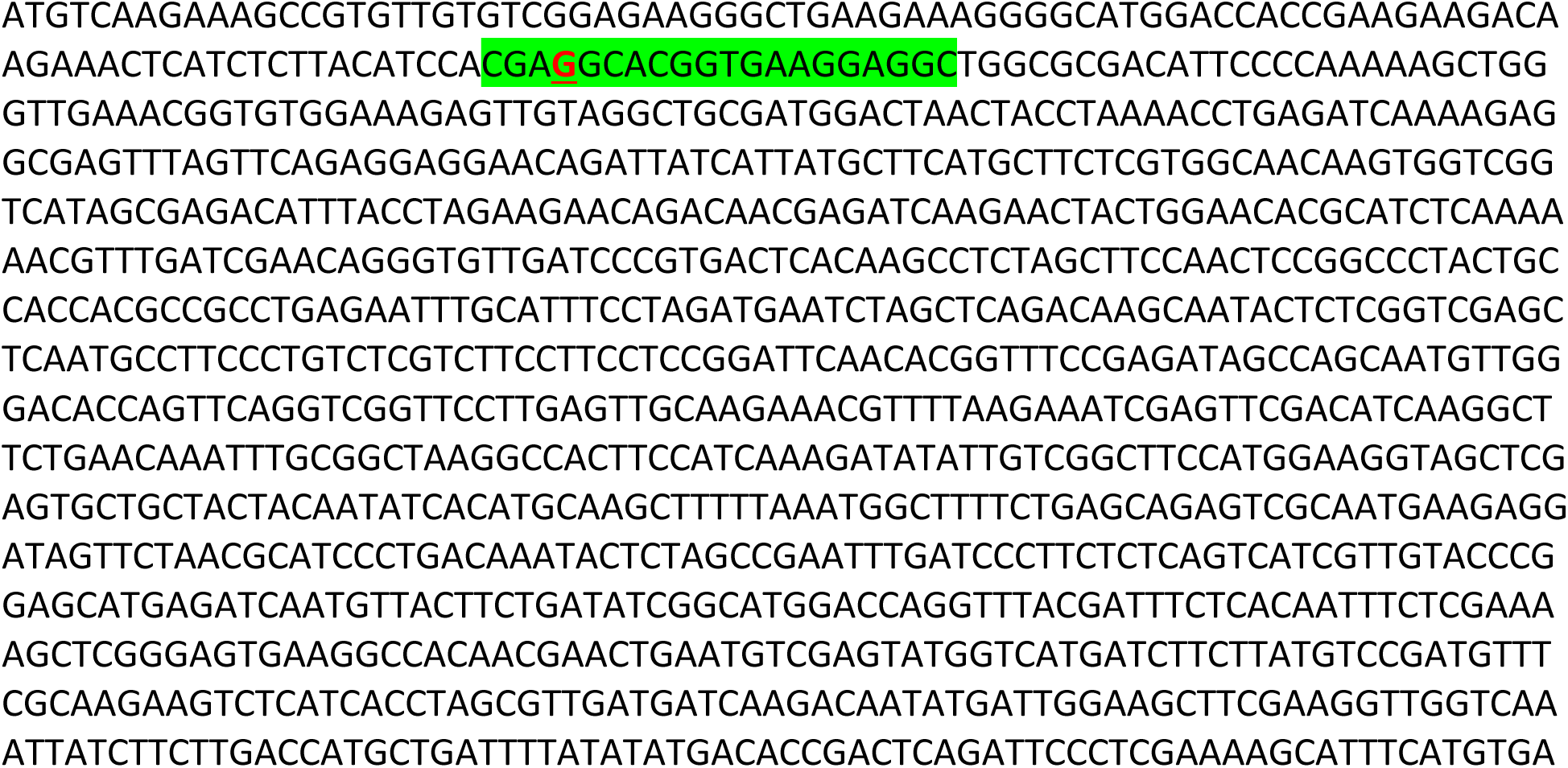

#### >*hag1* (+G) predicted amino acid sequence in *hag1hag2hag3* (+G; −9bp; −36bp) triple mutant

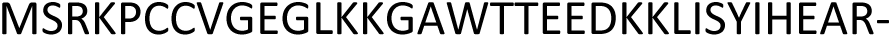

#### >*hag1* (−8bp) (red bold strikethrough) in *hag1hag2hag3* (−8bp; −3bp; +C) triple mutant

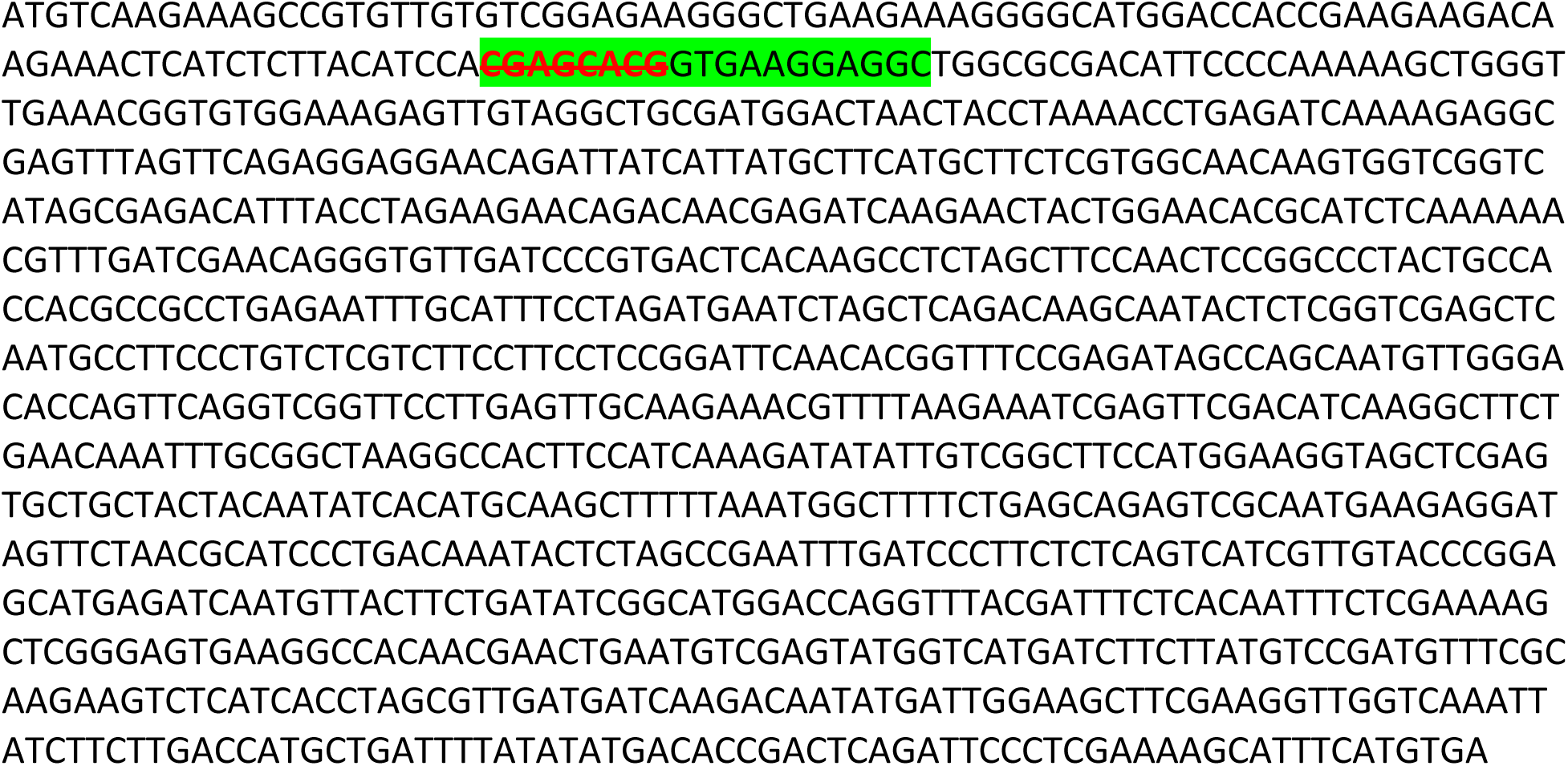

#### >*hag1* (−8bp) predicted amino acid sequence in *hag1hag2hag3* (−8bp; −3bp; +C) triple mutant

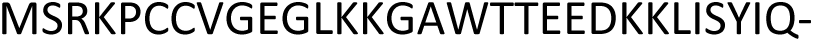

#### >Wild-Type *HAG2* ORF

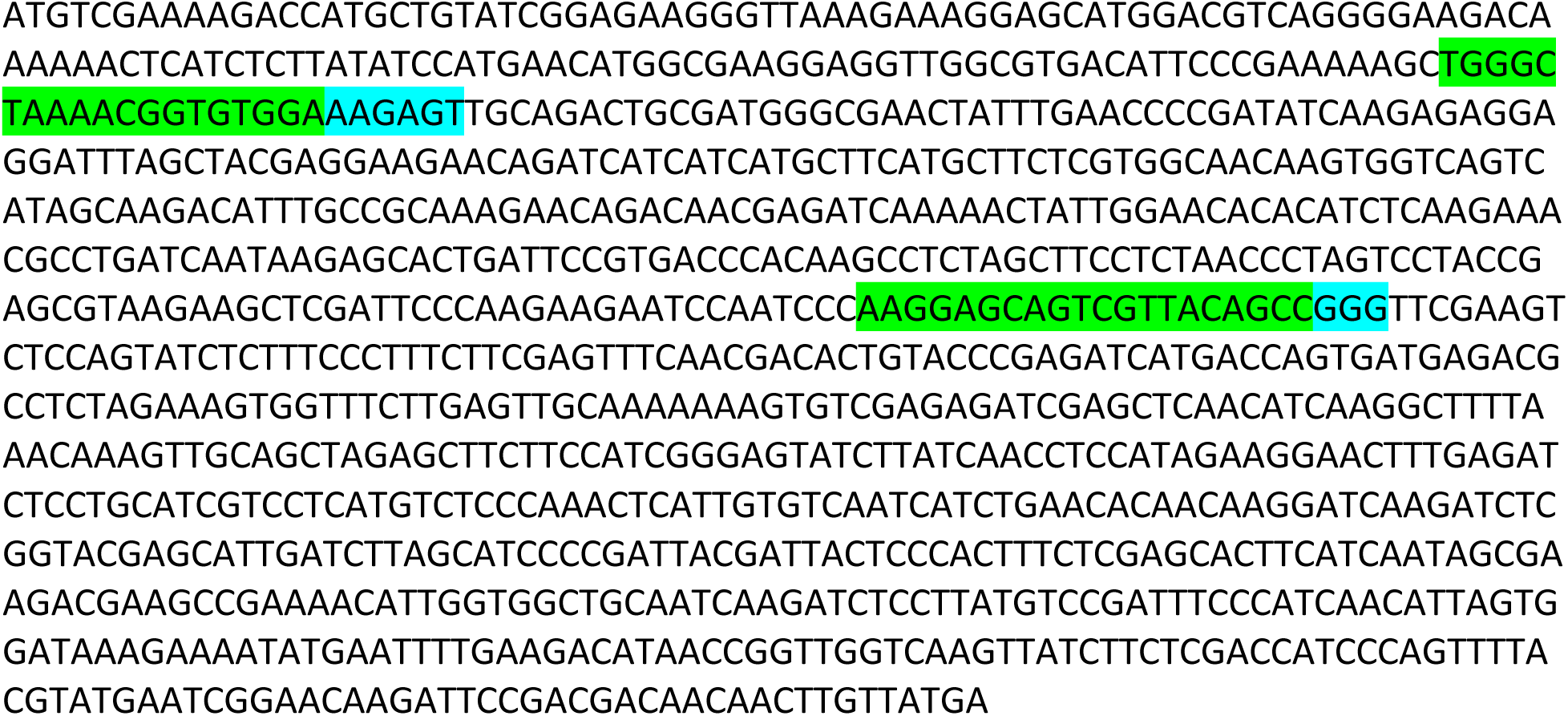

#### >Wild-Type *HAG2* amino acid sequence

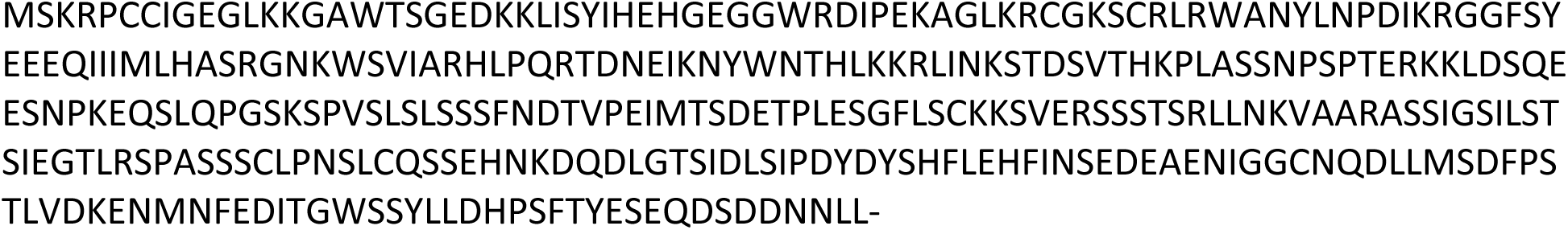

#### >*hag2* (+T) (bold red underlined) in *hag2* single and *hag1hag2* (−G; +T) double mutant

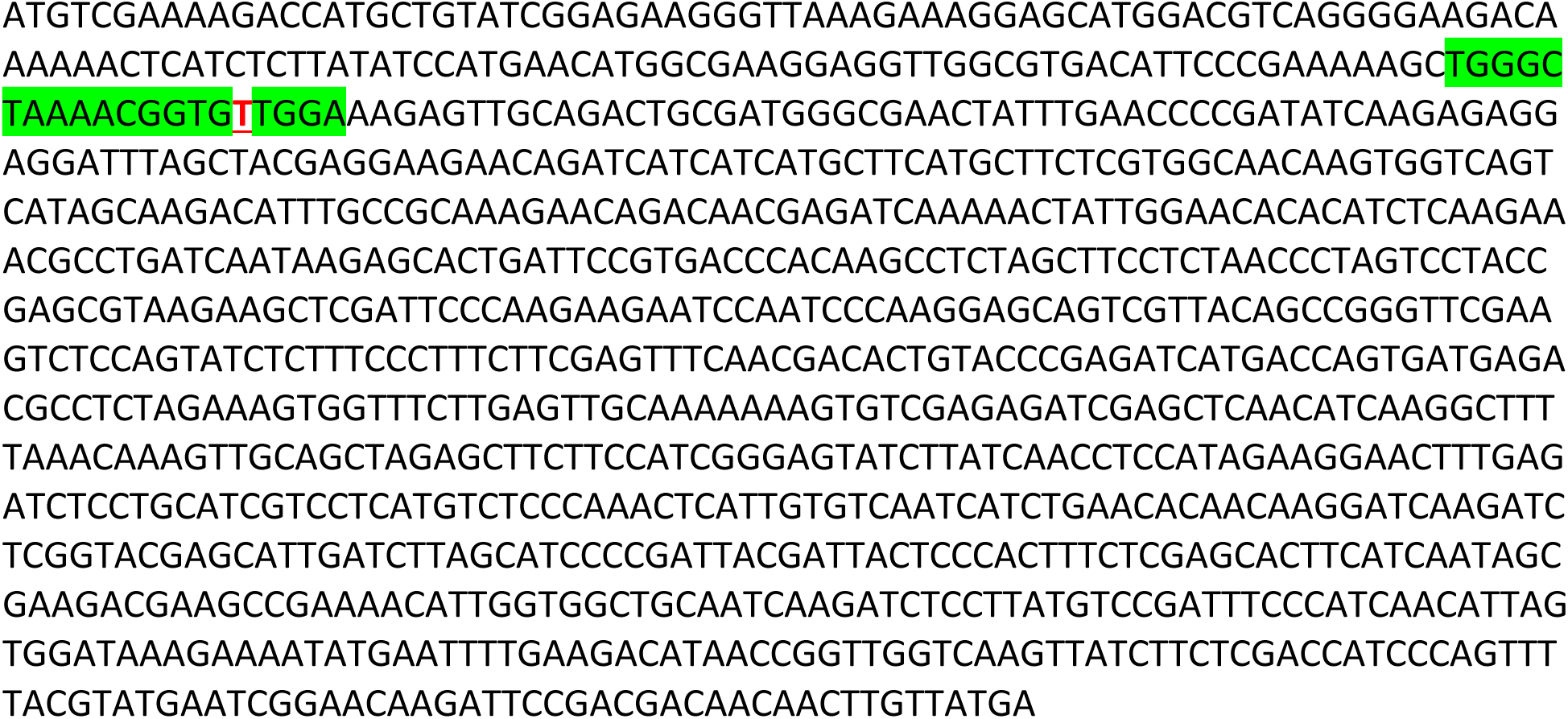

#### >*hag2* (+T) predicted amino acid sequence in *hag1hag2* (−G; +T) double mutant

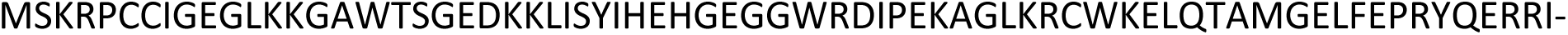

#### >*hag2* (−9bp) (red bold strikethrough) in *hag1hag2hag3* (+G; −9bp; −36bp) triple mutant

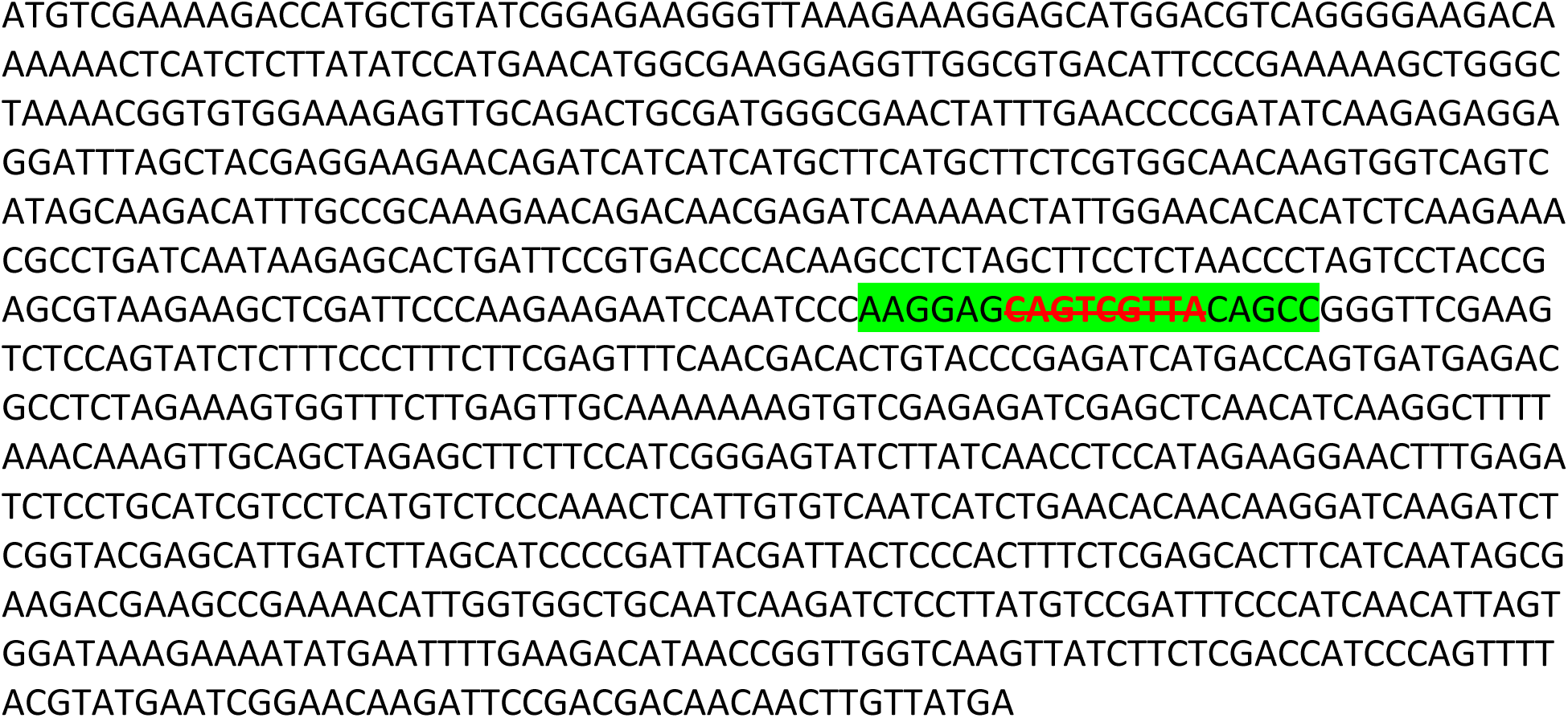

#### >*hag2* (−9bp) predicted amino acid sequence in *hag1hag2hag3* (+G; −9bp; −36bp) triple mutant

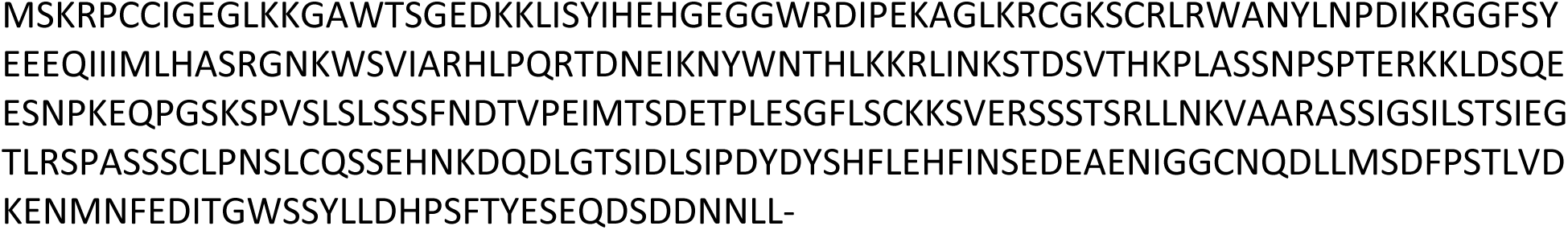

#### >*hag2* (−3bp) (red bold strikethrough) in *hag1hag2hag3* (−8bp; −3bp; +C) triple mutant

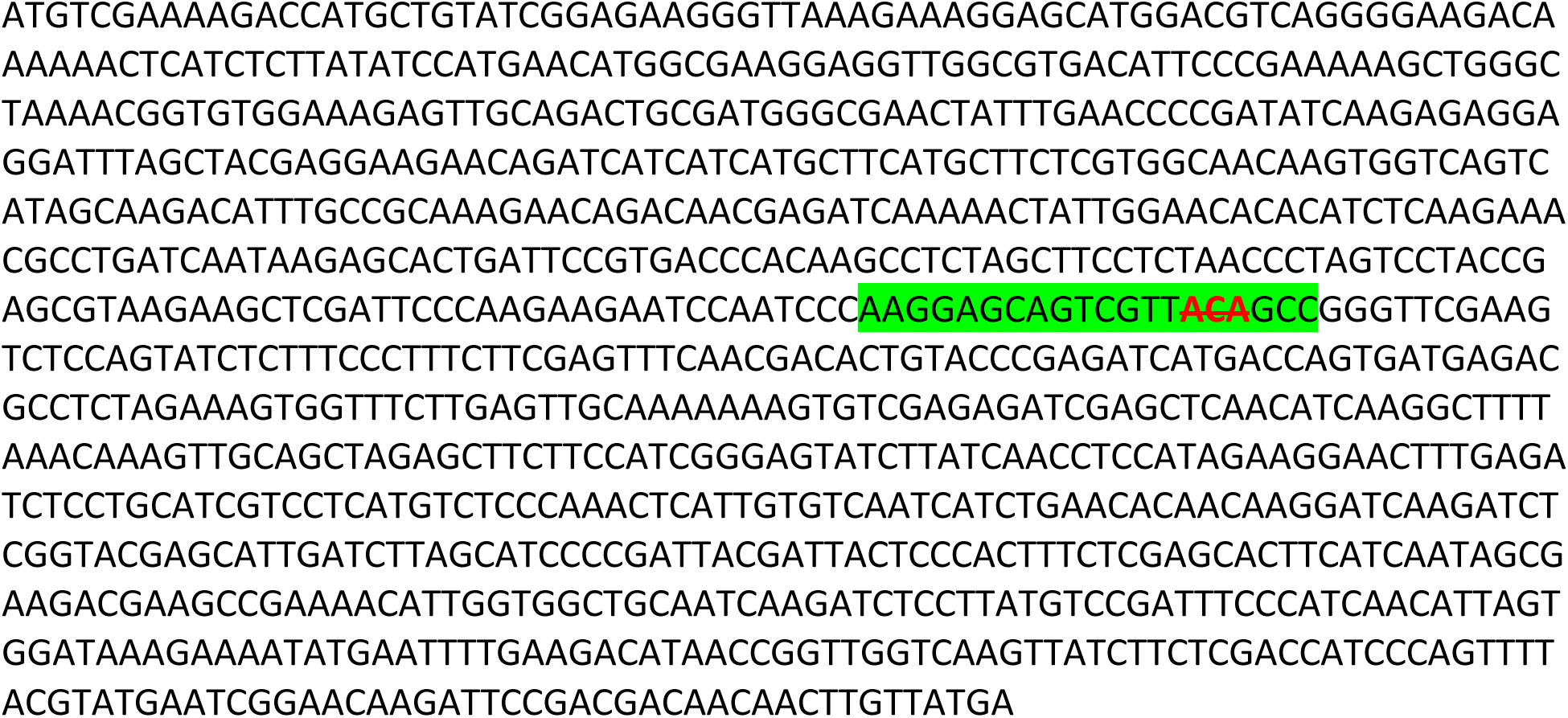

#### >*hag2* (−3bp) predicted amino acid sequence in *hag1hag2hag3* (−8bp; −3bp; +C) triple mutant

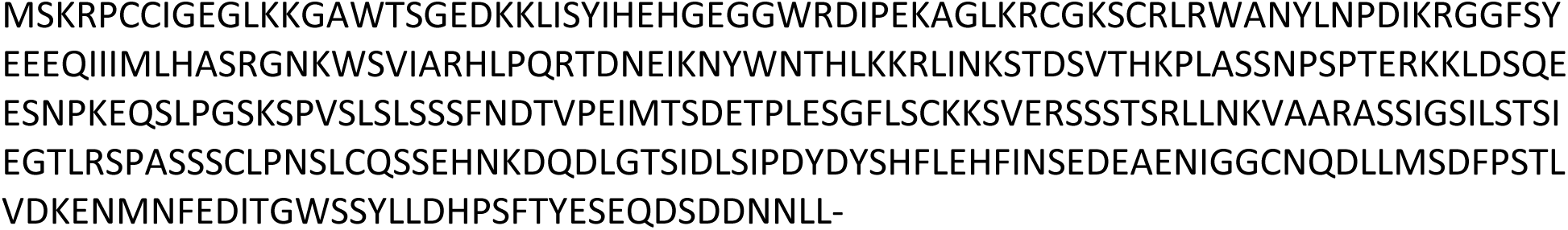

#### >Wild-Type *HAG3 ORF*

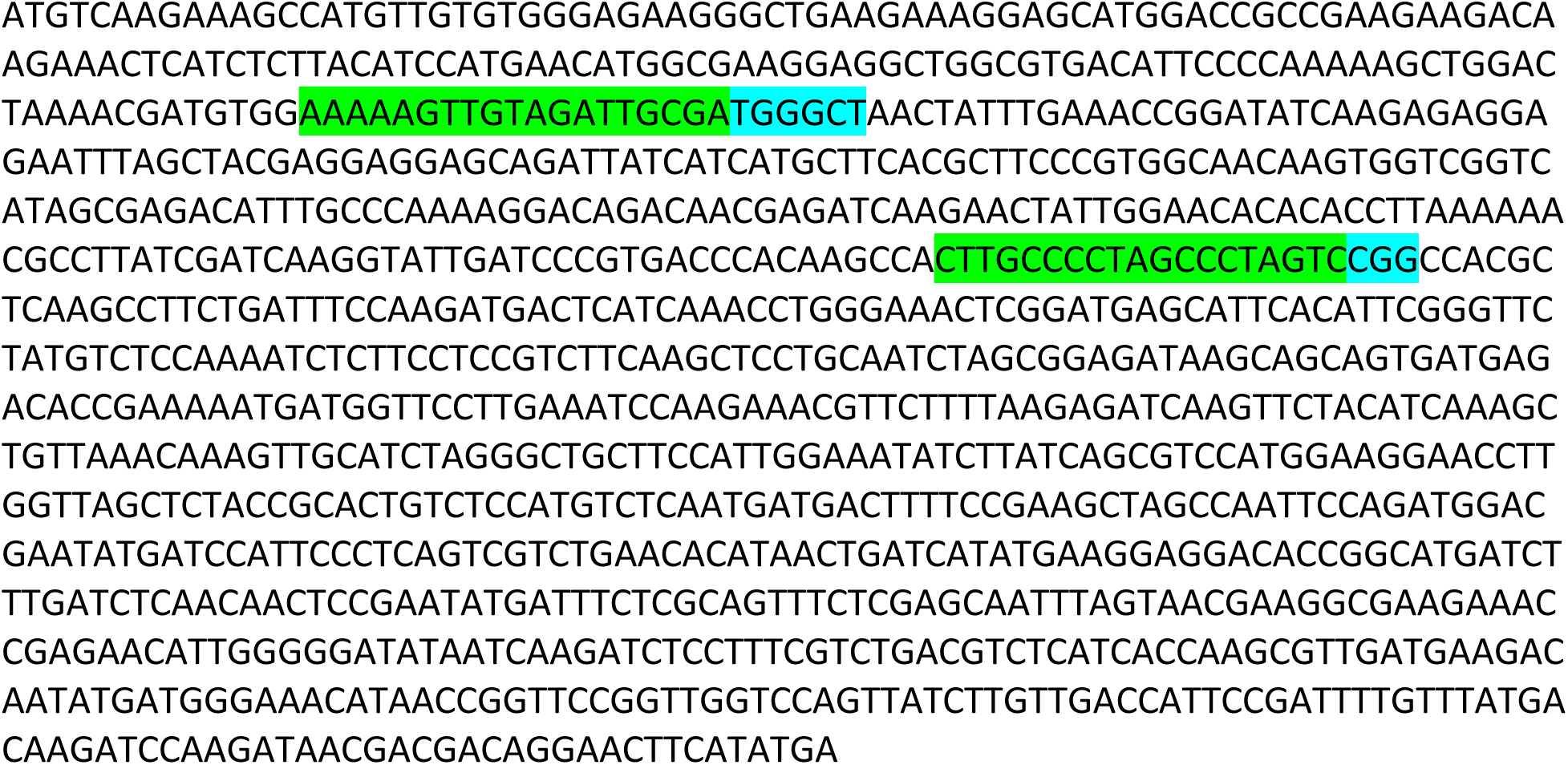

#### >Wild-Type *HAG3* amino acid sequence

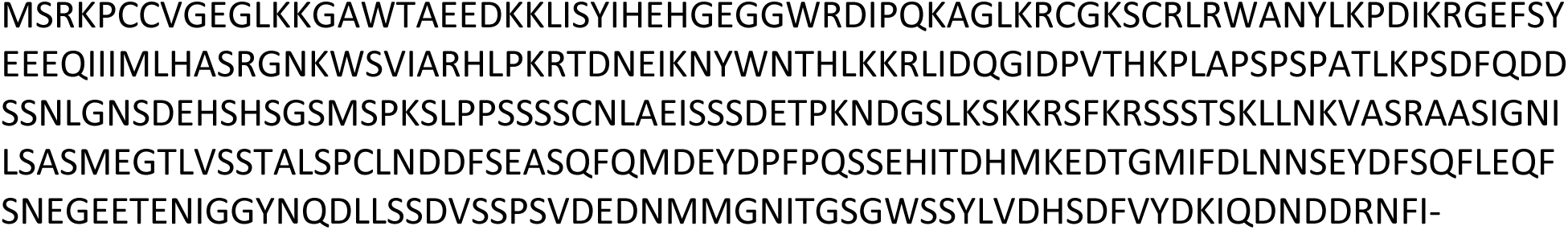

#### >*hag3-1* (−A) (red bold strikethrough)

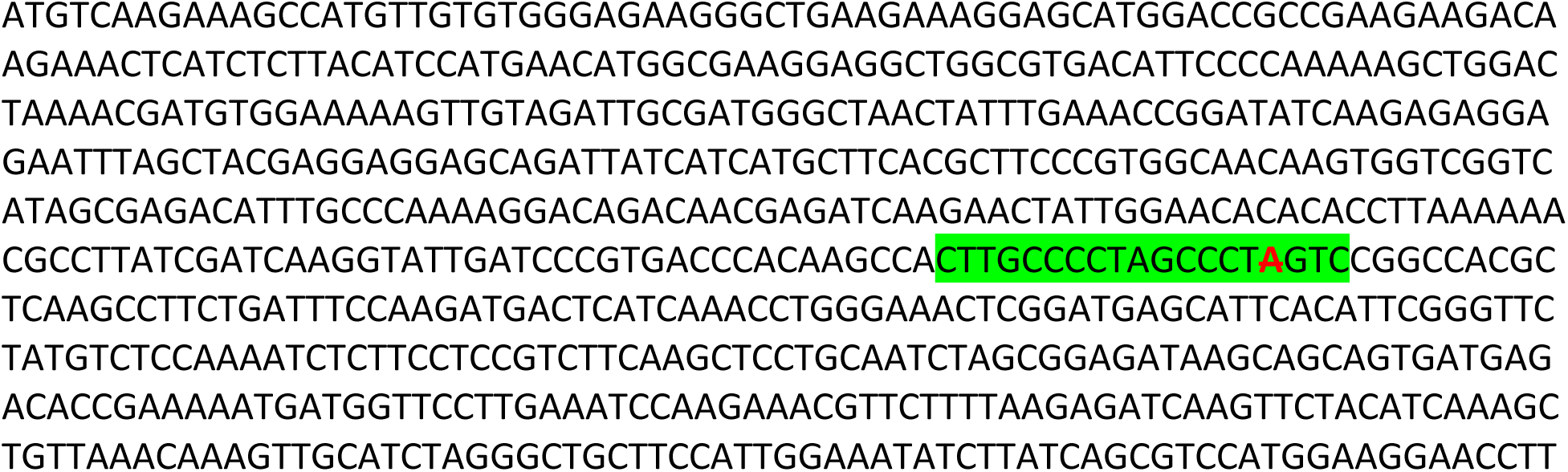
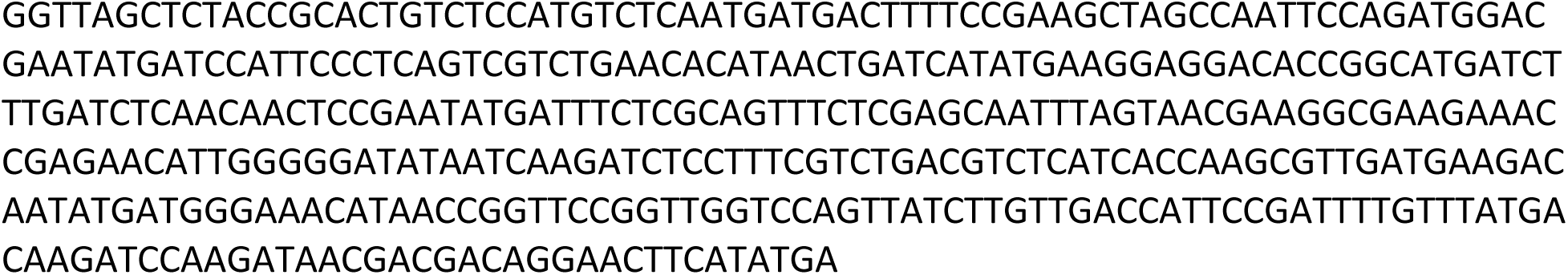

#### >*hag3-1* (−A) predicted amino acid sequence

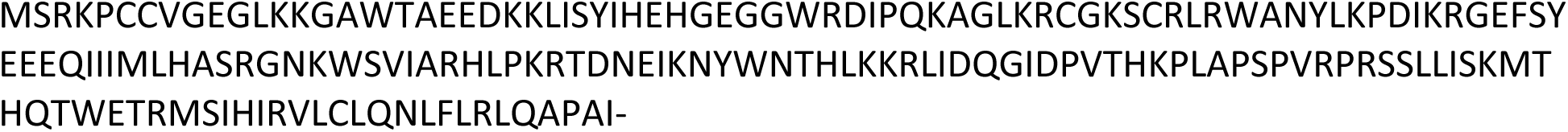

#### >hag*3* (−24bp) (red bold strikethrough)

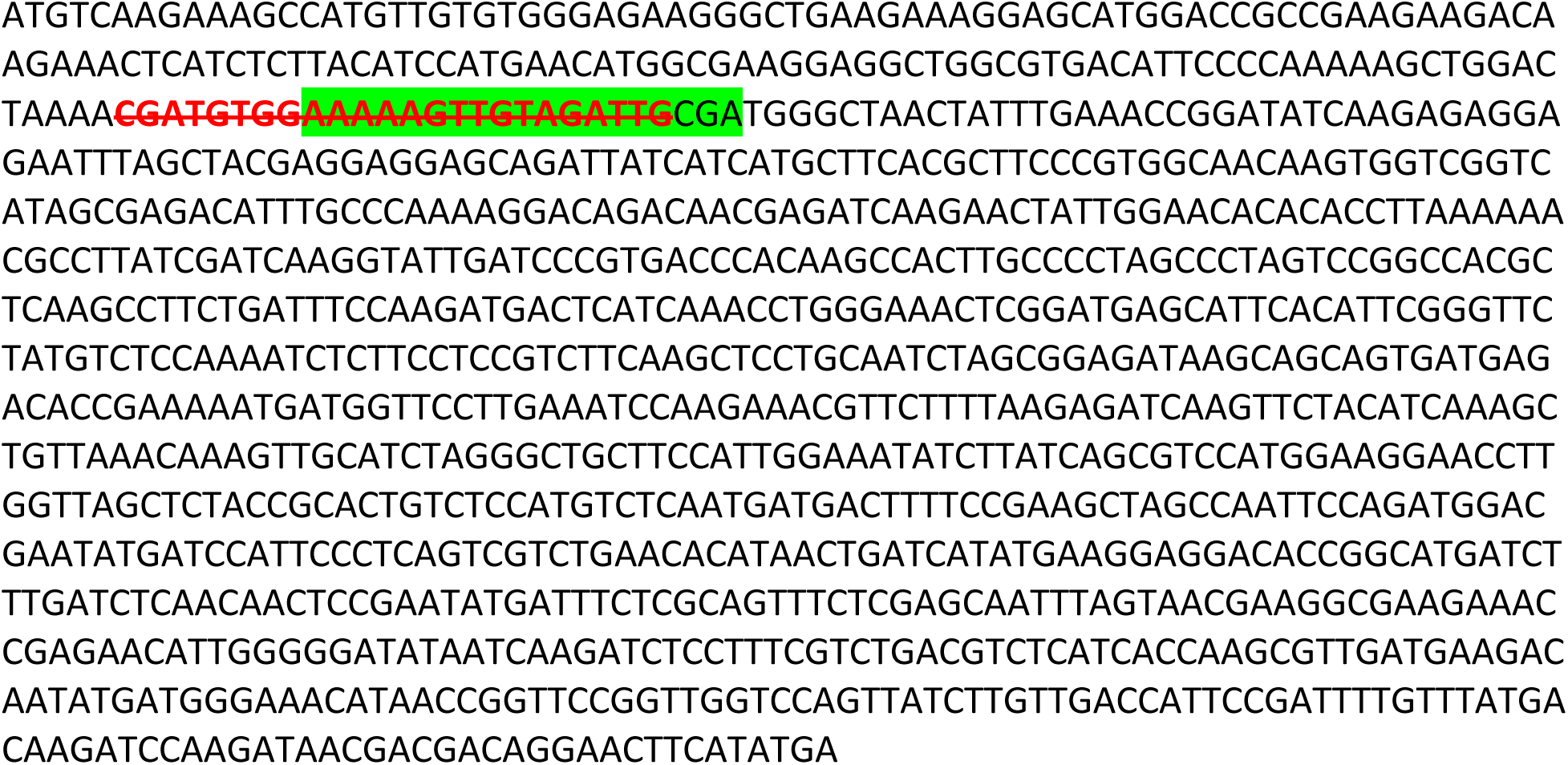

#### >*hag3* (−24bp) predicted amino acid sequence

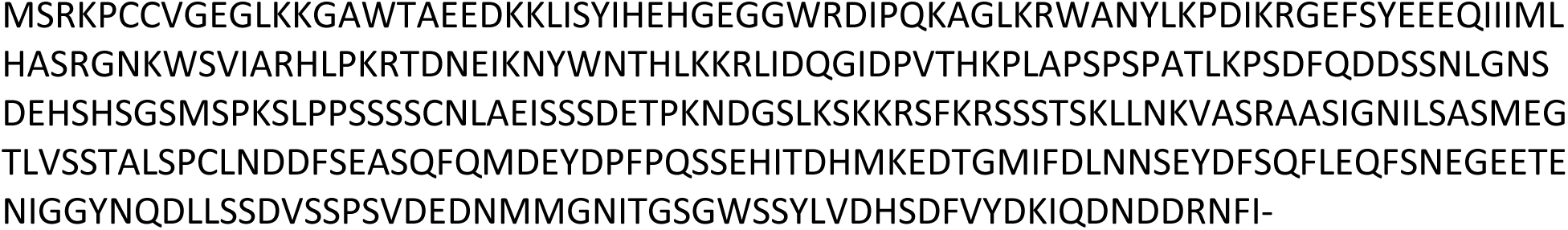

#### >*hag3* (−13bp) (red bold strikethrough) in *hag1hag3* (+A; −13bp) double mutant

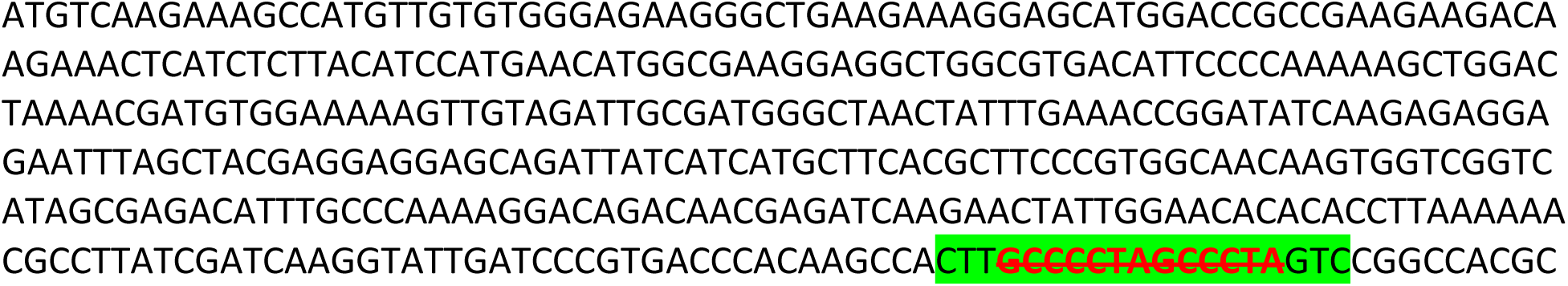
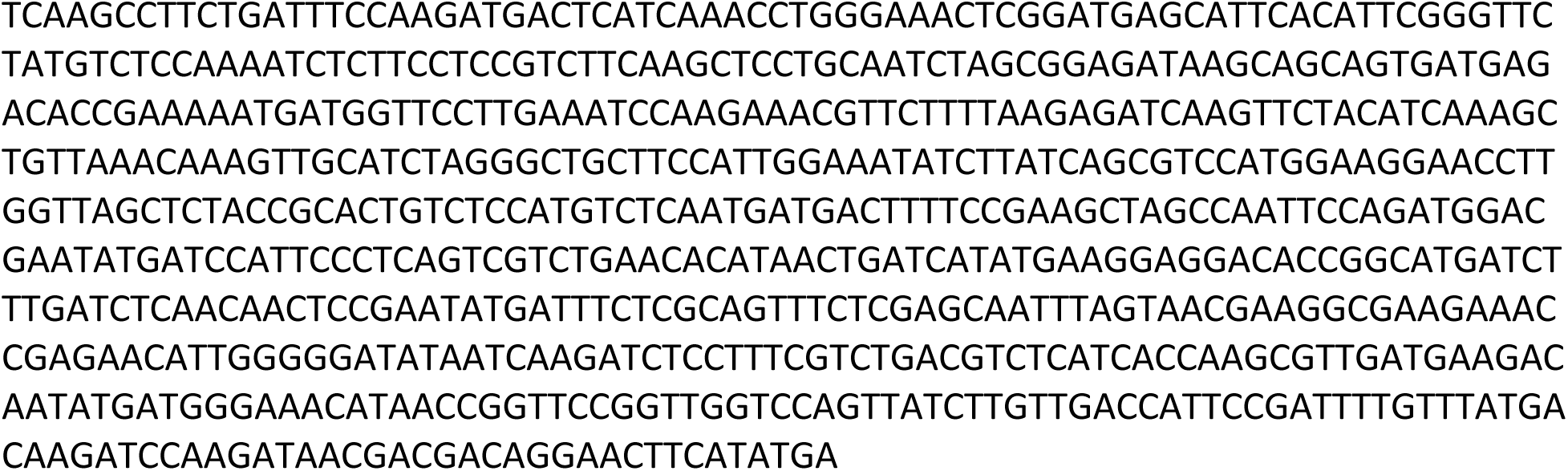

#### >*hag3* (−13bp) predicted amino acid sequence in *hag1hag3* (+A; −13bp) double mutant

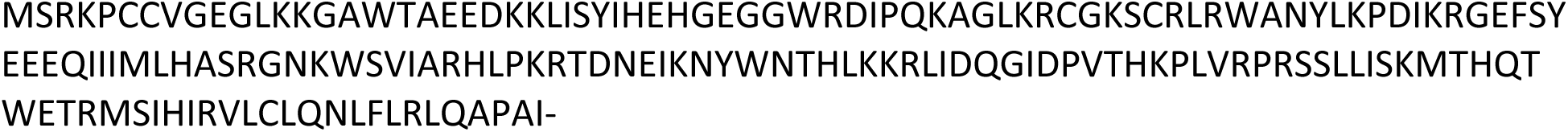

#### >*hag3* (−36bp) (red bold strikethrough) in *hag1hag2hag3* (+G; −9bp; −36bp) triple mutant

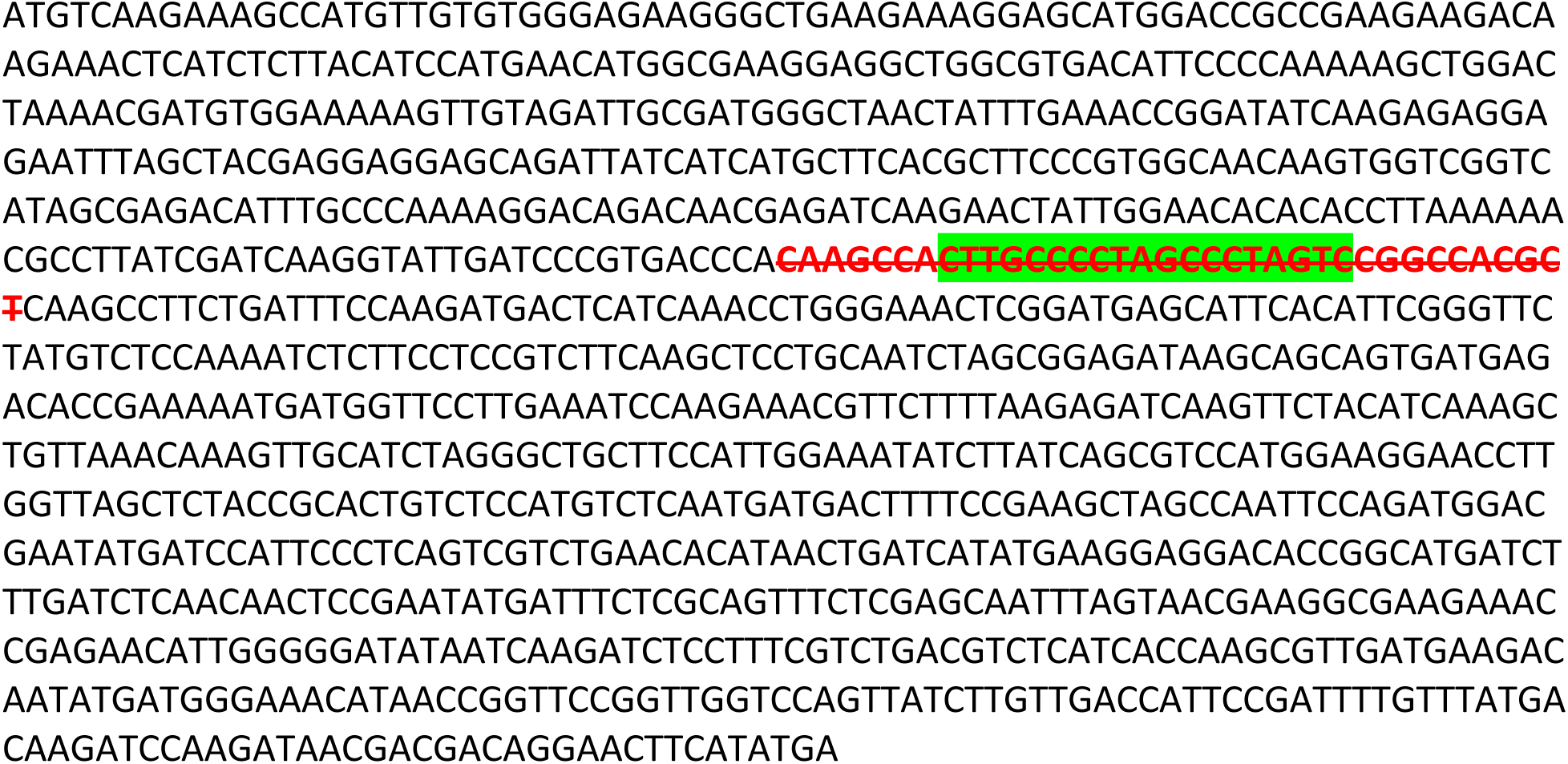

#### >*hag3* (−36bp) predicted amino acid sequence in *hag1hag2hag3* (+G; −9bp; −36bp) triple mutant

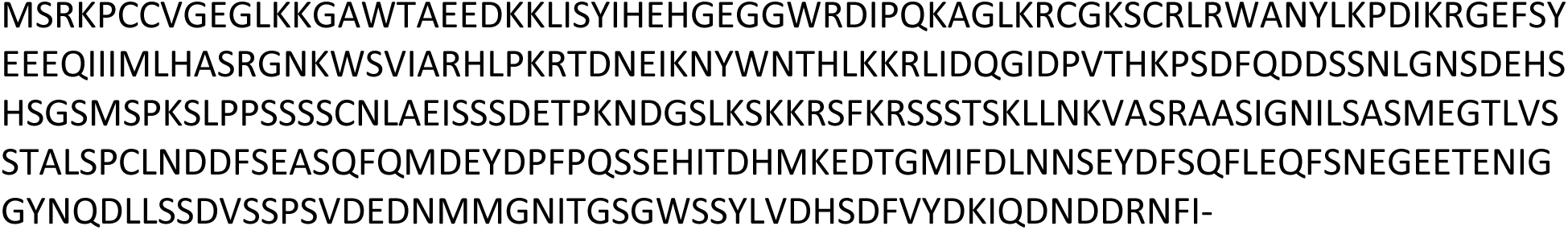

#### >*hag3* (+C) (bold red underlined) in *hag1hag2hag3* (−8bp; −3bp; +C) triple mutant

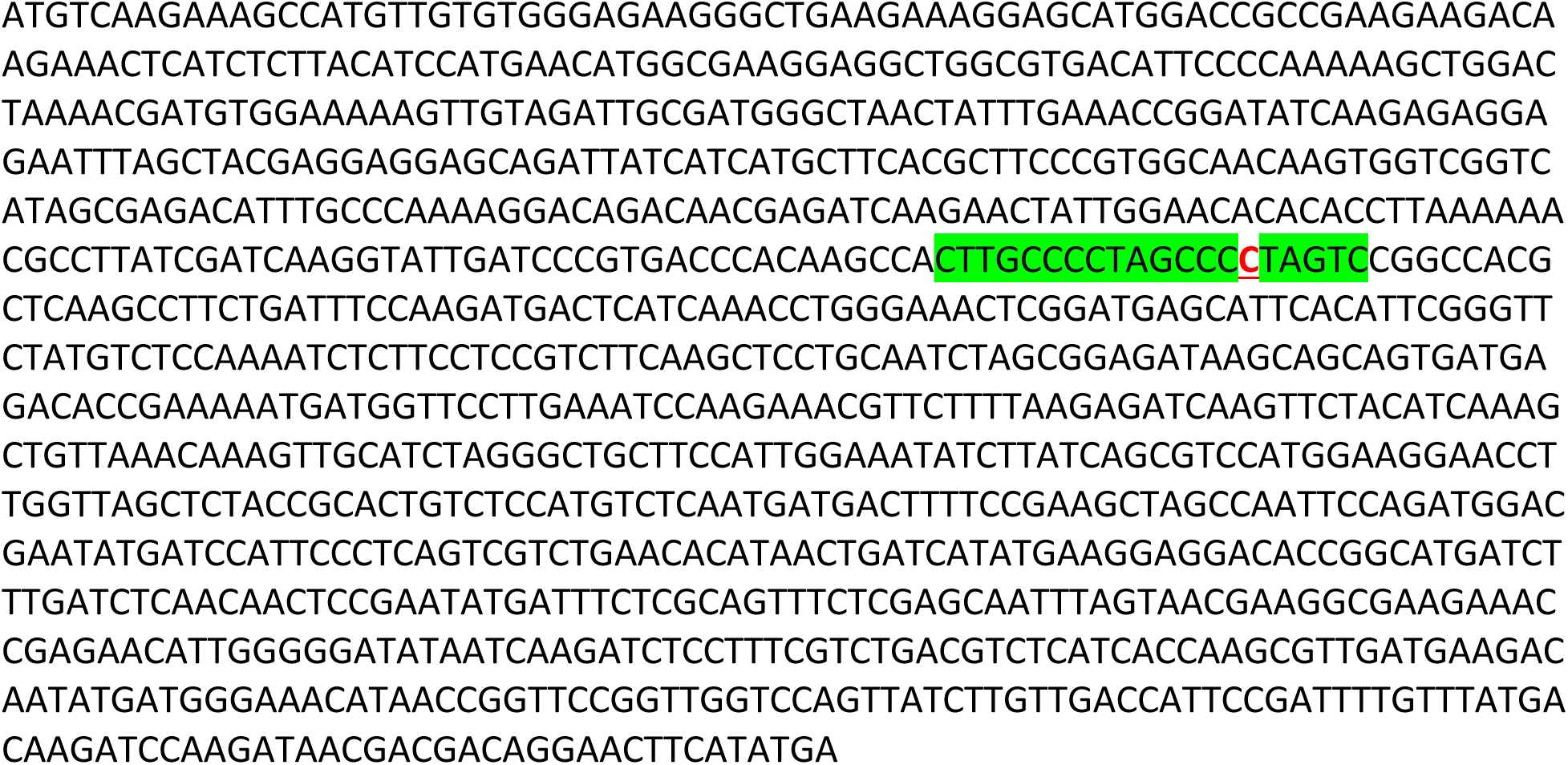

#### >*hag3* (+C) predicted amino acid sequence in *hag1hag2hag3* (−8bp; −3bp; +C) triple mutant

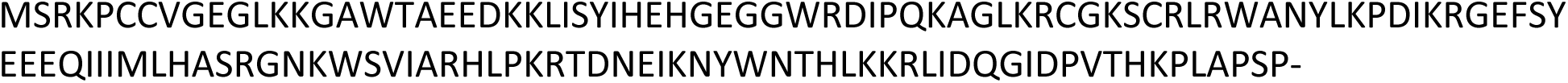

#### >Wild-Type *TT8* ORF sequence

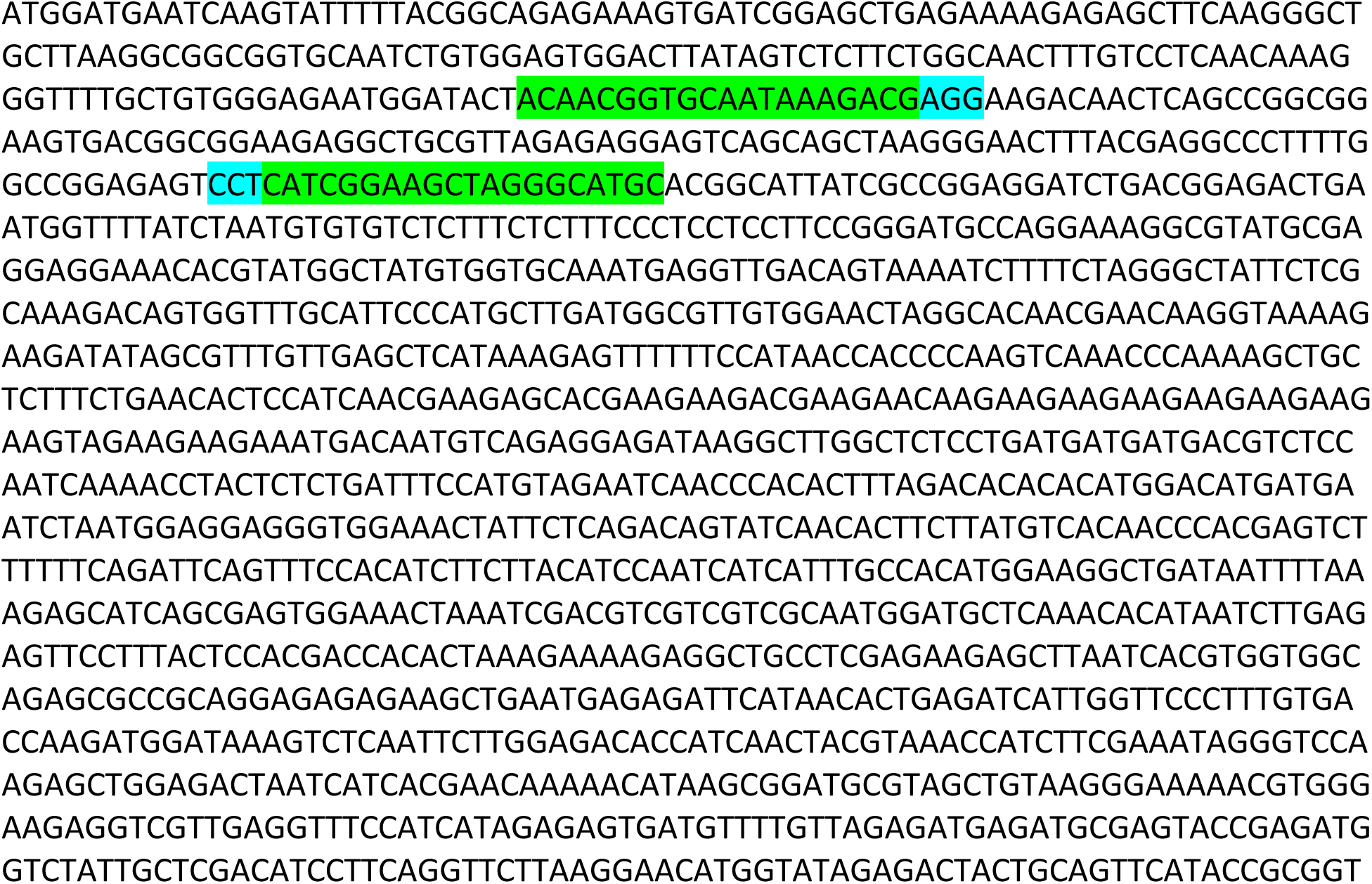
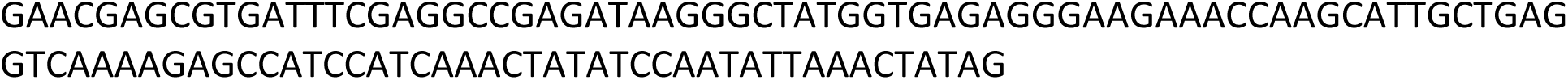

#### >Wild-Type *TT8* amino acid sequence

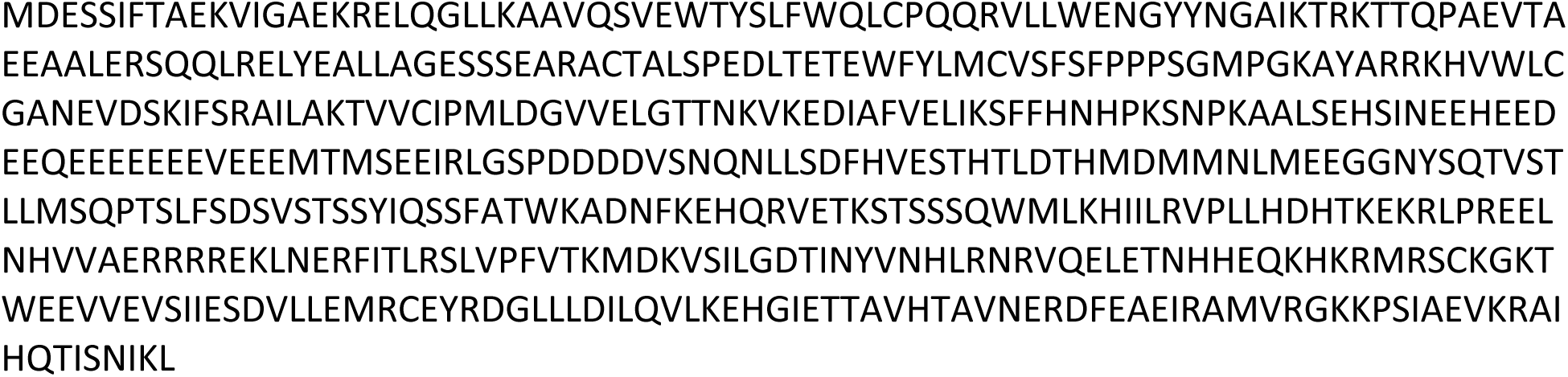

#### >*tt8* (−2bp) (red bold strikethrough)

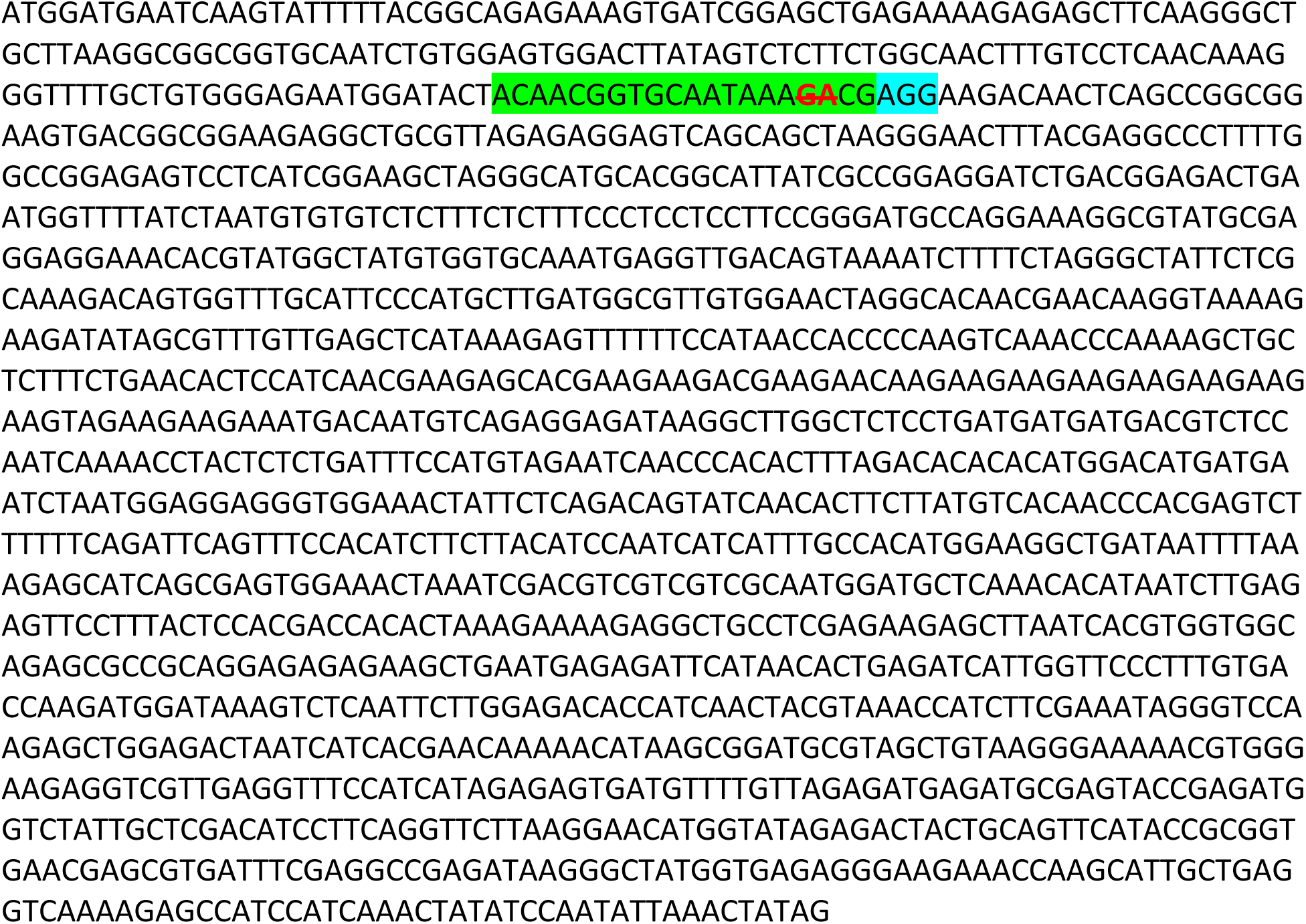

#### >*tt8* (−2bp) predicted amino acid sequence

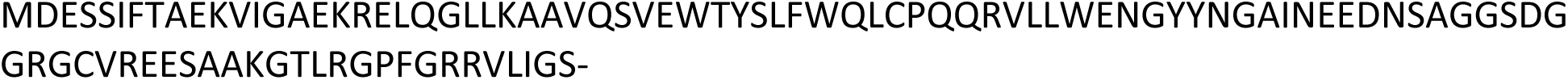

#### >*tt8* (+A) (red bold underlined)

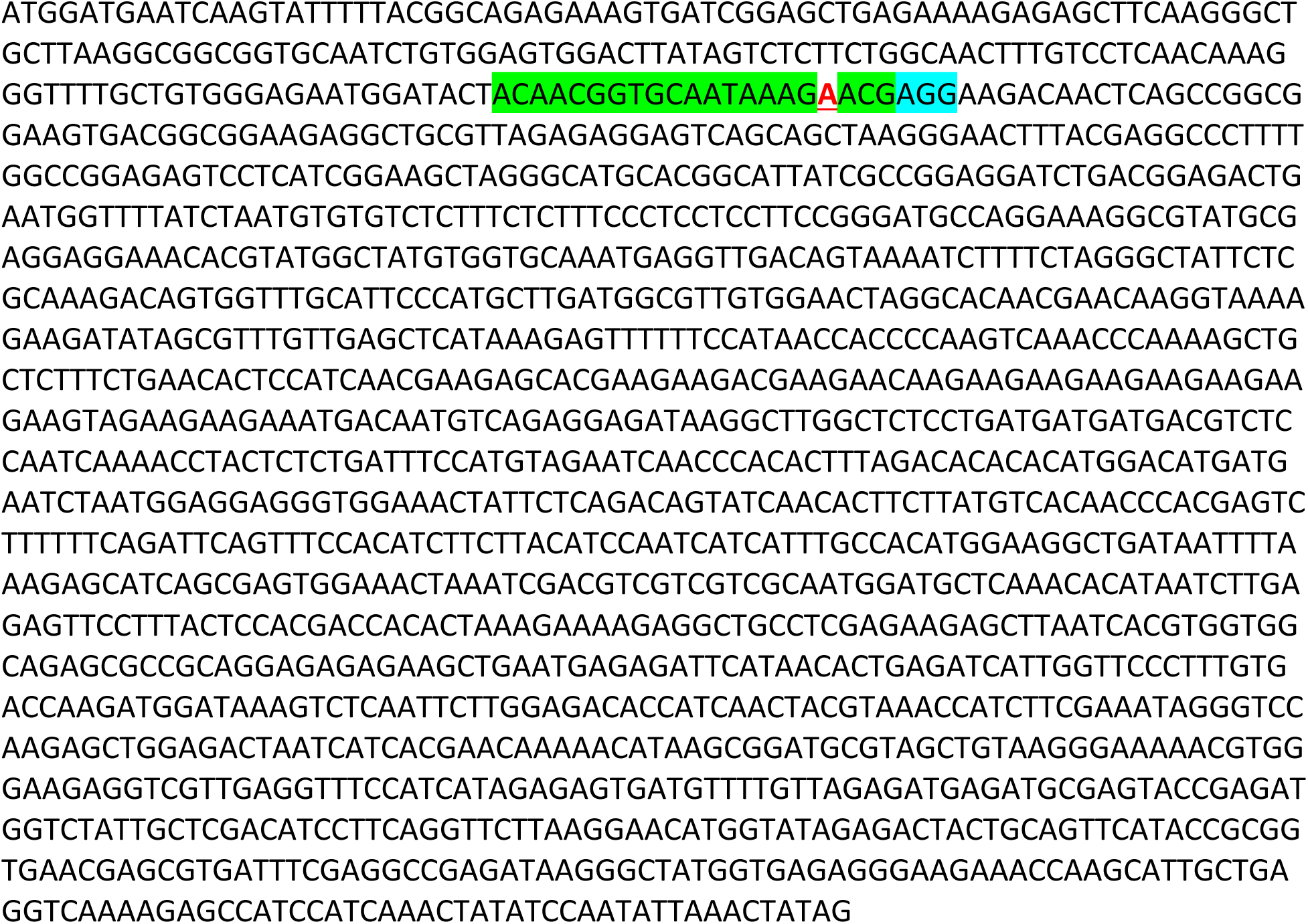

#### >*tt8* (+A) predicted protein sequence

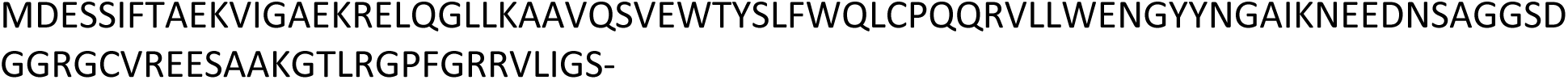

#### >*tt8* (−8bp) (red bold strikethrough)

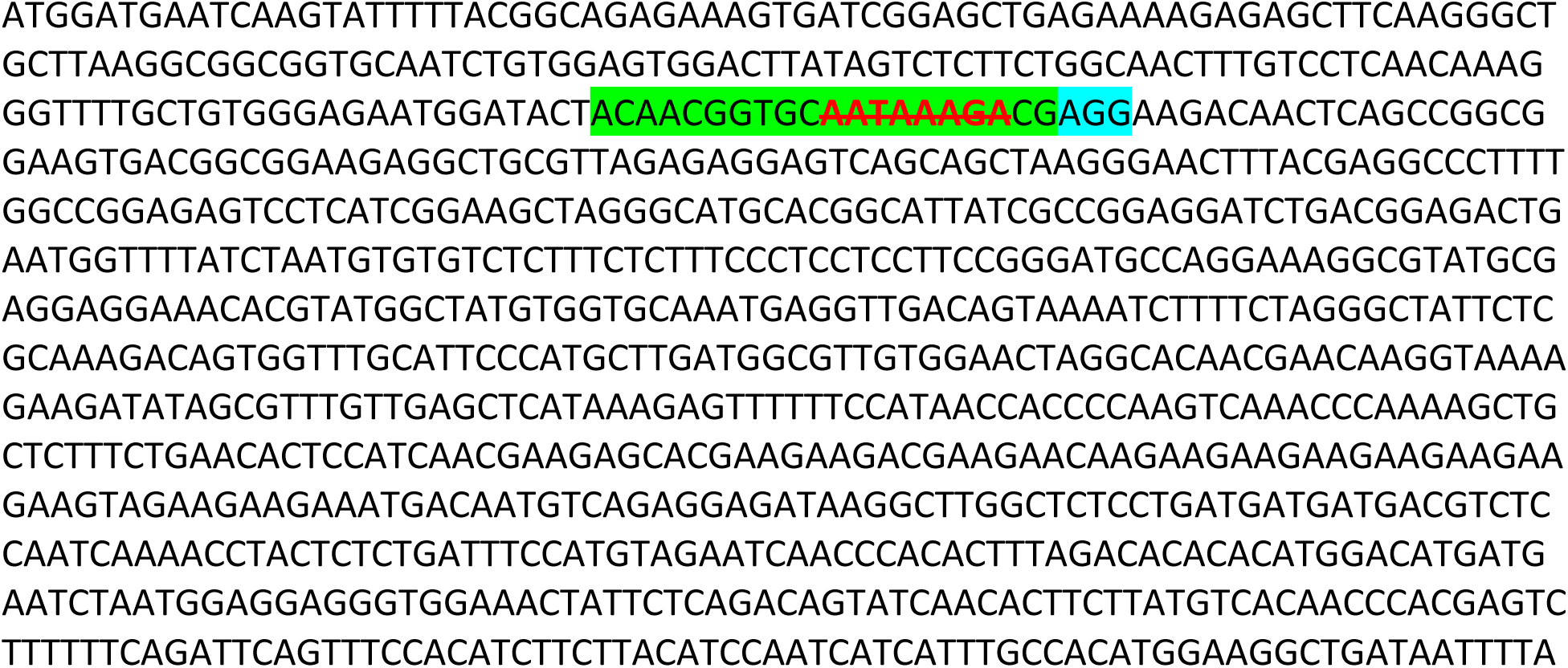
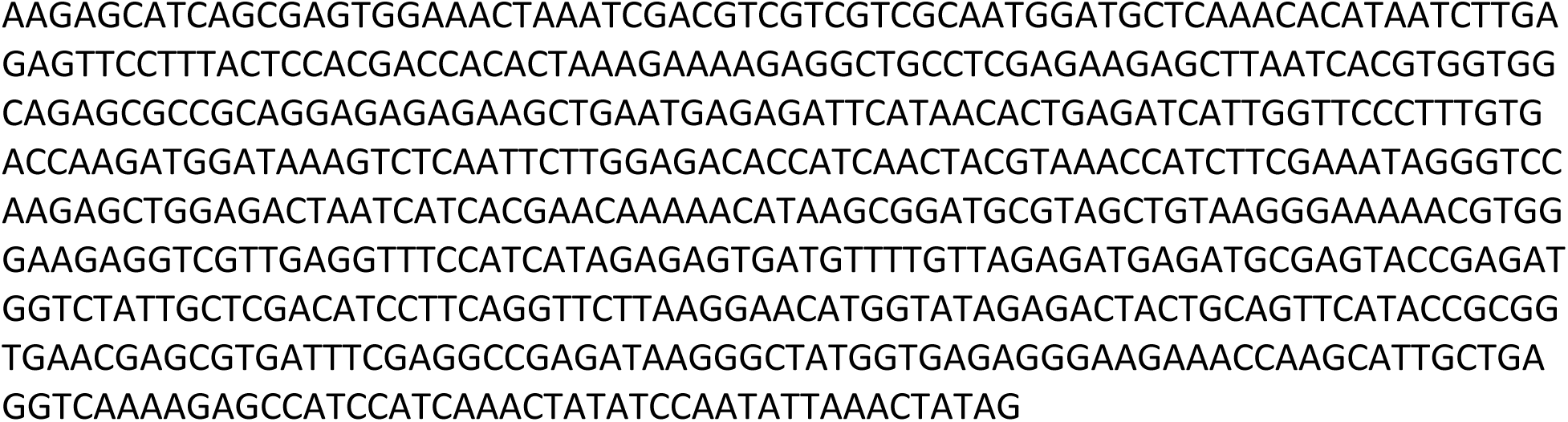

#### >*tt8* (−8bp) predicted amino acid sequence

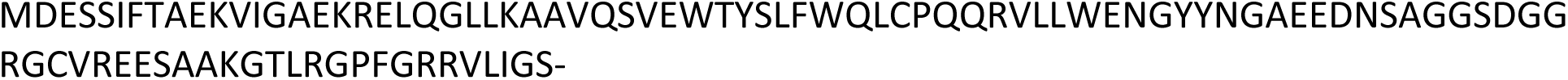

#### *>myc3hag3tt8fae1* (Bold orange: inversion of 595bp between PS1 and PS2 in HAG3, stacked in *(myc3tt8fae1)−1*)

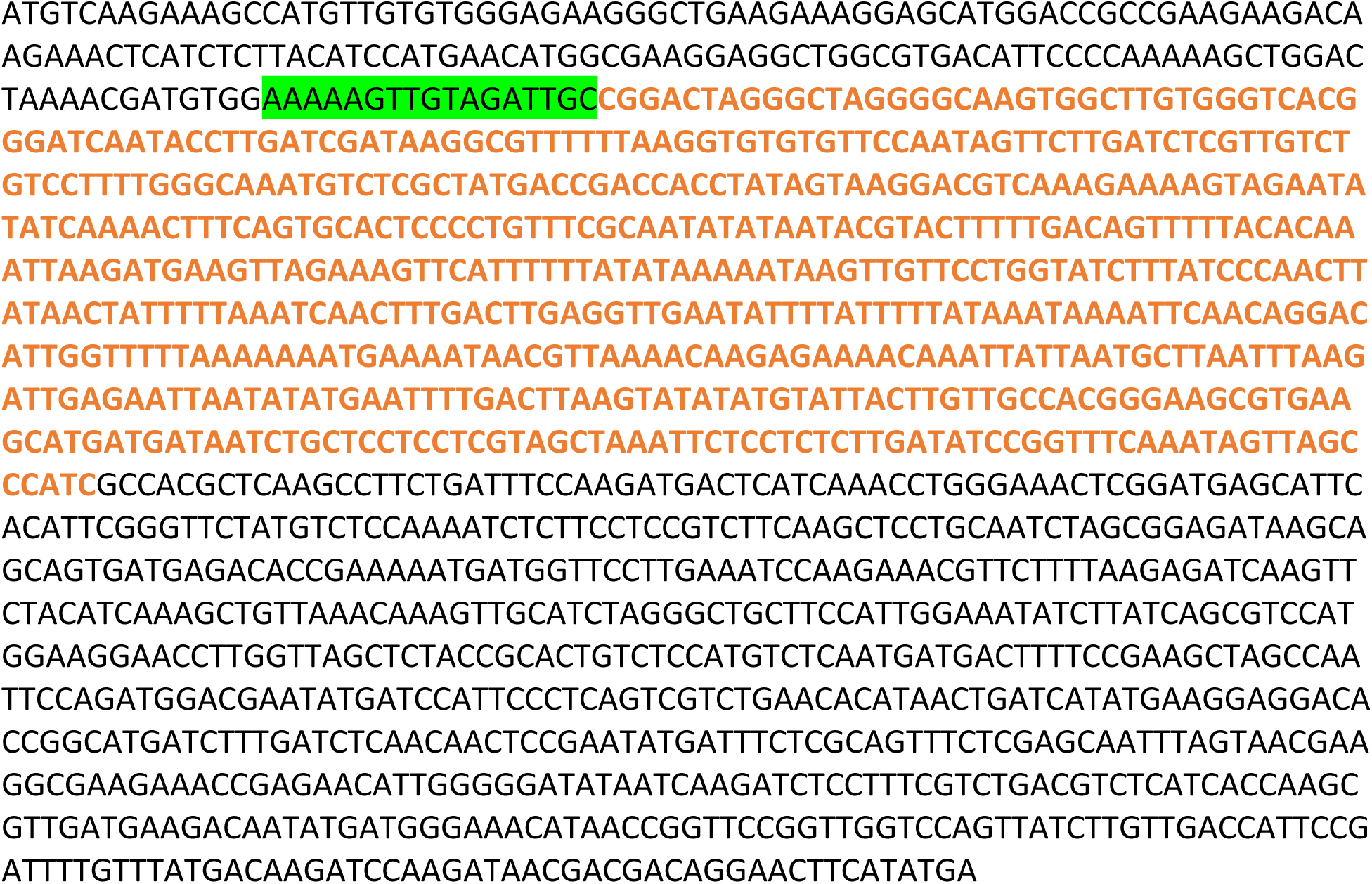

#### >myc3hag3tt8fae1 predicted amino acid sequence

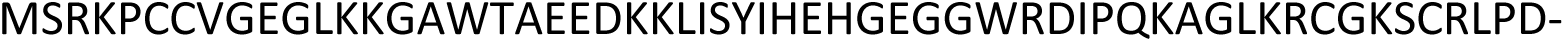

#### *>myc3hag1tt8fae1* (−6bp in PS1, -G in PS2 in HAG1, stacked in (*myc3tt8fae1)−1*) (red bold strikethrough)

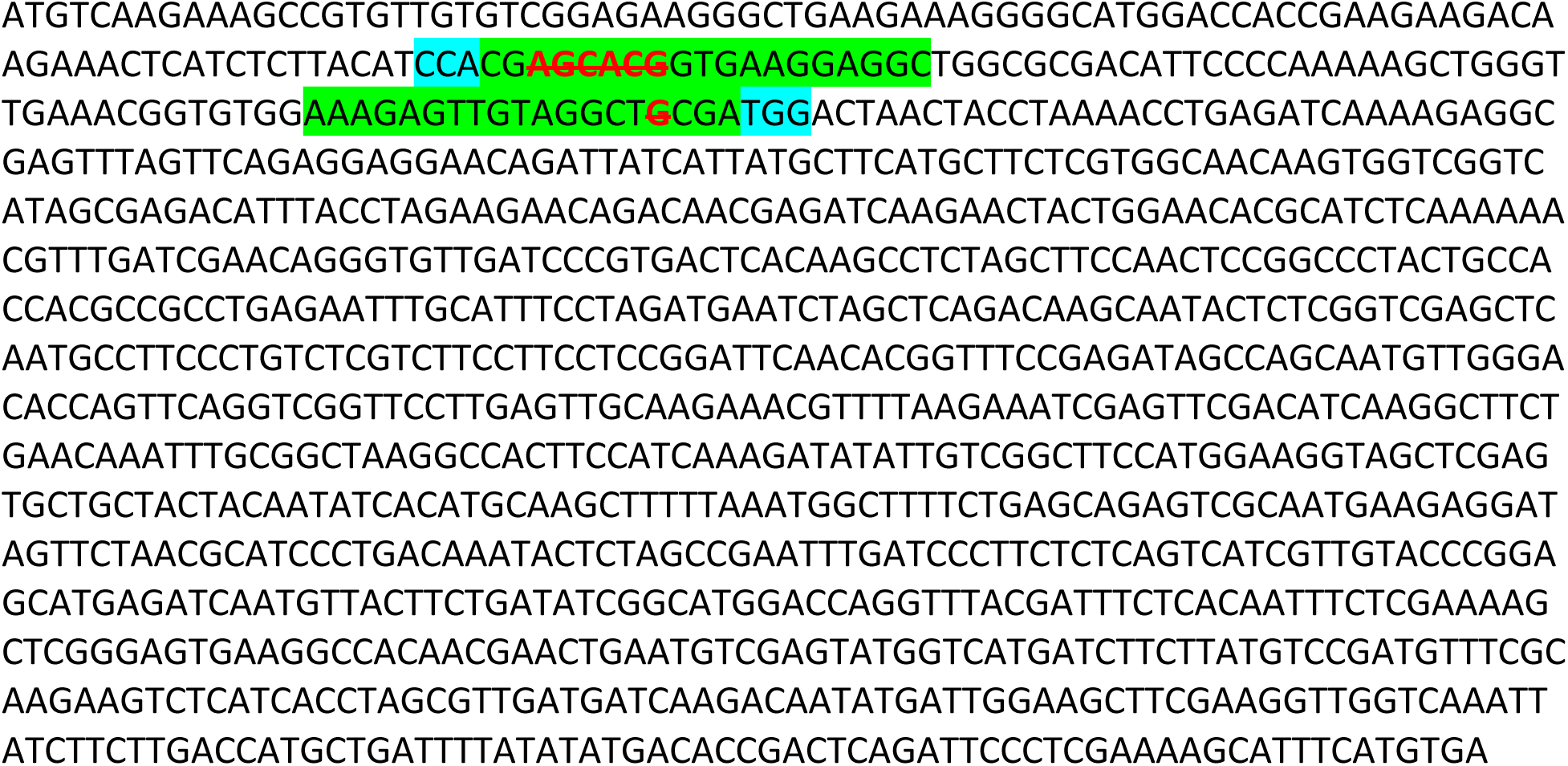

#### >myc3hag1tt8fae1 predicted amino acid sequence

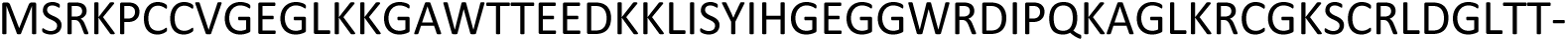

## REFERENCES

1. C. A. Eberle et al., Using pennycress, camelina, and canola cash cover crops to provision pollinators. Ind. Crops Prod. 75, 20–25 (2018). 10.1016/j.indcrop.2015.06.026

2. A. Plastina, F. Liu, F. Miguez, S. Carlson, Cover crops use in Midwestern US agriculture: perceived benefits and net returns. Renew. Agric. Food Syst. 36, 38–48 (2020). 10.1017/S1742170518000194

3. L. L. Van Eerd, I. Chahal, Y. Peng, J. C. Awrey, Influence of cover crops at the four spheres: A review of ecosystem services, potential barriers, and future directions for North America. Sci. Total Environ. 858, 159990 (2023). 10.1016/j.scitotenv.2022.159990

4. I. Vendig, A. Guzman, G. De La Cerda et al., Quantifying direct yield benefits of soil carbon increases from cover cropping. Nat. Sustain. 6, 1125–1134 (2023). 10.1038/s41893-023-01131-7

5. A. Borchers, E. Truex-Powell, S. Wallander, C. Nickerson, Multicropping practices: Recent trends in double cropping. Econ. Inf. Bull. 125, USDA, Washington, DC (2014). https://ageconsearch.umn.edu/record/262122

6. E. J. Ranck, L. A. Holden, J. A. Dillon, C. A. Rotz, K. J. Soder, Economic and environmental effects of double cropping winter annuals and corn using the Integrated Farm System Model. J. Dairy Sci. 103, 3804–3815 (2020). 10.3168/jds.2019-17525

7. K. Waha et al., Multiple cropping systems of the world and the potential for increasing cropping intensity. Glob. Environ. Change 64, 102181 (2020). 10.1016/j.gloenvcha.2020.102131

8. Y. C. Lu, K. B. Watkins, J. R. Teasdale, A. A. Abdul-Baki, Cover crops in sustainable food production. Food Rev. Int. 16, 121–187 (2000). 10.1081/FRI-100100285

9. V. Quintarel et al., Cover crops for sustainable cropping systems: a review. Agriculture 12, 2076 (2022). 10.3390/agriculture12122076

10. C. Zulauf, G. Schnitkey, N. Paulson, J. Coppess, Cover Crops and Covered Cropland, 2022 US Census of Agriculture. Farmdoc Daily 14, 37 (2024). Department of Agricultural and Consumer Economics, University of Illinois at Urbana-Champaign, February 22, 2024.

11. J. S. Bergtold, S. Ramsey, L. Maddy, J. R. Williams, A review of economic considerations for cover crops as a conservation practice. Renew. Agric. Food Syst. 34, 662–676 (2019). 10.1017/S1742170517000278

12. L. Chabal, R. Vyn, D. Mayers, L. L. Van Eerd, Cumulative impact of cover crops on soil carbon sequestration and profitability in a temperate humid climate. Sci. Rep. 10, 13381 (2020). 10.1038/s41598-020-70224-6

13. R. G. Trischuk, B. S. Schilling, N. H. Low, G. R. Gray, L. V. Gusta, Cold acclimation, de-acclimation and re-acclimation of spring canola, winter canola and winter wheat: The role of carbohydrates, cold-induced stress proteins and vernalization. Environ. Exp. Bot. 106, 156–163 (2014). 10.1016/j.envexpbot.2014.02.013

14. M. K. Walia, M. S. Wells, J. Cubins, D. Wyse, R. D. Gardner, F. Forcella, R. Gesch, Winter camelina seed yield and quality responses to harvest time. Ind. Crops Prod. 124, 765–775 (2018). 10.1016/j.indcrop.2018.08.025

15. F. Zanetti, T. A. Isbell, R. W. Gesch, R. L. Evangelista, E. Alexopoulou, B. Moser, A. Monti, Turning a burden into an opportunity: Pennycress (*Thlaspi arvense* L.) a new oilseed crop for biofuel production. Biomass Bioenergy 130, 105354 (2019). 10.1016/j.biombioe.2019.105354

16. M. J. Mulvaney, R. Seepaul, I. M. Small, D. L. Wright, S. V. Paula-Moraes, C. Crozier, P. Cockson, B. Whipker, R. Leon, Frost damage of Carinata grown in the Southeastern US: SS-AGR-420/AG420, 5/2018. EDIS 2018 (3), Gainesville, FL (2018). 10.32473/edis-ag420-2018

17. R. D. S. N. Júnior, C. W. Fraisse, M. Bashyal, M. J. Mulvaney, R. Seepaul, M. A. Z. Karrei, J. E. Iboyi, D. Perondi, V. A. Cerbaro, K. J. Boote, Brassica carinata as an off-season crop in the southeastern USA: Determining optimum sowing dates based on climate risks and potential effects on summer crop yield. Agric. Syst. 196, 103344 (2022). 10.1016/j.agsy.2021.103344

18. K. F. Best, G. I. McIntyre, The biology of Canadian weeds. IX. Thlaspi arvense. Can. J. Plant Sci. 55, 279–292 (1975). 10.4141/cjps75-039

19. S. I. Warwick, A. Francis, D. J. Susko, The biology of Canadian weeds. 9. Thlaspi arvense L. (updated). Can. J. Plant Sci. 82, 803–823 (2002). 10.4141/P01-159

20. L. F. Marek, B. Bingaman, C. A. C. Gardner, T. Isbell, *Thlaspi arvense*, a potential biodiesel crop: Preliminary evaluation of the USDA germplasm collection. In 20th Annual Meeting Association for the Advancement of Industrial Crops Abstracts, Maricopa, AZ, pp. 7–11 (2008). College Station, TX: AAIC.

21. T. A. Isbell, US effort in the development of new crops (*Lesquerella, Pennycress, Coriander, and Cuphea*). Oléagineux Corps Gras Lipides 16, 205–210 (2009). 10.1051/ocl.2009.0269

22. B. R. Moser, G. Knothe, S. F. Vaughn, T. A. Isbell, Production and evaluation of biodiesel from field pennycress (*Thlaspi arvense* L.) oil. Energy Fuels 23, 4149–4155 (2009). 10.1021/ef900337g

23. W. B. Phippen, M. E. Phippen, Soybean seed yield and quality as a response to field pennycress residue. Crop Sci. 52, 2767–2773 (2012). 10.2135/cropsci2012.03.0192

24. G. A. Johnson, M. B. Kantar, K. J. Betts, D. L. Wyse, Field pennycress production and weed control in a double crop system with soybean in Minnesota. Agron. J. 107, 532–540 (2015). 10.2134/agronj14.0292

25. J. A. Cubins, M. S. Wells, K. Frels, M. A. Ott, F. Forcella, G. A. Johnson, M. K. Walia, R. L. Becker, R. W. Gesch, Management of pennycress as a winter annual cash cover crop. A review. Agron. Sustain. Dev. 39, 1–11 (2019). 10.1007/s13593-019-0592-0

26. A. G. Thomas, Weed survey system used in Saskatchewan for cereal and oilseed crops. Weed Sci. 33, 34–43 (1985). 10.1017/S0043174500083892

27. G. F. Hartnell, S. Lemke, D. Moore, A. Matthews, M. A. Nemeth, R. Brister, S. Liu, C. Aulbach, Performance and health of broiler chickens fed low erucic acid, lower fiber pennycress (CoverCressTM) grain. Poult. Sci. 102, 102432 (2023). 10.1016/j.psj.2022.102432

28. L. O. L. O. Heijkenskjöld, L. A. R. S. Ernster, Studies of the mode of action of erucic acid on heart metabolism. Acta Med. Scand. 198, 75–83 (1975). 10.1111/j.0954-6820.1975.tb06560.x

29. M. Ishinaga, J. Sato, Y. Kitagawa, E. Sugimoto, M. Kito, Perturbation of phospholipid metabolism by erucic acid in male Sprague-Dawley rat heart. J. Biochem. 92, 253–263 (1982). 10.1093/oxfordjournals.jbchem.a133921

30. A. Galanty, M. Grudzińska, W. Paździora, P. Paśko, Erucic acid—both sides of the story: a concise review on its beneficial and toxic properties. Molecules 28, 1924 (2023). 10.3390/molecules28041924

31. L. Rask, E. Andréasson, B. Ekbom, S. Eriksson, B. Pontoppidan, J. Meijer, Myrosinase: gene family evolution and herbivore defense in Brassicaceae. Plant Mol. Evol., 93–113 (2000). https://link.springer.com/chapter/10.1007/978-94-011-4221-2_5

32. S. F. Vaughn, T. A. Isbell, D. Weisleder, M. A. Berhow, Biofumigant compounds released by field pennycress (Thlaspi arvense) seedmeal. J. Chem. Ecol. 31, 167–177 (2005). 10.1007/s10886-005-0982-4

33. F. Schweizer, P. Fernández-Calvo, M. Zander, M. Diez-Diaz, S. Fonseca, G. Glauser, M. G. Lewsey, J. R. Ecker, R. Solano, P. Reymond, *Arabidopsis* basic helix-loop-helix transcription factors MYC2, MYC3, and MYC4 regulate glucosinolate biosynthesis, insect performance, and feeding behavior. Plant Cell 25, 3117–3132 (2013). 10.1105/tpc.113.115139

34. A. Mazumder, A. Dwivedi, J. Du Plessis, Sinigrin and its therapeutic benefits. Molecules 21, 416 (2016). 10.3390/molecules21040416

35. M. E. Daxenbichler, G. F. Spencer, D. G. Carlson, G. B. Rose, A. M. Brinker, R. G. Powell, Glucosinolate composition of seeds from 297 species of wild plants. Phytochemistry 30, 2623–2638 (1991). 10.1016/0031-9422(91)85112-D

36. E. Y. Y. Eileen, I. J. Pickering, G. N. George, R. C. Prince, *In situ* observation of the generation of isothiocyanates from sinigrin in horseradish and wasabi. Biochim. Biophys. Acta Gen. Subj. 1527, 156–160 (2001). 10.1016/S0304-4165(01)00161-1

37. X. Wu, H. Huang, H. Childs, Y. Wu, L. Yu, P. R. Pehrsson, Glucosinolates in Brassica vegetables: Characterization and factors that influence distribution, content, and intake. Annu. Rev. Food Sci. Technol. 12, 485–511 (2021). 10.1146/annurev-food-070620-025744

38. P. Czerniawski, M. Piślewska-Bednarek, A. Piasecka, K. Kułak, P. Bednarek, Loss of MYB34 transcription factor supports the backward evolution of indole glucosinolate biosynthesis in a subclade of the Camelineae tribe and releases the feedback loop in this pathway in *Arabidopsis*. Plant Cell Physiol. 64, 80–93 (2023). 10.1093/pcp/pcac142

39. R. Chopra, E. B. Johnson, R. Emenecker, E. B. Cahoon, J. Lyons, D. J. Kliebenstein, E. Daniels, K. M. Dorn, M. Esfahanian, N. Folstad, K. Frels, M. McGinn, M. Ott, C. Gallaher, K. Altendorf, A. Berroyer, B. Ismail, J. A. Anderson, D. L. Wyse, T. Ulmasov, J. C. Sedbrook, M. D. Marks, Identification and stacking of crucial traits required for the domestication of pennycress. Nat. Food 1, 84–91 (2020). 10.1038/s43016-019-0007-z

40. S. K. Gupta, A. Pratap, History, origin, and evolution. Adv. Bot. Res. 45, 1–20 (2007). 10.1016/S0065-2296(07)45001-7

41. B. R. Stefansson, Z. P. Kondra, Tower summer rape. Can. J. Plant Sci. 55, 343–344 (1975).

42. J. M. Bell, From rapeseed to canola: A brief history of research for superior meal and edible oil. Poult. Sci. 61, 613–622 (1982). 10.3382/ps.0610613

43. J. K. Daun, M. N. Eskin, D. Hickling, Eds., Canola: Chemistry, Production, Processing, and Utilization (Elsevier, 2015).

44. J. K. Daun, D. R. DeClercq, Glucosinolates in Canadian canola. Glob. Counc. Innov. Rapeseed Canola Bull. (2003).

45. I. E. Sønderby, F. Geu-Flores, B. A. Halkier, Biosynthesis of glucosinolates—gene discovery and beyond. Trends Plant Sci. 15, 283–290 (2010). https://www.cell.com/trends/plant-science/fulltext/S1360-1385(10)00031-2?mobileUi%3D0=&code=cell-site

46. S. Mitreiter, T. Gigolashvili, Regulation of glucosinolate biosynthesis. J. Exp. Bot. 72, 70–91 (2021). 10.1093/jxb/eraa479

47. T. Zhang, R. Liu, J. Zheng, Z. Wang, T. E. Gao, M. Qin, X. Hu, Y. Wang, S. Yang, T. Li, Insights into glucosinolate accumulation and metabolic pathways in *Isatis indigotica* Fort. BMC Plant Biol. 22, 78 (2022). 10.1186/s12870-022-03455-6

48. H. Qin, G. J. King, P. Borpatragohain, J. Zou, Developing multifunctional crops by engineering Brassicaceae glucosinolate pathways. Plant Commun. 4, 100565 (2023). 10.1016/j.xplc.2023.100565

49. V. Kittipol, Z. He, L. Wang, T. Doheny-Adams, S. Langer, I. Bancroft, Genetic architecture of glucosinolate variation in *Brassica napus*. J. Plant Physiol. 240, 152988 (2019). 10.1016/j.jplph.2019.06.001

50. R. Chopra, E. B. Johnson, E. Daniels, M. McGinn, K. M. Dorn, M. Esfahanian, N. Folstad, K. Amundson, K. Altendorf, K. Betts, K. Frels, J. A. Anderson, D. L. Wyse, J. C. Sedbrook, M. D. Marks, Translational genomics using *Arabidopsis* as a model enables the characterization of pennycress genes through forward and reverse genetics. Plant J. 96, 1093–1105 (2018). 10.1111/tpj.14147

51. M. McGinn, W. B. Phippen, R. Chopra, S. Bansal, B. A. Jarvis, M. E. Phippen, K. M. Dorn, M. Esfahanian, T. J. Nazarenus, E. B. Cahoon, T. P. Durrett, M. D. Marks, J. C. Sedbrook, Molecular tools enabling pennycress (*Thlaspi arvense*) as a model plant and oilseed cash cover crop. Plant Biotechnol. J. 17, 776–788 (2019). 10.1111/pbi.13014

52. M. Esfahanian, T. J. Nazarenus, M. M. Freund, G. McIntosh, W. B. Phippen, M. E. Phippen, T. P. Durrett, E. B. Cahoon, J. C. Sedbrook, Generating pennycress (*Thlaspi arvense*) seed triacylglycerols and acetyl-triacylglycerols containing medium-chain fatty acids. *Front*. Energy Res. 10, 620118 (2021). 10.3389/fenrg.2021.620118

53. B. A. Jarvis, T. B. Romsdahl, M. McGinn, T. J. Nazarenus, E. B. Cahoon, K. D. Chapman, J. C. Sedbrook, CRISPR/Cas9-induced *fad2* and *rod1* mutations stacked with *fae1* confer high oleic acid seed oil in pennycress (*Thlaspi arvense* L.). Front. Plant Sci. 12, 652319 (2021). 10.3389/fpls.2021.652319

54. K. M. Dorn, J. D. Fankhauser, D. L. Wyse, M. D. Marks, *De novo* assembly of the pennycress (*Thlaspi arvense*) transcriptome provides tools for the development of a winter cover crop and biodiesel feedstock. Plant J. 75, 1028–1038 (2013). 10.1111/tpj.12267

55. K. M. Dorn, J. D. Fankhauser, D. L. Wyse, M. D. Marks, A draft genome of field pennycress (*Thlaspi arvense*) provides tools for the domestication of a new winter biofuel crop. DNA Res. 22, 121–131 (2015). 10.1093/dnares/dsu045

56. A. Nunn, I. Rodríguez-Arévalo, Z. Tandukar, K. Frels, A. Contreras-Garrido, P. Carbonell-Bejerano, P. Zhang, D. Ramos Cruz, K. Jandrasits, C. Lanz, A. Brusa, Chromosome-level *Thlaspi arvense* genome provides new tools for translational research and for a newly domesticated cash cover crop of the cooler climates. Plant Biotechnol. J. 20, 944–963 (2022). 10.1111/pbi.13775

57. Z. P. Wang, H. L. Xing, L. Dong, H. Y. Zhang, C. Y. Han, X. C. Wang, Q. J. Chen, Egg cell-specific promoter-controlled CRISPR/Cas9 efficiently generates homozygous mutants for multiple target genes in *Arabidopsis* in a single generation. Genome Biol. 16, 144 (2015). 10.1186/s13059-015-0715-0

58. B. R. Moser, S. N. Shah, J. K. Winkler-Moser, S. F. Vaughn, R. L. Evangelista, Composition and physical properties of cress (*Lepidium sativum* L.) and field pennycress (*Thlaspi arvense* L.) oils. Ind. Crops Prod. 30, 199–205 (2009). 10.1016/j.indcrop.2009.03.007

59. R. Chopra, N. Folstad, J. Lyons, T. Ulmasov, C. Gallaher, L. Sullivan, A. McGovern, R. Mitacek, K. Frels, K. Altendorf, A. Killam, The adaptable use of *Brassica* NIRS calibration equations to identify pennycress variants to facilitate the rapid domestication of a new winter oilseed crop. Ind. Crops Prod. 128, 55–61 (2019). 10.1016/j.indcrop.2018.10.079

60. B. Bushnell, BBMap: A fast, accurate, splice-aware aligner. Lawrence Berkeley National Laboratory (2014). LBNL Report #: LBNL-7065E. Retrieved from https://escholarship.org/uc/item/1h3515gn

61. P. J. Cock, C. J. Fields, N. Goto, M. L. Heuer, P. M. Rice, The Sanger FASTQ file format for sequences with quality scores, and the Solexa/Illumina FASTQ variants. Nucleic Acids Res. 38, 1767–1771 (2010). 10.1093/nar/gkp1137

62. A. Dobin, C. A. Davis, F. Schlesinger, J. Drenkow, C. Zaleski, S. Jha, P. Batut, M. Chaisson, T. R. Gingeras, STAR: ultrafast universal RNA-seq aligner. Bioinformatics 29, 15–21 (2013). 10.1093/bioinformatics/bts635

63. P. Ewels, M. Magnusson, S. Lundin, M. Käller, MultiQC: summarize analysis results for multiple tools and samples in a single report. Bioinformatics 32, 3047–3048 (2016). 10.1093/bioinformatics/btw354

64. M. I. Love, W. Huber, S. Anders, Moderated estimation of fold change and dispersion for RNA-seq data with DESeq2. Genome Biol. 15, 550 (2014). 10.1186/s13059-014-0550-8

65. Y. Benjamini, Y. Hochberg, Controlling the false discovery rate: a practical and powerful approach to multiple testing. J. R. Stat. Soc. B 57, 289–300 (1995). 10.1111/j.2517-6161.1995.tb02031.x

66. A. Rasoul, C. R. Johnston, J. LaChance, J. C. Sedbrook, A. P. Alonso, Propelling sustainable energy: Multi-omics analysis of pennycress *FATTY ACID ELONGATION1* knockout for biofuel production. Plant Physiol. 197, kiae650 (2025). 10.1093/plphys/kiae650

67. L. W. Mitich, Field pennycress (*Thlaspi arvense* L.)—the stinkweed. Weed Technol. 10, 675–678 (1996). 10.1017/S0890037X00040604

68. M. Y. Hirai, M. Klein, Y. Fujikawa, M. Yano, D. B. Goodenowe, Y. Yamazaki, S. Kanaya, Y. Nakamura, M. Kitayama, H. Suzuki, N. Sakurai, Elucidation of gene-to-gene and metabolite- to-gene networks in *Arabidopsis* by integration of metabolomics and transcriptomics. J. Biol. Chem. 280, 25590–25595 (2005). 10.1074/jbc.M502332200

69. H. Nour-Eldin, T. Andersen, M. Burow, et al., NRT/PTR transporters are essential for translocation of glucosinolate defense compounds to seeds. Nature 488, 531–534 (2012). 10.1038/nature11285

70. D. Xu, N. C. H. Sanden, L. L. Hansen, Z. M. Belew, S. R. Madsen, L. Meyer, M. E. Jørgensen, P. Hunziker, D. Veres, C. Crocoll, A. Schulz, Export of defensive glucosinolates is key for their accumulation in seeds. Nature 617, 132–138 (2023). 10.1038/s41586-023-05969-x

71. N. C. H. Sanden, C. Kanstrup, C. Crocoll, A. Schulz, H. H. Nour-Eldin, B. A. Halkier, D. Xu, An UMAMIT-GTR transporter cascade controls glucosinolate seed loading in *Arabidopsis*. Nat. Plants 10, 172–179 (2024). 10.1038/s41477-023-01598-4

72. T. Gigolashvili, R. Yatusevich, B. Berger, C. Müller, U. I. Flügge, The R2R3-MYB transcription factor HAG1/MYB28 is a regulator of methionine-derived glucosinolate biosynthesis in *Arabidopsis thaliana*. Plant J. 51, 247–261 (2007). 10.1111/j.1365-313X.2007.03133.x

73. M. Y. Hirai, K. Sugiyama, Y. Sawada, T. Tohge, T. Obayashi, A. Suzuki, R. Araki, N. Sakurai, H. Suzuki, K. Aoki, H. Goda, Omics-based identification of *Arabidopsis* Myb transcription factors regulating aliphatic glucosinolate biosynthesis. Proc. Natl. Acad. Sci. U.S.A. 104, 6478–6483 (2007). 10.1073/pnas.0611629104

74. J. Beekwilder, W. van Leeuwen, N. M. van Dam, M. Bertossi, V. Grandi, L. Mizzi, et al., The impact of the absence of aliphatic glucosinolates on insect herbivory in *Arabidopsis*. PLoS ONE 3, e2068 (2008). 10.1371/journal.pone.0002068

75. A. L. Harper, M. Trick, J. Higgins, F. Fraser, L. Clissold, R. Wells, C. Hattori, P. Werner, I. Bancroft, Associative transcriptomics of traits in the polyploid crop species *Brassica napus*. Nat. Biotechnol. 30, 798–802 (2012). 10.1038/nbt.2302

76. S. Jhingan, H. J. Harloff, A. Abbadi, C. Welsch, M. Blümel, D. Tasdemir, C. Jung, Reduced glucosinolate content in oilseed rape (*Brassica napus* L.) by random mutagenesis of *BnMYB28* and *BnCYP79F1* genes. Sci. Rep. 13, 2344 (2023). 10.1038/s41598-023-28661-6

77. P. Chowdhury, M. Hasanuzzaman, K. Nahar, Glucosinolates and its role in mitigating abiotic and biotic stress in *Brassicaceae*. Plant Stress Physiol.—Perspect. Agric. 10, (2022). 10.5772/intechopen.102367

78. D. J. Kliebenstein, Is specialized metabolite regulation specialized? J. Exp. Bot. 74, 4942–4948 (2023). 10.1093/jxb/erad209

79. B. Li, A. Gaudinier, M. Tang, M. Taylor-Teeples, N. T. Nham, C. Ghaffari, D. S. Benson, M. Steinmann, J. A. Gray, S. M. Brady, D. J. Kliebenstein, Promoter-based integration in plant defense regulation. Plant Physiol. 166, 1803–1820 (2014). 10.1104/pp.114.248716

80. C. Gao, S. Qi, K. Liu, D. Li, C. Jin, Z. Li, G. Huang, J. Hai, M. Zhang, M. Chen, *MYC2, MYC3*, and *MYC4* function redundantly in seed storage protein accumulation in *Arabidopsis*. Plant Physiol. Biochem. 108, 63–70 (2016). 10.1016/j.plaphy.2016.07.004

81. S. Duan, C. Jin, D. Li, C. Gao, S. Qi, K. Liu, J. Hai, H. Ma, M. Chen, *MYB76* inhibits seed fatty acid accumulation in *Arabidopsis*. Front. Plant Sci. 8, 226 (2017). 10.3389/fpls.2017.00226

82. S. Karim, K. O. Holmström, A. Mandal, P. Dahl, S. Hohmann, G. Brader, E. T. Palva, M. Pirhonen, *AtPTR3*, a wound-induced peptide transporter needed for defense against virulent bacterial pathogens in *Arabidopsis*. Planta 225, 1431–1445 (2007). 10.1007/s00425-006-0451-5

83. C. Xie, X. Zhou, X. Deng, Y. Guo, *PKS5*, a SNF1-related kinase, interacts with and phosphorylates *NPR1*, and modulates expression of *WRKY38* and *WRKY62*. J. Genet. Genomics 37, 359–369 (2010). 10.1016/S1673-8527(09)60054-0

84. M. L. Falcone Ferreyra, J. Emiliani, E. J. Rodriguez, V. A. Campos-Bermudez, E. Grotewold, P. Casati, The identification of maize and *Arabidopsis* type I flavone synthases links flavones with hormones and biotic interactions. Plant Physiol. 169, 1090–1107 (2015). 10.1104/pp.15.00515

85. Y. Narusaka, T. Shinya, M. Narusaka, N. Motoyama, H. Shimada, K. Murakami, N. Shibuya, Presence of *LYM2*-dependent but *CERK1*-independent disease resistance in *Arabidopsis*. Plant Signal. Behav. 8, e25345 (2013). 10.4161/psb.25345

86. S. C. Jiang, N. L. Engle, Z. Z. Banday, N. M. Cecchini, H. W. Jung, T. J. Tschaplinski, J. T. Greenberg, *ALD1* accumulation in *Arabidopsis* epidermal plastids confers local and non-autonomous disease resistance. J. Exp. Bot. 72, 2710–2726 (2021). 10.1093/jxb/eraa609

87. A. Arnaiz, L. Talavera-Mateo, P. Gonzalez-Melendi, M. Martinez, I. Diaz, M. E. Santamaria, *Arabidopsis* Kunitz trypsin inhibitors in defense against spider mites. Front. Plant Sci. 9, 986 (2018). 10.3389/fpls.2018.00986

88. E. Contreras, M. Martinez, Comparative and evolutionary analysis of *Arabidopsis RIN4*-like/*NOI* proteins induced by herbivory. PLoS ONE 17, e0270791 (2022). 10.1371/journal.pone.0270791

89. I. Debeaujon, K. M. Leon-Kloosterziel, M. Koornneef, Influence of the testa on seed dormancy, germination, and longevity in *Arabidopsis*. Plant Physiol. 122, 403–414 (2000). 10.1104/pp.122.2.403

90. B. Auger, N. Marnet, V. Gautier, A. Maia-Grondard, F. Leprince, M. Renard, S. Guyot, N. Nesi, J. M. Routaboul, A detailed survey of seed coat flavonoids in developing seeds of *Brassica napus* L. J. Agric. Food Chem. 58, 6246–6256 (2010). 10.1021/jf903619v

91. R. A. Dixon, S. Sarnala, Proanthocyanidin biosynthesis—a matter of protection. Plant Physiol. 184, 579–591 (2020). 10.1104/pp.20.00973

92. M. A. Ott, G. Gardner, K. M. Rai, D. L. Wyse, M. D. Marks, R. Chopra, *TRANSPARENT TESTA 2* allele confers major reduction in pennycress (*Thlaspi arvense* L.) seed dormancy. Ind. Crops Prod. 174, 114216 (2021). 10.1016/j.indcrop.2021.114216

93. I. Appelhagen, K. Thiedig, N. Nordholt, N. Schmidt, G. Huep, M. Sagasser, B. Weisshaar, Update on *TRANSPARENT TESTA* mutants from *Arabidopsis thaliana*: Characterization of new alleles from an isogenic collection. Planta 240, 955–970 (2014). 10.1007/s00425-014-2088-0

94. M. Chen, L. Xuan, Z. Wang, L. Zhou, Z. Li, X. Du, E. Ali, G. Zhang, L. Jiang, *TRANSPARENT TESTA8* inhibits seed fatty acid accumulation by targeting several seed development regulators in *Arabidopsis*. Plant Physiol. 165, 905–916 (2014). 10.1104/pp.114.235507

95. M. Griffiths, B. Gautam, C. Lebow, K. Duncan, X. Ding, P. Handakumbura, J. C. Sedbrook, C. N. Topp, Evaluation of 3D seed structure and cellular traits *in situ* using X-ray microscopy. Sci. Rep. 15, 4532 (2025). 10.1038/s41598-025-88482-7

96. A. Rai, S. Umashankar, M. Rai, L. B. Kiat, J. A. S. Bing, S. Swarup, Coordinate regulation of metabolite glycosylation and stress hormone biosynthesis by *TT8* in *Arabidopsis*. Plant Physiol. 171, 2499–2515 (2016). 10.1104/pp.16.00421

